# False discovery rate control enables trustworthy deep-learning-based de novo peptide sequencing

**DOI:** 10.64898/2026.06.29.735174

**Authors:** Zhendong Liang, Chengxin Dai, Tianze Ling, Tingpeng Yang, Yun Yang, Yeye Leng, Linhai Xie, Yonghong He, Fuchu He, Yu Wang, Cheng Chang

**Affiliations:** Peng Cheng Laboratory, Shenzhen, 518055, China; Tsinghua Shenzhen International Graduate School, Shenzhen, 518055, China; State Key Laboratory of Medical Proteomics, National Center for Protein Sciences (Beijing), Research Unit of Proteomics-driven Cancer Precision Medicine (Chinese Academy of Medical Sciences), Beijing 102206, China; International Academy of Phronesis Medicine (Guangdong), 510320, Guangdong, China; School of Life Sciences, Tsinghua University, Beijing, 100084, China

## Abstract

Deep-learning-based de novo peptide sequencing identifies peptides beyond predefined databases, but confidence thresholds do not consistently indicate false discovery rates (FDRs). We develop Counterpart Calibration Theory (CCT), which uses counterpart counts to estimate incorrect target predictions and defines conditions for conservative FDR estimation and threshold selection. We implement CCT in π-NovoQC at the peptide-spectrum match and peptide levels, integrating it with π-xNovo sequencing models, protein inference and quantification. Evaluations across sequencing models and datasets examine FDR estimation and correct-identification retention. Isotope-labelled data provide complementary evidence for identifications recovered from database-unidentified spectra. In the Deep Human Proteome dataset, the workflow recovers low-abundance peptide evidence that extends protein-group inference and quantification. These results support the incorporation of de novo identifications into proteomic analyses at specified FDR thresholds.

## Introduction

Deep-learning-based de novo peptide sequencing predicts peptide sequences from tandem mass spectra, enabling identification beyond predefined databases. This is useful when reference databases are incomplete or peptides contain sequence variants, unexpected modifications or poorly annotated protein products [1, 2]. Sequencing models commonly derive confidence scores from amino acid predictions: PepNet [3] uses their product, whereas Casanovo [4] uses their mean. Such scores rank predictions within a dataset, but a fixed threshold need not retain the same proportion of incorrect predictions across models, datasets or acquisition conditions. Reliable false discovery rate (FDR) estimation is therefore needed for selection and comparison under a common FDR criterion.

Recent models have improved sequencing across acquisition modes. For data-dependent acquisition (DDA), DeepNovo introduced stepwise decoding with convolutional and recurrent networks [5], while Casanovo formulated sequencing as Transformer-based sequence translation [4, 6]. π-HelixNovo, GraphNovo, π-PrimeNovo and InstaNovo use alternative architectures and decoding strategies [7, 8, 9, 10]. DeepNovo-DIA [11] and Cascadia [12] extend sequencing to chimeric data-independent acquisition (DIA) spectra. These advances improve candidate generation but do not establish a common relationship between model confidence and FDR.

In database search, target-decoy competition (TDC) estimates FDR using artificial decoy sequences as references for incorrect matches [13, 14]. With balanced search spaces and equal threshold-passing probabilities for incorrect target and decoy assignments, retained decoy counts estimate incorrect target counts. Dividing this estimated error count by the retained target count gives the FDR estimate. This construction is not directly transferable to de novo predictions (**Fig. 1a**). The sequencing model predicts each amino acid using spectral evidence and the context provided by other amino acids within the same peptide. This contextual dependence and the decoding procedure shape its errors; reversing, shuffling or replacing predicted residues does not ensure representative error sequences or scores. Because prediction correctness is unknown in routine applications, constructed references associated with correct and incorrect predictions cannot be separated in advance. An alternative is to count all constructed references passing a threshold. References associated with correct predictions contribute additional counts, which can produce a conservative FDR estimate when the total exceeds the number of retained incorrect predictions. The required scoring and cumulative-count conditions must therefore be established.

**Fig. 1.**
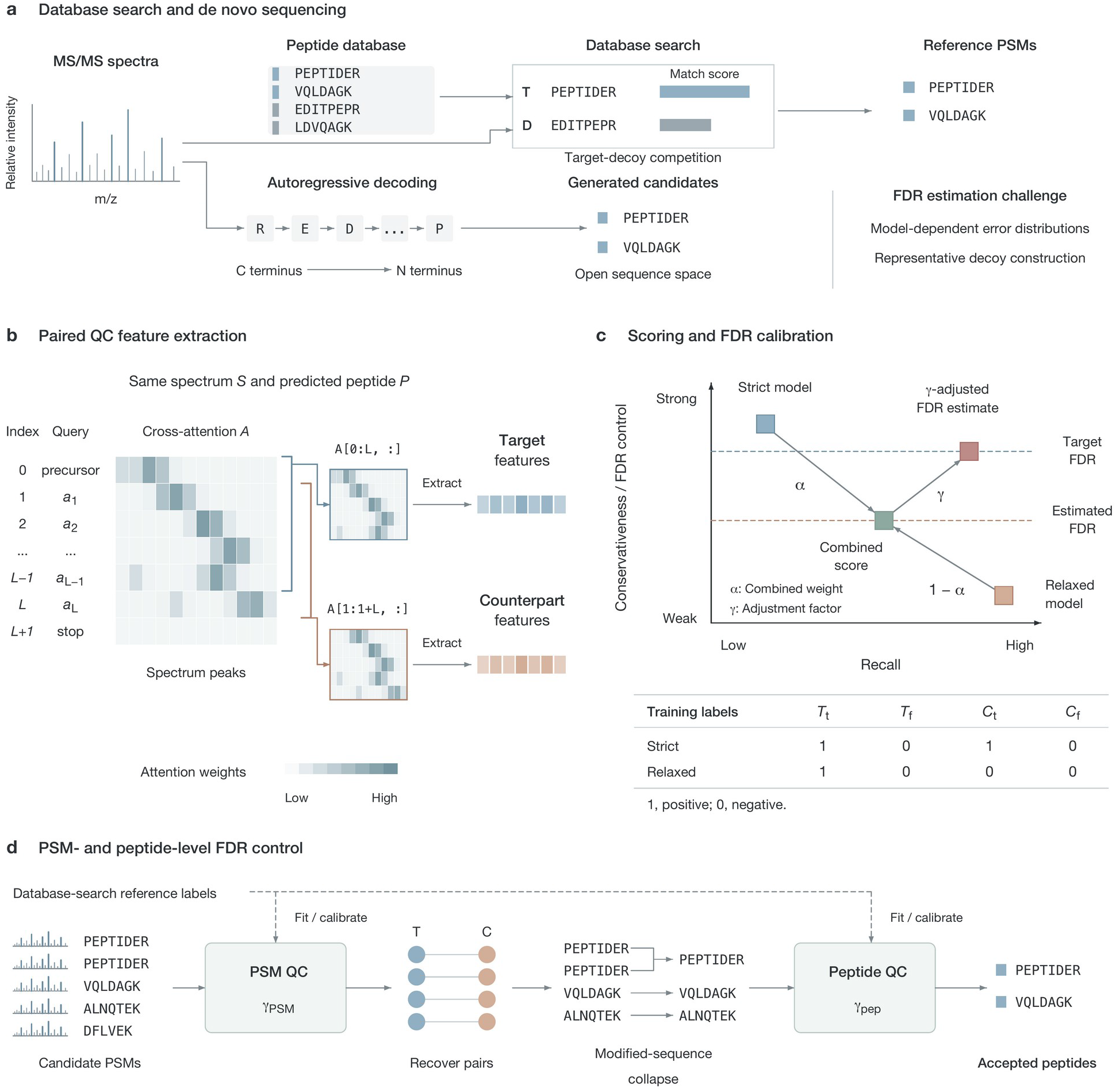
Conceptual framework of target-counterpart FDR control. a, Database-search target-decoy competition and model-generated de novo peptide predictions. b, Paired target and counterpart QC features extracted from the same spectrum and predicted peptide. c, Combination of strict and relaxed scoring followed by FDR calibration. T and C denote target and counterpart entries; subscripts t and f indicate agreement or disagreement with the database-search reference. d, Dual-level quality-control workflow.

Existing methods improve the reliability of de novo identifications in different ways. PostNovo [15] combines outputs from multiple de novo sequencers to rescore candidate sequences and support selection at a specified FDR. NovoBoard [16] constructs decoy spectra and uses model confidence scores for FDR estimation, while Winnow [17] combines model confidence scores with peptide-spectrum match (PSM) features to estimate calibrated probabilities of correctness and FDR. Glissade [18] integrates de novo predictions and database-search results for FDR control. Reliable FDR estimation and appropriate threshold selection are prerequisites for obtaining large sets of de novo identifications with controlled FDR and using them reliably in downstream proteomic analyses [19, 20, 21]. For the reference strategy outlined above, which we implement using counterparts, we investigate the scoring and cumulative-count conditions under which the number of references passing a score threshold can overestimate the number of incorrect predictions, thereby yielding conservative FDR estimates, and how this relationship can guide threshold selection at both the PSM and peptide levels.

We introduce Counterpart Calibration Theory (CCT), which formalizes the scoring and cumulative-count conditions for overestimating incorrect target counts and obtaining conservative FDR estimates, together with a corresponding threshold-selection procedure. Paired target and counterpart features from the same spectrum and predicted peptide are scored and ranked together. The retained counterpart count estimates the number of incorrect targets, and its ratio to retained targets estimates FDR. We implement CCT in π-NovoQC and develop DDA and DIA versions of π-xNovo to generate candidates and features. The workflow combines PSM- and peptide-level FDR estimation and control, peptide-to-protein mapping, separate target-decoy protein inference and MS1 quantification. Integrating accepted de novo identifications with database-search results extends peptide evidence from database-unidentified spectra into protein inference and quantitative analysis.

## Results

### 1. CCT framework for FDR estimation and control

CCT constructs paired target and counterpart QC features from each predicted peptide and its experimental spectrum, without generating a separate false peptide. Each feature vector represents an entry for scoring and counting: a candidate PSM at the PSM level or a unique modified sequence at the peptide level. Paired entries describe the same prediction, not independent identifications. After pooling and ranking the entries, CCT uses all counterparts passing a threshold to estimate incorrect targets. Counterparts associated with correct predictions contribute additional counts. When the complete counterpart count exceeds the incorrect-target count, its ratio to retained targets gives a conservative FDR estimate.

Using database-search identifications for the same spectra as references, target and counterpart entries are classified into four groups: target entries associated with correct predictions (*T_t_*), target entries associated with incorrect predictions (*T_f_*), counterpart entries associated with correct predictions (*C_t_*), and counterpart entries associated with incorrect predictions (*C_f_*). These group definitions allow score distributions and count-based FDR estimation and control to be evaluated on PSMs with database-search identifications. For a scoring function and threshold *s_i_*, let *T_t_* (*s_i_*), *T_f_* (*s_i_*), *C_t_* (*s_i_*) and *C_f_* (*s_i_*) denote the cumulative numbers of entries passing the threshold in the four groups. The total numbers of passing target and counterpart entries are *T* (*s_i_*)= *T_t_* (*s_i_*)+ *T_f_* (*s_i_*) and *C* (*s_i_*)= *C_t_* (*s_i_*)+ *C_f_* (*s_i_*), respectively. The observed false discovery proportion among accepted target predictions is

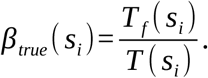

When it is unknown whether a prediction is correct, CCT uses *C* (*s_i_*), the cumulative count of all counterparts passing the same threshold, to estimate the unknown error count *T_f_* (*s_i_*). For thresholds with *T* (*s_i_*)> 0, the corresponding FDR estimate is

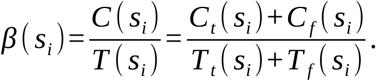

At a given threshold, *C* (*s_i_*)*≥T_f_* (*s_i_*) gives an estimate no smaller than the observed FDP. The score distributions determine which entries contribute to these cumulative counts and therefore where conservative estimation becomes possible. The assumptions and proofs underlying the scoring and count relationships are given in Supplementary Note 1.

To examine how scoring affects this relationship, we analyzed predictions from MassIVE-KB-trained [22] π-xNovo on the GraphNovo three-species test set [8], comprising *Arabidopsis thaliana*, *Caenorhabditis elegans* and *Escherichia coli*. After side-specific feature filtering, the analyzed population contained 37,321 target entries and 36,887 counterpart entries at the PSM level. The same population of PSM-level entries was used to compare relaxed, strict and combined scoring, with prediction correctness assessed against database-search identifications (**Fig. 2**).

**Fig. 2.**
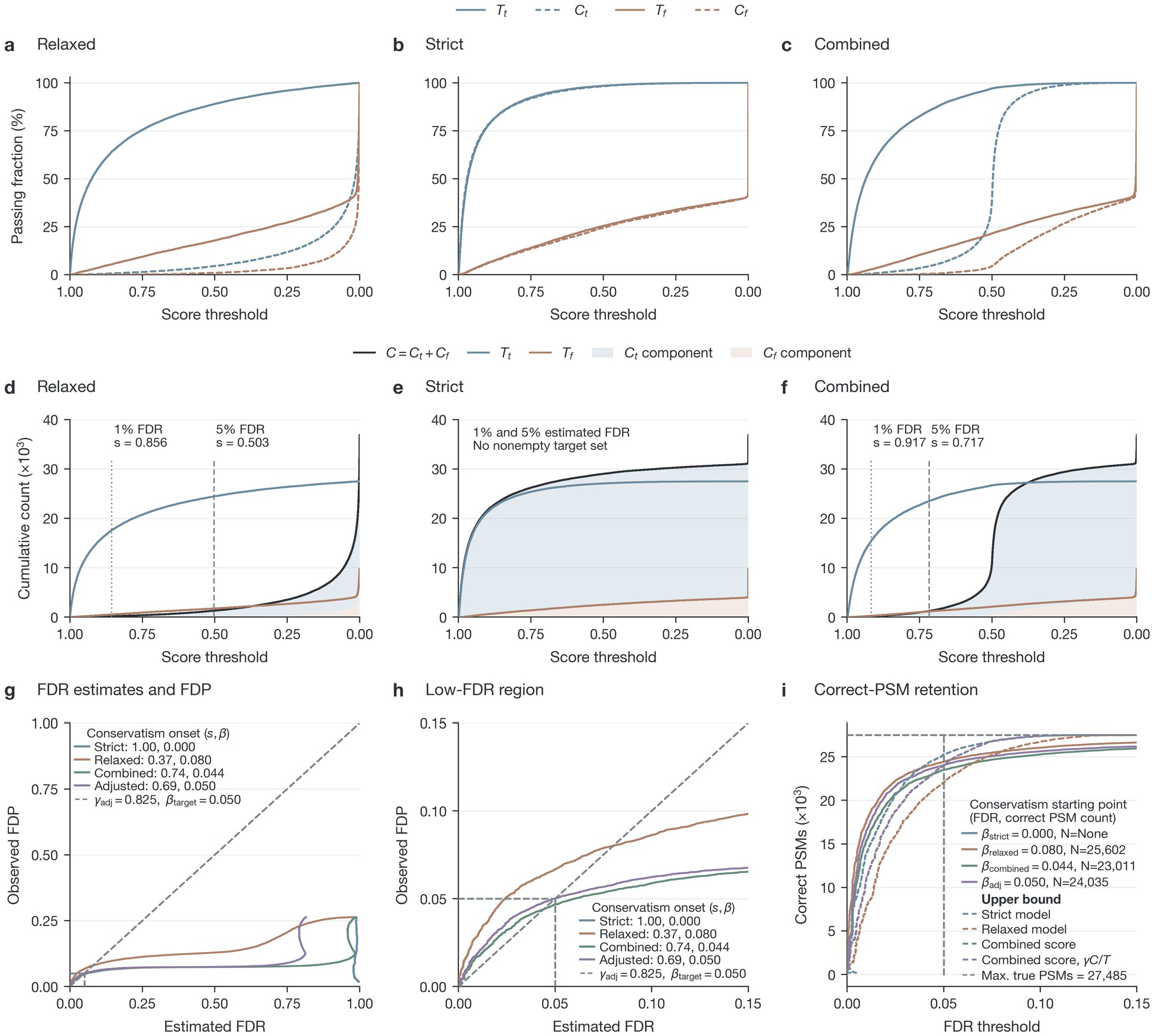
Score distributions, cumulative counts and FDR estimation on the GraphNovo three-species test set. a-c, Within-group fractions passing each score threshold for relaxed, strict and combined scoring, respectively. d-f, Corresponding cumulative target and counterpart counts, grouped by agreement or disagreement with database-search references. Score cutoffs retain complete score ties and use the lowest score yielding a nonempty target set at the indicated FDR threshold, before calibration. g, Estimated FDR versus observed FDP. h, Enlarged view of the low-FDR region. i, Retained reference-correct PSMs across FDR thresholds.

We first train a relaxed multilayer perceptron (MLP), assigning a positive training label only to *T_t_* and negative labels to *T_f_*, *C_t_* and *C_f_*. This objective preferentially enriches correct targets at high scores, as illustrated by the four groups’ threshold-passing fractions (**Fig. 2a**). However, it also pushes *C_t_*, which supplies the additional counts needed for conservative estimation, toward lower scores. In the analyzed data, the complete counterpart count *C* (*s_i_*) remained below *T_f_* (*s_i_*) over part of the threshold range and entered a sustained conservative region as the threshold was relaxed (**Fig. 2d**). Wherever *C_f_* (*s_i_*)< *T_f_* (*s_i_*), the additional count *C_t_* (*s_i_*) must cover the difference *T_f_* (*s_i_*)-*C_f_* (*s_i_*) for the complete counterpart count to satisfy *C* (*s_i_*)*≥T_f_* (*s_i_*). Thus, enriching correct targets alone is insufficient for efficient conservative selection: the ranking must also make sufficient counterpart counts available to estimate the errors among the accepted targets. Suppressing these compensating counts can leave the estimate optimistic in the high-score region and limit correct-PSM retention when a conservative QC criterion is imposed.

To improve this ordering, we additionally train a strict MLP, assigning positive labels to *T_t_* and *C_t_* and negative labels to *T_f_* and *C_f_*. This objective builds on the correctness information carried by the prediction confidence and integrates the other QC features to distinguish correct from incorrect predictions across target and counterpart entries. Under strict scoring, the threshold-passing curves of *T_t_* and *C_t_*, and separately those of *T_f_* and *C_f_*, were closely aligned (**Fig. 2b**). The alignment of the error-associated groups provides empirical support for the equal threshold-passing probabilities assumed by the strict-model lemma. In these data, the corresponding total counts were also close, *C* (*s_i_*)*≈T* (*s_i_*), giving an unadjusted FDR estimate near 1 over the illustrated operating range (**Fig. 2e,g**). Consequently, strict scoring cannot serve as a standalone score for useful low-FDR filtering with this estimator. Its discrimination of prediction correctness nevertheless provides complementary information for improving the relaxed ranking.

We therefore combine the two MLP outputs through a linear weighted score,

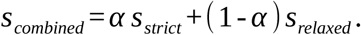

with *α* = 0.5 by default. This combination introduces the strict model’s correctness information into the relaxed model’s preferential scoring of correct targets. In the combined distribution, *T_t_* remained enriched at high scores, while *C_t_* occupied an intermediate score region above the error-associated groups *T_f_* and *C_f_* (**Fig. 2c**). This ordering made the compensating counterpart counts available earlier in the ranking. The cumulative curves show the resulting relationship between the complete counterpart count *C* (*s_i_*), the incorrect-target count *T_f_* (*s_i_*) and the retained correct-target count *T_t_* (*s_i_*) (**Fig. 2f**). Together, these distributions and counts explain how the combined score refines the relaxed ranking for count-based QC. The 1% and 5% guides in **Fig. 2d-f** locate the score cutoffs selected by each unadjusted estimate, allowing the corresponding score prefixes to be compared across rankings.

For quality control at a user-specified target FDR, *β_target_*, we calibrate the estimated FDR on the fixed combined ranking using a multiplicative adjustment factor, *γ* :

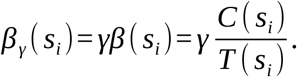

Using database-search identifications as references, π-NovoQC determines *γ* and selects the most permissive feasible score threshold at which the calibrated estimate satisfies the target-FDR criterion while remaining conservative in the reference-labelled calibration data. The unadjusted and adjusted combined-score curves show how this calibration changes the estimated FDR relative to the observed FDP (**Fig. 2g,h**), and the corresponding retention curves show the numbers of correct PSMs retained after filtering (**Fig. 2i**). The factor rescales the FDR estimate, leaving the model scores, their ranking and the cumulative counts at a fixed score threshold unchanged. Applying the calibrated estimate can change the selected score cutoff and, consequently, the retained PSM set.

For comparison, we also applied the same calibration procedure to the relaxed ranking, determining *γ* separately for each scoring strategy using the same reference population. The target-FDR scan compared the adjustment factors, selected score thresholds and fractions of reference-correct PSMs excluded by QC across the two strategies. At a nominal FDR of 5%, the combined and relaxed rankings used adjustment factors of 0.825 and 2.057, respectively. Both calibrated estimates were conservative at the selected thresholds in these reference data, while combined scoring retained 24,035 reference-correct PSMs, compared with 22,077 for relaxed scoring. This comparison shows that relaxed scoring can also support conservative selection after calibration, but incorporating strict-score information improves correct-PSM retention at the same nominal FDR target. The corresponding calibration scans and species-level results are shown in Supplementary Fig. 1 and Supplementary Table 1.

Applying the fitted models and adjustment factors unchanged to each species preserved the retention advantage: combined scoring recovered 602, 757 and 599 additional reference-correct PSMs in *A. thaliana*, *C. elegans* and *E. coli*, respectively. These results support combined scoring followed by *γ* calibration.

### 2. Feature construction and dual-level quality control in π-NovoQC

π-NovoQC implements CCT using features from the acquisition-specific DDA and DIA π-xNovo models developed here (Fig. 3a). The encoder represents MS/MS peaks, and the decoder predicts the peptide from the carboxyl (C) to amino (N) terminus. Decoder cross-attention links residue positions to spectral peaks, providing paired target and counterpart features. The DDA and DIA sequencing-model architectures are illustrated in Supplementary Figs. 2 and 3.

**Fig. 3.**
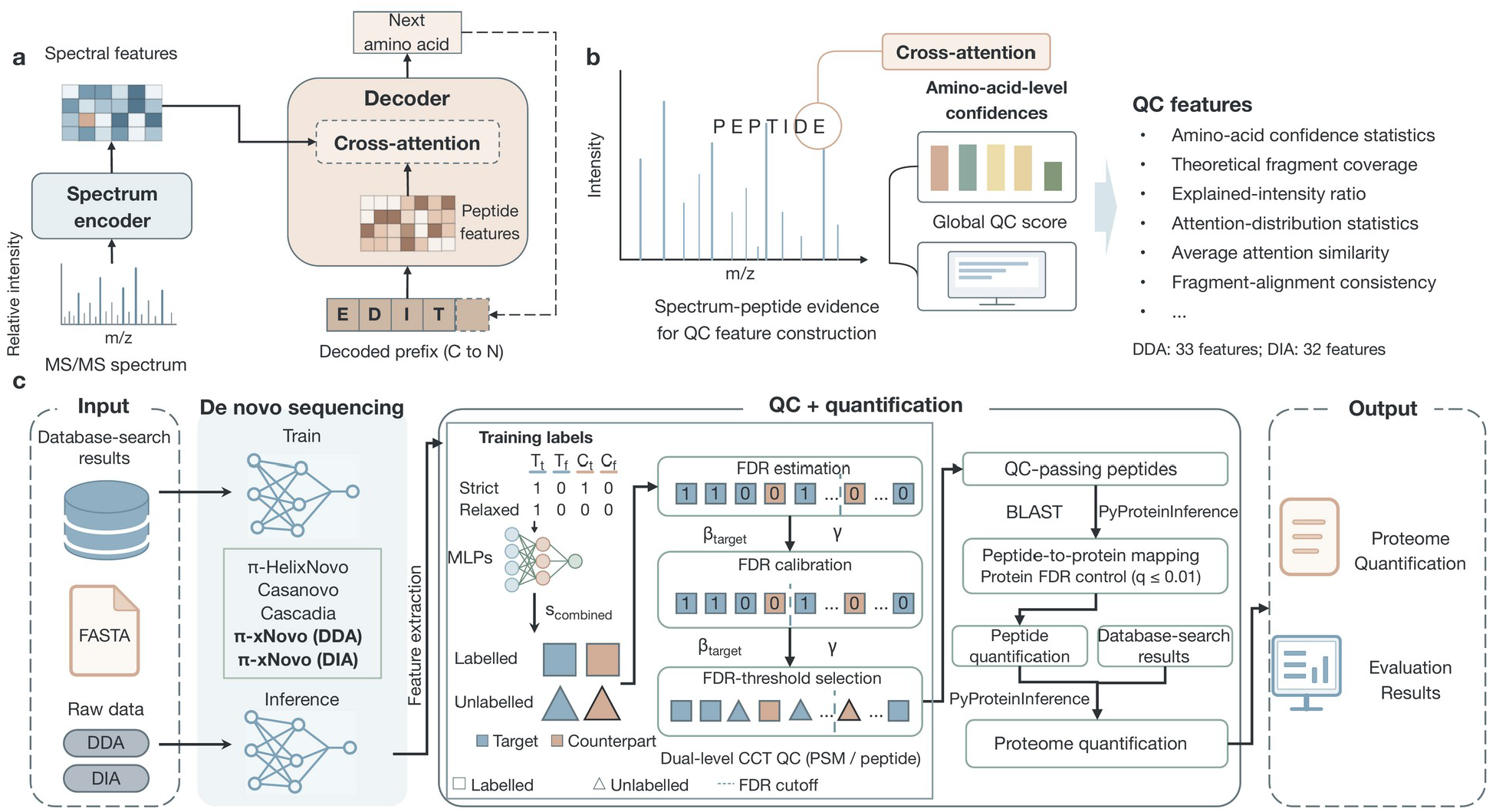
Cross-attention-based feature construction and the π-NovoQC workflow. a, Encoder-decoder core of π-xNovo. b, Construction of paired QC feature vectors. The DDA and DIA vectors contain 33 and 32 scalars, respectively; DIA omits precursor-mass error. c, Identification and quantification workflow integrating PSM- and peptide-level QC, protein inference and MS1 quantification.

For a candidate peptide of length *L*, row 0 of the decoder cross-attention matrix corresponds to the precursor query, rows 1 to *L* correspond to the input residues in decoding order, and the final row corresponds to the stop token. Target features are extracted from rows 0 : *L*, and counterpart features from rows 1 : 1 + *L*, using zero-based, end-exclusive indexing (**Fig. 1b**). These windows represent adjacent decoder states with different sequence prefixes while attending to the same experimental spectrum. Fragment-ion evidence can inform multiple residue positions through the decoder’s sequence context. The one-position shift therefore varies the attention-derived features while preserving shared spectrum-peptide matching information, including precursor-mass error, fragment coverage and explained intensity.

The paired QC feature vectors summarize residue-level evidence, spectrum-peptide agreement and information from decoder cross-attention (Fig. 3b). The DDA and DIA implementations use 33 and 32 scalar features, respectively; the DIA implementation omits precursor-mass error.

Database-search identifications provide reference labels for fitting strict and relaxed PSM-scoring MLPs and calculating *γ*. Target and counterpart entries are pooled and ranked by their combined scores. Their cumulative counts give the adjusted FDR estimate, and the most permissive threshold satisfying the target FDR selects PSMs (**Fig. 3c**).

To construct the peptide-level population, π-NovoQC collects the pair identifiers associated with target or counterpart entries passing the PSM threshold and recovers the corresponding entries and feature vectors from both sides. The recovered population is then collapsed by full modified peptide sequence, retaining the representative with the highest combined PSM score for each sequence. Strict and relaxed peptide-level MLPs are fitted to the resulting peptide-level population, followed by separate calculation of peptide-level *γ* and the peptide selection threshold (Fig. 3c). Peptide-level score and threshold comparisons are provided in Supplementary Figs. 5–8.

Peptides predicted from spectra without database-search identifications and accepted by dual-level CCT QC are mapped by BLAST [23] to a target-decoy protein database and subjected to parsimonious protein inference using PyProteinInference [24]. A separate protein q-value threshold of 0.01 is applied during protein inference. Peptide intensities are extracted from MS1 chromatograms and integrated with database-search evidence for protein quantification. Protein groups are re-inferred after merging this evidence, without an additional protein-FDR filter at the quantification stage (Fig. 3c).

We evaluated peptide-level QC on the GraphNovo three-species benchmark described above. After PSM selection, paired-feature recovery and sequence collapsing, 16,736 reference-correct peptides remained available, defining the attainable upper bound for the subsequent peptide stage. At a nominal peptide FDR of 5%, the unadjusted relaxed-score estimate remained optimistic, whereas the combined-score estimate was markedly overconservative. After *γ* calibration, combined scoring retained 16,691 reference-correct peptides, close to this conditional upper bound, compared with 14,479 retained using calibrated relaxed scoring. Peptide-level calibration thus preserved the retention advantage of combined scoring while reducing the loss of correct peptides caused by an overconservative unadjusted estimate.

Across the PSM/peptide FDR grid, combined scoring used peptide-level *γ* below 1, whereas relaxed scoring often required *γ* > 1 under stringent filtering. Combined scoring retained approximately 41% of reference-correct peptides at 1%/1% target FDRs and over 90% when the PSM and peptide targets reached at least 6% and 2%, respectively.

A separate fixed-score analysis varied the strict-score weight from 0 to 1 in 0.1 increments, using the same peptide population obtained after default PSM selection. At weight 0.5 and nominal 5% FDR, 24,024 correct PSMs and 16,690 correct peptides were retained, with observed FDPs of 5.0097% and 4.1851%. Relative to relaxed scoring, PSM AUC increased from 0.9352 to 0.9514 and peptide AUC from 0.7455 to 0.9328, alongside improved correct-identification retention.

We compared eight DDA feature configurations at both QC levels using five paired seeds: the complete 33-feature set, five feature-group omissions, and compact sets of 11 and 19 features. The compact sets retained 73.81% and 86.50% of the correct peptides retained by the complete set. Removing attention geometry caused the largest decrease among single-group omissions, retaining 87.54% of correct peptides. Thus, attention-derived features contributed information beyond prediction confidence and spectrum support.

The corresponding eight DIA configurations (All 32, five group omissions, Compact 10/18) were evaluated on all eligible candidates in each of six Astral samples [25] with seed 454. Each configuration ran both QC stages and constructed peptides from its own PSM-filtered results. Across 48 comparisons, observed FDP exceeded 5% in nine PSM and seven peptide results, reaching 5.0476% and 5.5057%, respectively. Identification yield must therefore be interpreted alongside calibration at each QC level.

### 3. Applications of CCT across sequencing models and acquisition settings

We evaluated CCT across sequencing models, training datasets and acquisition settings (Fig. 4). Each application fitted QC models and calculated *γ* using corresponding features and database-reference PSMs, with a target FDR of 5% at both QC levels unless specified otherwise.

**Fig. 4.**
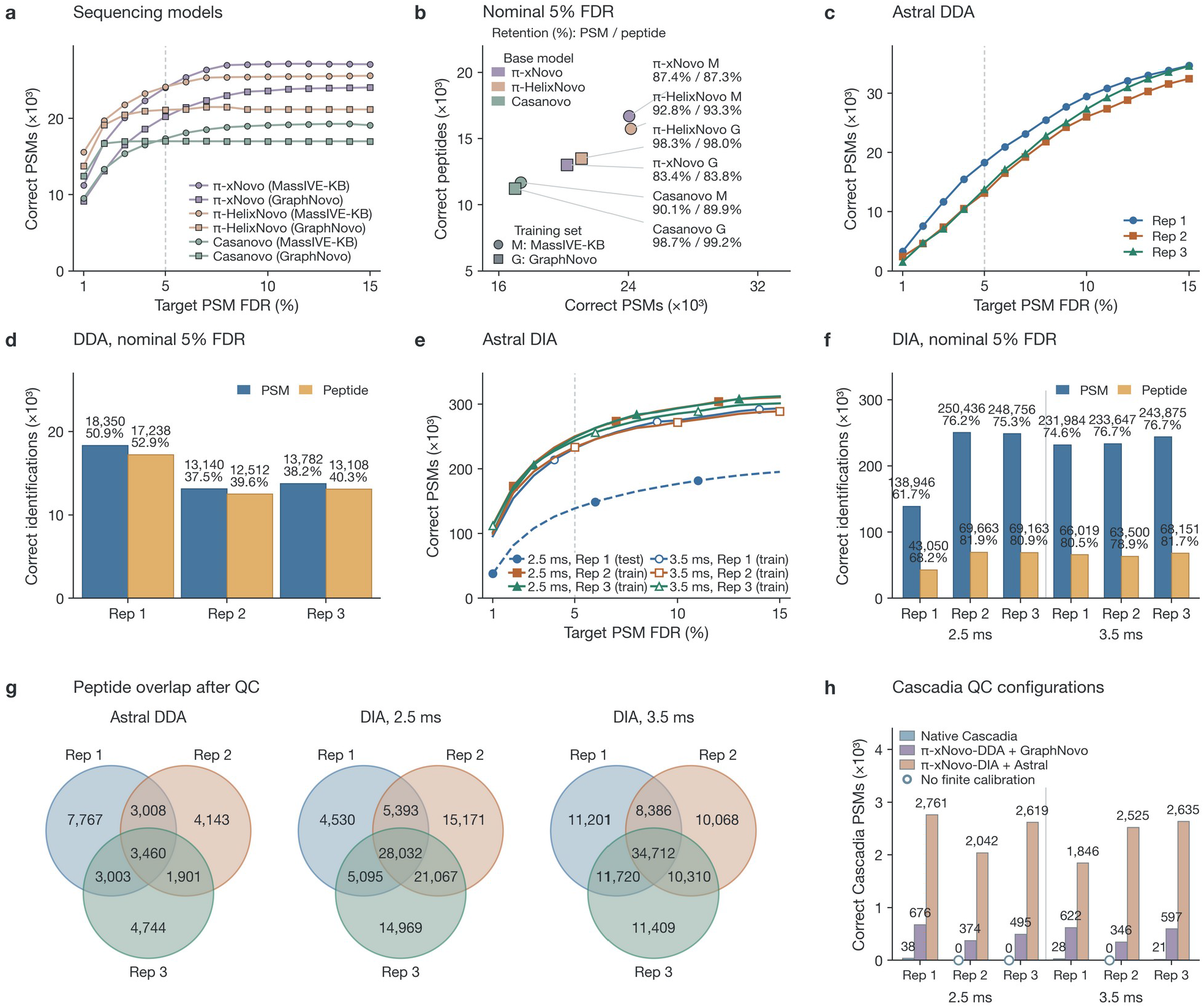
CCT quality control across sequencing models, training datasets and acquisition settings. a, Reference-correct PSMs retained across target FDR values on the GraphNovo test set for π-xNovo, π-HelixNovo and Casanovo. b, Reference-correct PSM and peptide counts and retention at nominal 5% FDR for the same six configurations. c,d, Corresponding PSM threshold curves (c) and PSM/peptide results at nominal 5% FDR (d) for three Astral DDA technical replicates. e,f, Corresponding results for six Astral DIA runs at maximum injection times of 2.5 or 3.5 ms. Only 2.5-ms Rep 1 was held out from sequencing-model training. g, Peptide overlap across replicates after per-entry filtering at an adjusted FDR estimate ≤0.05; sets retain modification states and distinguish I from L. h, Reference-correct Cascadia PSMs retained using native features alone or combined with auxiliary reference-PSM feature sets extracted by π-xNovo. Auxiliary references are excluded from displayed counts. The three native-feature runs shown at zero had no finite adjustment factor. Correctness denotes agreement with database-search references. Retention denominators are the correct PSM or unique peptide counts before QC. Panels b, d and f use the final cumulative point whose adjusted FDR estimate differs from 0.05 by less than 0.0001.

We first compared π-xNovo, π-HelixNovo [7] and Casanovo [26] on the GraphNovo three-species test set [8] using sequencing models trained on either MassIVE-KB [22] or GraphNovo [8] data. The MassIVE-KB-trained models achieved mean peptide recalls of 74.77%, 69.87% and 67.46%, respectively, compared with 65.33%, 57.33% and 50.36% for the same models trained on GraphNovo. π-xNovo achieved the highest peptide recall among the three sequencers under both training conditions and was used as the primary sequencing model in π-NovoQC. The retained-PSM curves and peptide-level results illustrate how the sequencing model and its training data affect the identification yield after QC (**Fig. 4a,b**).

At an adjusted FDR estimate ≤5%, MassIVE-KB training increased retained reference-correct PSM and peptide counts for both Casanovo and π-HelixNovo relative to GraphNovo training.

In nine-species leave-one-species-out evaluation [5], each sequencer was trained on eight species and tested on the ninth. Downstream QC used an 8:2 reference split within that test species for model training/validation, followed by *γ* calculation and QC on the full eligible population. Mean PSM retention was 85.56%, 70.44% and 67.44% for π-xNovo, π-HelixNovo and Casanovo. Retention correlated negatively with *γ* at both QC levels.

The Astral DDA sequencer [25] was trained on approximately 500,000 Astral PSMs. Across three technical replicates, curves had similar shapes but different yields: PSM retention ranged from 37.5% to 50.9% and peptide retention from 39.6% to 52.9% (**Fig. 4c,d**).

Astral DIA analyses used maximum injection times of 2.5 and 3.5 ms. Only the 2.5-ms Rep 1 sample was held out from sequencing-model training; these designations refer to the sequencer rather than downstream QC fitting. This held-out run retained 61.7% of reference-correct PSMs and 68.2% of reference-correct peptides. The three 3.5-ms sequencing-model training runs retained 74.6%–76.7% and 78.9%–81.7%, respectively (**Fig. 4e,f**).

To assess peptide recovery across technical replicates, we calculated the fraction of each replicate’s peptides shared with at least one other replicate and averaged it across the three replicates. These overlap rates were 61.88% for Astral DDA, 82.02% for 2.5-ms DIA and 83.48% for 3.5-ms DIA (**Fig. 4g**). The sets use per-entry adjusted FDR ≤0.05 and exact modified-sequence strings, whereas **Fig. 4b,d,f** use near-threshold cumulative selection.

For Cascadia predictions [12], we compared the native Cascadia feature set alone with analyses that combined it with an auxiliary reference-PSM feature set extracted by π-xNovo-DDA from GraphNovo or by π-xNovo-DIA from Astral data. The combined feature sets were used for QC model training and FDR calibration. Only retained reference-correct Cascadia PSMs are reported (Fig. 4h; Methods). The per-run counts are listed in Supplementary Table 5.

Across six runs, these settings retained 87, 3,110 and 14,428 correct Cascadia PSMs, respectively (Fig. 4h). Native-feature calibration yielded no usable threshold in three runs. Because both feature and reference sources changed, their individual contributions cannot be separated.

Finally, we examined dual-level QC in the DHP [27] and Orbitrap DIA [28] datasets. Across 2,458 DHP raw files, the numbers of retained reference-correct PSMs and peptides were strongly correlated (*r ≈* 0.99). Mean PSM and peptide retention rates were 94.94% and 99.47%, respectively. PSM retention was negatively correlated with *γ*, with most samples having *γ* < 1; the peptide-level correlation was weaker (*r ≈* −0.30). Across 186 Orbitrap DIA samples, retained reference-correct PSM and peptide counts were also positively correlated (*r ≈* 0.42), and peptide retention was 96.91%. The corresponding results are shown in Supplementary Fig. 13.

### 4. Comparison of π-NovoQC with existing quality-control methods

We compared π-NovoQC with NovoBoard [16] and Winnow [17] using candidate PSMs from the GraphNovo three-species test set and 85 DHP HCD evaluation files. Each method applied a nominal 5% PSM FDR threshold. GraphNovo was used for QC-method development and NovoBoard parameter selection. The DHP evaluation files comprised 27 Trypsin, 26 AspN and 32 LysN runs. Winnow was trained and validated using 22 separate DHP files, and its fitted model was then applied to the 85 evaluation files. Evaluation datasets and method comparisons are detailed in Supplementary Tables 6–13; NovoBoard parameter selection is shown in Supplementary Fig. 19.

On GraphNovo, π-NovoQC retained 25,146 PSMs at 5.0107% observed FDP, compared with 23,446 at 6.7304% for NovoBoard and 8,581 at 2.4356% for Winnow. Thus, π-NovoQC retained 7.25% more PSMs than NovoBoard with lower FDP, while its higher yield than Winnow accompanied higher FDP.

Across the 85 DHP files, π-NovoQC, NovoBoard and Winnow retained 567,258, 162,507 and 467,876 PSMs, with pooled observed FDPs of 1.5628%, 8.9664% and 4.7476%, respectively (**Fig. 5a**). Filtering was performed separately within each raw file. The pooled FDP is the total reference-incorrect count divided by the total retained count across files.

**Fig. 5.**
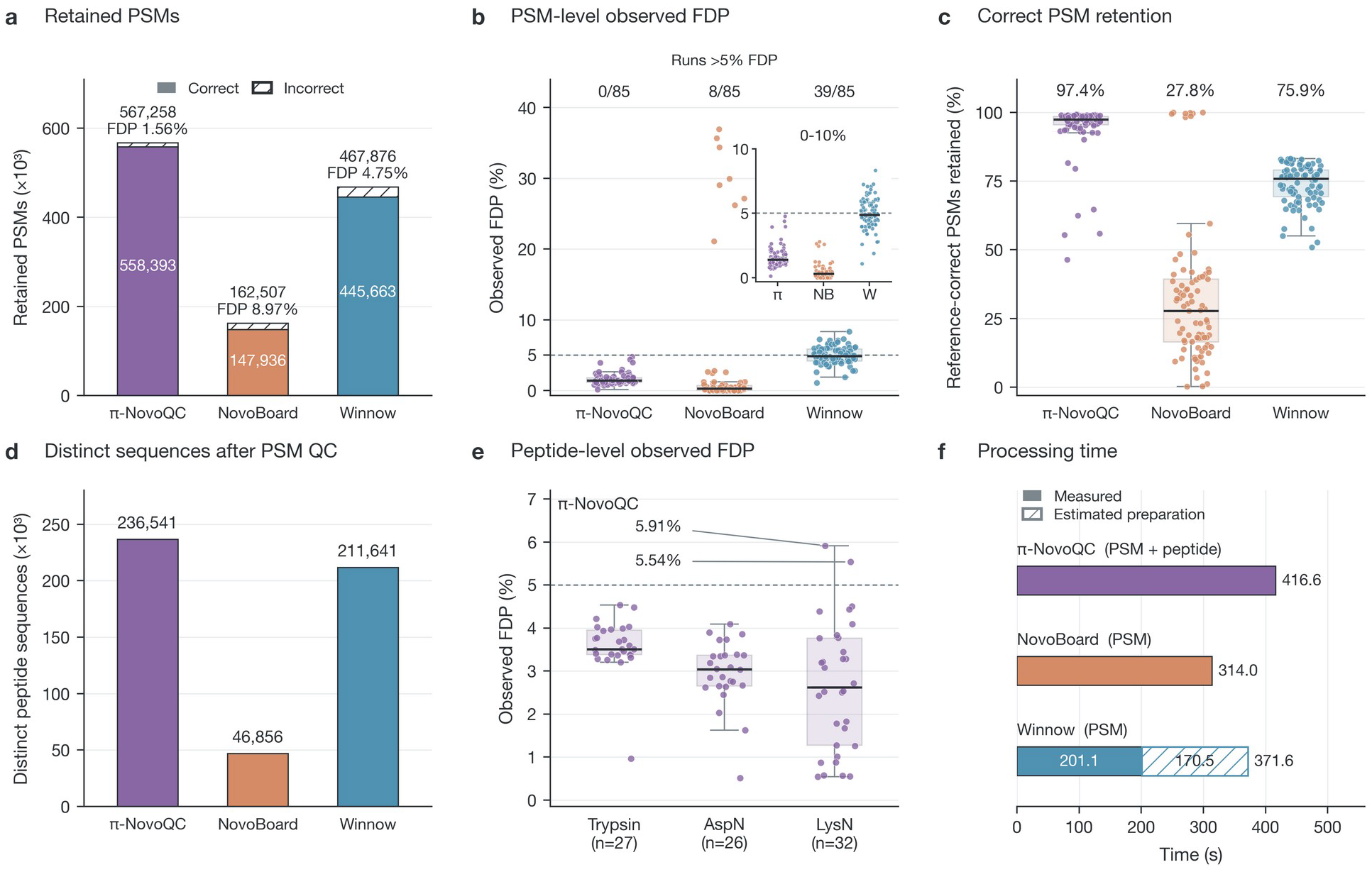
Comparison of quality-control methods on 85 DHP HCD evaluation raw files. a, Filtering was performed separately for each raw file. Retained reference-correct and reference-incorrect PSMs at nominal 5% PSM FDR. Pooled observed FDP is the total number of reference-incorrect PSMs divided by the total retained PSM count. b, Per-file PSM observed FDP. c, Retention relative to each file’s initial reference-correct PSMs. d, Peptide sequences pooled across PSM-filtered files and deduplicated by sequence and modification state, before peptide-level QC. e, Peptide-level observed FDP for π-NovoQC by protease. f, Processing time for 9,777 candidate PSMs from the representative DHP raw file. π-NovoQC includes both QC stages; NovoBoard and Winnow end at PSM filtering. Winnow preparation time is estimated from measured throughput. In b, c and e, each observation is one raw file; boxes span the interquartile range, central lines indicate medians and whiskers extend to observations within 1.5 interquartile ranges. Correctness denotes agreement with database-search references. Timing measurements and the steps included are detailed in Supplementary Tables 14 and 15.

All 85 π-NovoQC PSM results had observed FDP below 5% (maximum, 4.7686%), whereas eight NovoBoard and 39 Winnow results exceeded 5% (Fig. 5b). π-NovoQC retained 3.49 and 1.21 times as many PSMs as these methods, respectively, with lower pooled FDP. Per-file correct-PSM retention is shown in **Fig. 5c**.

After PSM filtering, sequences were harmonised by full modified sequence and represented by each method’s highest-scoring PSM. π-NovoQC, NovoBoard and Winnow covered 16,705, 15,740 and 5,985 distinct sequences on GraphNovo. Pooling the 85 DHP files before deduplication yielded 236,541, 46,856 and 211,641 sequences (Fig. 5d). These coverage summaries precede peptide-level QC.

The native two-stage π-NovoQC workflow additionally applies peptide-level scoring, *γ* calibration and filtering after paired-feature recovery and sequence collapsing. Retained peptide counts summed to 302,573 across the 85 DHP files, with pooled FDP of 3.3784%. Eighty-three files had peptide FDP below 5%; the HepG2 LysN fraction-21 and fraction-26 raw files reached 5.5355% and 5.9144% (Fig. 5e).

Database-search identification fractions ranged from 2.0662% to 28.0146% across the DHP evaluation files [27]. Sixty-nine files used their own reference PSMs and 16 used auxiliary references. Reference availability is shown in Supplementary Figs. 14 and 15.

The raw file for fraction 11 of the LysN-digested HepG2 sample contained 2,023 database identifications among 97,910 MS2 spectra. Using its own references, π-NovoQC retained 702 PSMs and 367 peptides at observed FDPs of 0.1425% and 0.5450%. The raw file for fraction 5 of the AspN-digested GM12878 sample contained 1,860 identifications among 73,407 spectra and retained 1,129 PSMs and 454 peptides at FDPs of 0.7972% and 3.0837%.

We timed 9,777 candidate PSMs from the HCD raw file for fraction 10 of the AspN-digested GM12878 sample on one Tesla V100 32 GB GPU with eight CPU threads. π-NovoQC required 416.566 seconds from spectrum reading through feature generation, scoring-model training and both QC stages; NovoBoard required 313.986 seconds through PSM filtering. Winnow required 201.086 seconds for local feature recalculation and filtering, or an estimated 371.599 seconds including candidate generation and intensity/iRT/RT preparation (Fig. 5f).

### 5. Proteomic application and orthogonal validation of π-NovoQC

To assess whether FDR-controlled de novo sequencing could recover proteomic evidence beyond conventional database-search results, we applied π-NovoQC to the DHP dataset [27], comprising 2,458 raw files and approximately 1.4 × 10^8^ database-unidentified spectra. The π-xNovo model was trained on multi-protease data acquired using higher-energy collisional dissociation and electron-transfer dissociation. Candidate peptides were processed using the combined score and the adjusted FDR estimate at a target FDR of 5%, followed by protein inference at a separate protein-level FDR threshold of 1%. BLAST comparisons were used to classify recovered peptides as fully matched, partially matched or unmatched under the applied alignment criteria, while single-amino-acid polymorphism (SAP) peptides were analyzed separately (**Fig. 6**).

**Fig. 6.**
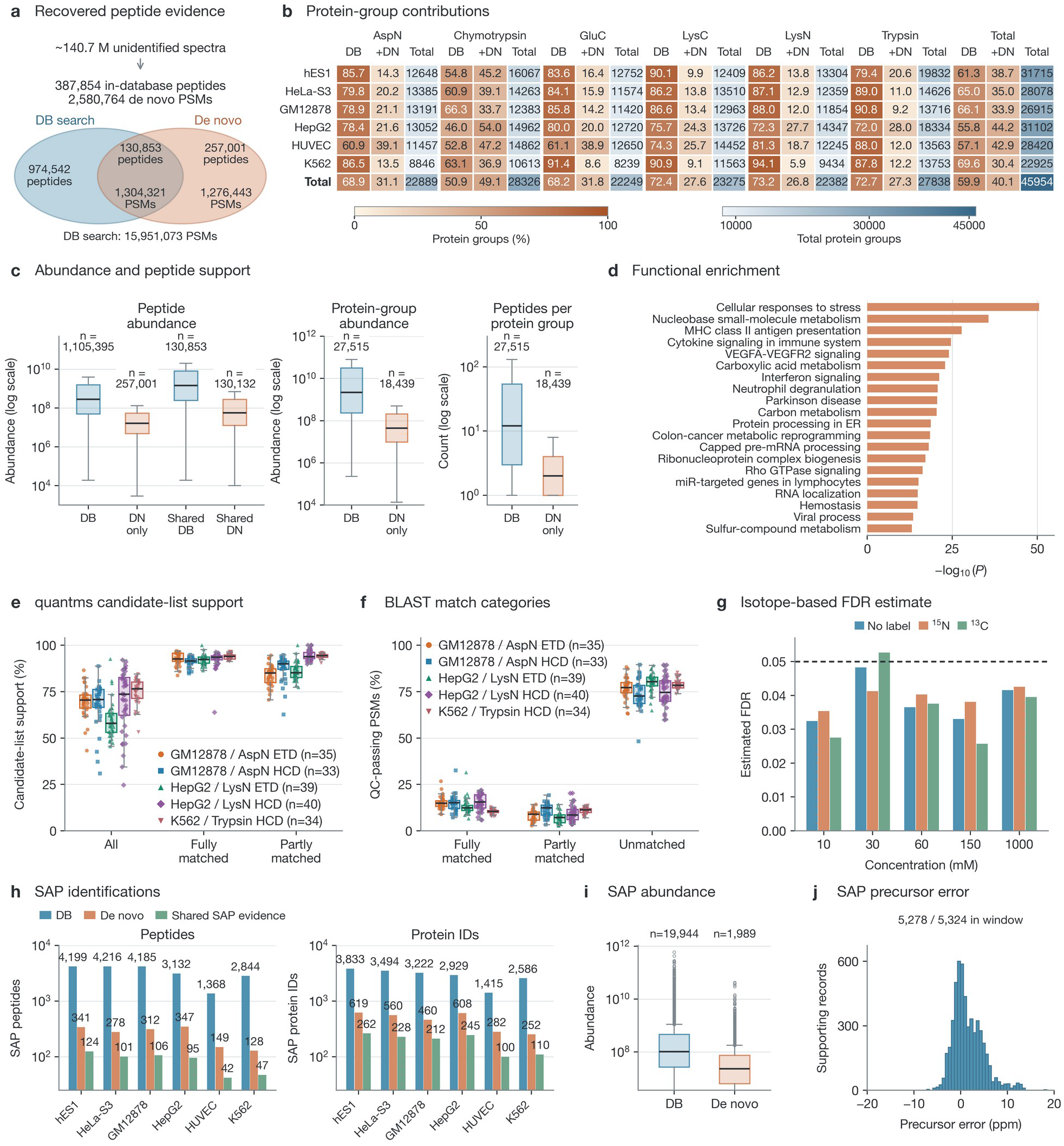
DHP proteomic analysis and isotope-labeling validation of π-NovoQC. a, Overlap of database-search and de novo peptide identifications with positive quantified intensity. b, Protein-group contributions across DHP cell lines and proteases. c, Peptide abundance, protein-group abundance and peptides per group. d, Functional enrichment of the most abundant 10% of de novo-supported protein groups; P values are from unadjusted hypergeometric tests. e, quantms candidate-list support for recovered PSMs. f, BLAST alignment categories. Each point in e and f represents one raw file. g, Isotope-based FDR estimates for QC-passing PSMs in the Dong-Ecoli-QE dataset. h, SAP peptide and variant-protein counts. i, SAP peptide abundance. j, Precursor mass errors of SAP-supporting assignments. Boxplots show medians and interquartile ranges, with whiskers extending to observations within 1.5 interquartile ranges.

Of 390,319 in-database peptides passing QC, 387,854 had positive quantified intensity, supported by 2,580,764 PSMs (**Fig. 6a**). Quantification included 2,456 of the 2,458 QC raw files. Database-search results shared 130,853 peptides; another 257,001 were recovered by de novo sequencing. Of the latter, 175,739 mapped to DB-supported protein groups and 81,262 supported 18,439 additional groups. These peptides were absent from database-search results, not from the reference protein sequences. Additional spectral and quantitative comparisons are provided in Supplementary Figs. 16–18.

Joint parsimony-based inference yielded 45,954 protein groups: 27,515 DB-supported and 18,439 additional groups supported by de novo evidence (**Fig. 6b**). Contributions from de novo evidence ranged from 5.9% to 54.0% across proteases, with the largest contribution for chymotrypsin. Recovered peptides and protein groups generally had intensities one to two orders of magnitude lower and fewer supporting peptides than database-search identifications (**Fig. 6c**), extending coverage into lower-abundance regions.

To characterize the functional composition of de novo-supported protein groups, we selected the most abundant 10% and retained Ensembl and UniProt entries, yielding 1,055 input protein identifiers. Metascape [29] identified enriched terms related to stress responses, proteostasis, RNA metabolism, translational regulation and immune processes (**Fig. 6d**). Applying a common intensity floor of 8.14 *×* 10^8^, defined by the minimum intensity in the de novo subset, gave an approximately 9.5-fold lower median abundance than for DB-supported proteins (1.70 × 10^9^ versus 1.62 × 10^10^). These enrichments describe the composition of recovered groups without establishing a causal biological relationship.

We further used an independent database-search workflow to provide additional search-based support for the recovered predictions. We selected 181 raw files spanning five experimental conditions and re-searched their spectra with quantms [30] after adding the predicted peptides to the search space. Support required agreement in peptide sequence and spectrum index within the list of up to five candidate peptides per spectrum before 1% FDR filtering. The mean per-file support fraction among QC-passing de novo PSMs was 68.28%; the corresponding means for the fully and partly BLAST-matched categories were 92.38% and 89.28%, respectively (**Fig. 6e**). The mean BLAST-unmatched fraction was 76.95% (**Fig. 6f**). These are candidate-list support fractions, rather than top-ranked identification accuracy. quantms was not used for model training, threshold selection or FDR calibration, and lack of search support does not by itself establish that a prediction is incorrect. The experimental conditions and numbers of raw files are summarized in Supplementary Table 16.

To assess the reliability of de novo identifications recovered from spectra not identified by database searching, we used isotope-labeling evidence from the Dong-Ecoli-QE dataset [40]. Expected isotope-shifted precursor signals allow a predicted peptide to be checked independently of whether its spectrum has a database-search identification. Across five concentrations and three isotope channels, the isotope-based FDR estimates for the complete QC-passing de novo PSM sets ranged from 2.568% to 5.267% (mean, 3.815%; **Fig. 6g**). These sets included PSMs from both database-identified and database-unidentified spectra; the missingness model is described in Methods.

Variant-database analysis yielded 1,989 distinct raw-file/peptide combinations and 4,116 peptide/variant-protein pairs, averaging 173 additional SAP identifications per cell line (**Fig. 6h**). De novo SAP evidence occupied lower abundance ranges, although workflows used different aggregation units and unnormalized abundances (**Fig. 6i**). Of 5,324 SAP-supporting assignments, 5,278 had absolute precursor error below 20 ppm (**Fig. 6j**). Assignments include repeated peptide support and are not independent peptide counts.

Across 64 DHP conditions, π-NovoQC peptide intensities correlated with MaxQuant [31, 32] at a median Pearson r of 0.886, comparable to PANDA [33] (0.894; paired t-test, P = 0.97). Median protein-level correlation with PANDA was 0.818. Across six Astral DIA runs, peptide-intensity correlations with DIA-NN [34] were approximately 0.57–0.59.

## Discussion

De novo sequencing extends peptide identification beyond database-search results, but downstream use requires reliable false-discovery assessment. Model confidence ranks candidates without establishing the error proportion among retained predictions. CCT connects ranking to FDR estimation through paired target and counterpart counts, providing a basis for selection at a specified FDR.

The unadjusted estimate *C* / *T* uses cumulative counterpart (*C*) and target (*T*) counts. Counterparts associated with correct predictions supply additional counts; the estimate is conservative when the complete counterpart count exceeds the incorrect-target count. Database-search identifications provide reference labels for defining the four groups, fitting scoring models and calculating the adjustment factor. Consequently, the observed FDP used for evaluation measures agreement or disagreement with the database-search peptide for the same spectrum. Once the models and adjustment factor have been determined, selection of candidates with unknown correctness requires their scores and cumulative counts rather than individual reference labels.

Separating ranking from FDR calibration is central to π-NovoQC. Relaxed scoring enriches correct targets but can suppress the counterpart counts needed for conservative estimation. Strict scoring contributes correctness information, although similar target/counterpart distributions limit standalone filtering with *C* / *T*. Their combination improves ranking; the adjustment factor rescales FDR estimates without changing scores or ordering. It can nevertheless change the selected threshold and retained set.

The dual-level implementation extends the same logic from PSMs to peptide sequences. At the PSM level, π-NovoQC uses paired target and counterpart features to select candidate spectrum-peptide matches. After PSM filtering, paired entries and their feature vectors are recovered, and the entries are collapsed by full modified peptide sequence. This changes both the candidate population and the counting unit, so PSM-level FDR control does not automatically establish peptide-level FDR control. π-NovoQC therefore fits peptide-level scoring models and calculates a separate adjustment factor on the resulting population.

Feature-removal experiments further indicate that the practical performance of the scoring procedure depends on retaining sufficient model and spectrum information. Compact feature sets reduced discrimination and correct-identification retention relative to the complete input, while removing attention geometry produced one of the largest reductions among the tested feature groups. The DDA and DIA sensitivity experiments also show that calibration performance varies across feature configurations, samples and acquisition modes. These results support the contribution of attention-derived features to QC ranking within the evaluated decoders, feature definitions and data populations.

Applications across sequencing models demonstrate flexibility when compatible features and suitable reference PSMs are available. Sequencer generalisation and downstream QC training remain distinct, as in the nine-species evaluation. The Cascadia analyses show that auxiliary reference-PSM feature sets can support QC of existing Cascadia predictions. Because the auxiliary feature sets differed in both the extraction model and the reference data, the comparison does not isolate the contribution of either factor.

Comparisons with NovoBoard and Winnow illustrate why yield and calibration must be considered jointly. π-NovoQC retained more PSMs at lower observed FDP than NovoBoard on GraphNovo, whereas its higher yield than Winnow accompanied higher FDP. It achieved higher retention and lower pooled FDP than both on DHP. PSM retention, per-file FDP and sequence coverage describe different aspects of performance; peptide-level results require separate assessment, including the DHP files that exceeded the nominal target.

DHP results show that de novo sequencing can recover low-abundance evidence from database-unidentified spectra and support additional protein groups. Many recovered sequences nevertheless occur in the reference database. Absence from database-search results therefore demonstrates additional analytical recovery, without by itself establishing sequence novelty.

The validation analyses provide complementary evidence. quantms re-search identified many of the recovered peptides in its candidate lists, particularly those classified as fully or partially matched by BLAST. quantms was not used for π-NovoQC training, threshold selection or FDR calibration, so this analysis provides additional search-based support rather than an independent definition of truth. The isotope-labelled Dong-Ecoli-QE experiment provides a different type of check based on isotope-shifted precursor signals. The resulting FDR estimates account for undetected isotopologues using a reference-derived missingness coefficient and the reference-error assumption specified in Methods. The SAP analysis extends the application to variant-supported evidence, but SAP assignments supported by variant databases and mass-spectrometric evidence still require genetic or transcriptomic validation.

The current implementation depends on database-reference PSMs for fitting and calibration. Sparse, biased or unrepresentative references can affect performance near the target FDR, and reference-relative observed FDP inherits database-search errors. Database-unidentified spectra and noncanonical peptides are distinct populations. The isotope experiment included identified and unidentified spectra but did not separately quantify noncanonical-peptide error or false-negative rates.

CCT cannot recover a correct sequence absent from the sequencer’s predictions. Applying the current attention-based features to another sequencing model requires a compatible decoder; models without this structure require a different feature-extraction design. QC models and adjustment factors should be fitted using suitable references from the current data source; uncalibrated transfer across instruments, acquisition modes or populations may alter the relationship between counterpart and incorrect-target counts.

Database search and de novo sequencing can be combined: the former supplies reference PSMs, while the latter expands evidence from unidentified spectra. After dual-level QC, mapping accepted peptides to genomic or transcriptomic protein databases supports downstream inference. Protein-group and variant interpretations remain conditional on mapping, parsimony, database completeness and orthogonal biological evidence.

CCT thus provides a framework for FDR estimation and threshold selection at the PSM and peptide levels. Integrating accepted de novo predictions with database-search evidence extends peptide coverage for protein inference and quantitative analysis.

## Methods

### π-NovoQC quality-control workflow

After autoregressive peptide prediction, decoder cross-attention was evaluated using the predicted peptide and the same experimental spectrum. For a candidate of length *L*, target features were extracted from attention rows 0 : *L*, and counterpart features from rows 1 : 1 + *L*, using zero-based, end-exclusive indexing. Row 0 corresponded to the precursor query, and subsequent rows to the input residues in decoding order. The two feature representations therefore used different decoder-query positions while retaining the same candidate sequence and experimental spectrum. Feature definitions and ablation groups are summarized in Supplementary Note 2 and Supplementary Table 2; feature-ablation results are shown in Supplementary Figs. 9 and 10.

For spectra with database-search identifications, agreement between the predicted peptide and the database-search peptide for the same spectrum determined the assignments to the four entry groups. Target and counterpart entries associated with reference-consistent predictions were assigned to *T_t_* and *C_t_*, respectively; those associated with reference-inconsistent predictions were assigned to *T_f_* and *C_f_*. The strict scoring MLP used *T_t_* and *C_t_* as positive training examples and *T_f_* and *C_f_* as negative examples. The relaxed MLP used only *T_t_* as positive examples and the other three categories as negative examples. The default combined score was the arithmetic mean of the two model outputs, *s_combined_* = 0.5 *s_strict_* + 0.5 *s_relaxed_*.

Target and counterpart entries were pooled and ranked in descending order of the combined score. At a threshold *_s_*, *T* (*s*) and *C* (*s*) counted all target and counterpart PSM-level entries, respectively, with scores at least *s*. For *T* (*s*)> 0, the unadjusted FDR estimate was *C* (*s*) / *T* (*s*), and reference labels supplied the observed FDP, *T_f_* (*s*) / *T* (*s*), for calibration. The adjustment factor *γ* was calculated using the reference-labelled population. The calibrated estimate was *γC* (*s*) / *T* (*s*). The most permissive feasible score threshold satisfying the user-specified target-FDR criterion was selected, and target predictions passing that threshold were retained. The calibration left the scores, ranking and counts at a fixed threshold unchanged, but could change the selected threshold and retained set. Relaxed-score comparisons used a separately calculated *γ* for the relaxed ranking; strict-score comparisons used the unadjusted counterpart-count estimate. Calibration conditions and proofs are set out in Supplementary Note 1.

To construct the peptide-level population, pair identifiers associated with either target or counterpart entries passing the PSM threshold were collected, and the corresponding entries and feature vectors from both sides were recovered. The recovered population was ranked by combined PSM score and collapsed by full modified peptide sequence, retaining the highest-scoring representative for each sequence. Repeated PSM scores were not summed or averaged. Strict and relaxed peptide-level MLPs were fitted again using the available reference labels for this population, followed by separate calculation of peptide-level *γ* and the selection threshold. Processing was performed within each sample, except for the predefined pooled GraphNovo three-species evaluation. Unless otherwise specified, the target FDR was 5% at both the PSM and peptide levels.

Peptides predicted from database-unidentified spectra and accepted by both CCT QC stages were mapped by BLAST [23] to a target-decoy protein database and subjected to parsimonious protein inference using PyProteinInference [24]. A separate protein q-value threshold of 0.01 was applied during protein inference. For quantification, MS1-based peptide intensities and database-search evidence were integrated, and protein groups were re-inferred from the merged peptide evidence without an additional protein-FDR filter at that stage. Additional workflow details are provided in Supplementary Note 2; datasets and preprocessing are described in Supplementary Note 3 and Supplementary Table 3.

### De novo sequencing model framework: π-xNovo

#### π-xNovo-DDA model

π-xNovo-DDA uses a Transformer encoder-decoder architecture [6] to predict peptide sequences from MS/MS spectra. The spectral encoder represents peak masses and intensities together with precursor information. The decoder predicts amino acids autoregressively from the peptide C terminus to the N terminus. Decoder cross-attention connects sequence positions with spectral evidence and provides information for paired QC feature construction (Fig. 3a,b). Training uses masked representations, and relative positional encoding follows established Transformer approaches [35, 36].

#### π-xNovo-DIA model

π-xNovo-DIA extends the encoder-decoder framework to chimeric DIA spectra through a precursor-conditioned peak-selection stage followed by peptide prediction. The peak selector identifies spectral evidence associated with precursor candidates, and the peptide decoder predicts sequences from the selected evidence. Acquisition-specific models were used for Orbitrap and Astral DIA data.

#### Training strategy

Sequencing models were trained on the datasets specified for each evaluation. Training and evaluation used NVIDIA Tesla V100 GPUs. The DDA and DIA models were trained separately for their acquisition settings; pretrained DDA models were used to initialize the corresponding DIA models. Sequencing-model training is distinct from the downstream training of QC scoring models and FDR calibration.

#### QC with auxiliary reference-PSM feature sets

Across six Astral DIA runs, we evaluated QC of Cascadia predictions [12] using three settings: the native Cascadia feature set alone; the native set combined with an auxiliary GraphNovo reference-PSM feature set extracted by π-xNovo-DDA [8]; and the native set combined with an auxiliary Astral reference-PSM feature set extracted by π-xNovo-DIA [25]. Auxiliary reference PSMs contributed to the QC analysis but were excluded from the reported counts of retained reference-correct Cascadia PSMs (Fig. 4h). The comparison evaluates these combined settings, rather than isolating the contribution of the auxiliary reference data or the model used to extract their features.

#### Baseline models and settings

We used published code and pretrained models for π-HelixNovo [7], Casanovo [26] and Cascadia [12]. The π-HelixNovo and Casanovo evaluations used the GraphNovo [8] and nine-species [5] datasets. Cascadia was fine-tuned on the Orbitrap DIA [28] and Astral DIA [25] datasets. These sequencing models were evaluated with CCT at both the PSM and peptide levels.

For Winnow [**17**], candidate peptide strings were converted to the residue and modification representation required by the supported Prosit models [37]; unsupported sequence or charge combinations were excluded before prediction. Predicted fragment intensities and indexed retention times (iRTs) were matched to candidates by sequence and the model inputs. Winnow features combined these predictions with observed spectra, precursor metadata and native sequencing confidence. Retention-time regression was fitted separately to the complete eligible candidate population in each raw file to calculate the iRT-error feature.

The Winnow probability calibrator was fitted using 275,832 valid PSMs from 22 DHP development files [27]: seven Trypsin, seven AspN and eight LysN files. A split with random seed 42 assigned 248,249 PSMs to training and 27,583 to validation; labels were defined by agreement with MaxQuant reference identifications [31, 32]. The 85 evaluation files (27 Trypsin, 26 AspN and 32 LysN files; 1,138,277 candidates) were excluded from calibrator training. The fitted calibrator was applied unchanged to evaluation features. Within each evaluation file, Winnow’s non-parametric FDR procedure fitted the calibrated confidences, produced PSM posterior error probabilities (PEPs), FDR estimates and q-values, and returned the confidence cutoff at the nominal 5% FDR. All candidates at or above this cutoff were retained. Reference correctness labels were joined only after the cutoff had been fixed.

For NovoBoard [**16**], decoy spectra were generated by peak replacement and processed with the same sequencing model as the target spectra. For a spectrum containing N peaks and a replacement fraction r, ⌊ *N* (1 - *r*) ⌋ original peaks were sampled and retained, and the remaining peaks were sampled from a common pool of target-spectrum peaks. Python’s random generator was reinitialised with seed 99 for each spectrum, and the resulting peaks were sorted by m/z. Target and decoy predictions were scored by mean token log probability and ranked together in descending order. Cumulative target and decoy counts were evaluated at the final position of each group of tied scores; the most permissive endpoint with a decoy-to-target ratio no greater than the nominal FDR was selected. Filtering was performed separately within each evaluation file.

The NovoBoard peak-replacement fraction was selected on the GraphNovo development population [8] by comparing observed FDP with estimated FDR across candidate settings. The selected fraction, 0.5, was then fixed for DHP evaluation without retuning on the evaluation results.

The Winnow and NovoBoard comparisons used their PSM-level filtering workflows without adding a separate peptide-level model-training, calibration or filtering stage. For nonredundant sequence comparisons after PSM filtering, full modified peptide sequence representations were harmonised and deduplicated; the highest-scoring PSM under each method was retained as the representative of each sequence. These sequence-level summaries were kept distinct from the peptide-level results of the native two-stage π-NovoQC workflow.

### Database re-search with quantms

To provide additional search-based support for π-NovoQC-recovered peptides, we used quantms (v1.3) [30] to re-search 181 DHP raw files [27] spanning five combinations of cell line, protease and fragmentation method. The search database was constructed by combining the human reference database with de novo-predicted peptide sequences and removing redundant sequences. Searches were performed using MS-GF+ [38] and Comet [39], with up to five candidate PSMs reported per MS/MS spectrum. Precursor and fragment mass tolerances were set to 20 ppm and 0.35 Da, respectively. Enzyme specificity was set according to the protease used, allowing up to two missed cleavages. Carbamidomethylation of cysteine was specified as a fixed modification, and methionine oxidation and protein N-terminal acetylation were specified as variable modifications.

For the support analysis, we used the search-output lists containing up to five candidate peptides per spectrum before 1% FDR filtering. For each sample, we calculated the proportion of π-NovoQC-recovered peptides found in these candidate lists and summarized the proportions as the mean across samples. Support proportions were also calculated for the peptide categories defined by BLAST alignment [23]. Candidate-list matches provided additional search-based corroboration and were not interpreted as identifications passing a 1% FDR threshold. The quantms results were not used for model training, QC threshold selection or FDR calibration, and absence from the candidate lists was not treated as evidence that a recovered prediction was incorrect.

#### Isotope-labeling validation of de novo identifications

We used the Dong-Ecoli-QE dataset [40] to examine the reliability of de novo identifications recovered from database-unidentified spectra using expected isotope-shifted precursor signals. The dataset contains unlabeled, ¹⁵N-labeled and ¹3C-labeled *E. coli* samples mixed at a ratio of 1:1:1. For a true peptide detected in the unlabeled channel, the corresponding ¹⁵N and ¹3C isotopologues are expected at predictable precursor m/z shifts. However, labeled counterparts can be missed owing to noise, co-elution or limited sensitivity. We therefore first estimated a missingness coefficient (θ) from the reference PSM table used in this analysis and then used θ to convert isotopologue missingness among QC-passing de novo PSMs into an empirical FDR estimate.

Estimation of θ from the reference PSM table. The calculation used the reference PSMs corresponding to each raw file and isotope channel. For each reference PSM, signals at the theoretical isotope-shifted precursor m/z values were quantified using ±3 min retention-time and ±20 ppm precursor-m/z windows. An expected isotopologue was considered missing when its MS1 area was zero or absent. Let 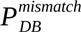 denote the fraction of reference PSMs lacking the expected isotopologue. The database-search reference labels used a 0.001 search-FDR threshold; this threshold is distinct from the mixture-model error proportion. For the estimates shown in **Fig. 6g, t**he source analysis uses a reference error-proportion parameter of *ρ*_0_ = 0.05 and models the probability of isotopologue missingness as one for incorrect PSMs and θ for correct PSMs. Under this mixture model, the missingness coefficient is:

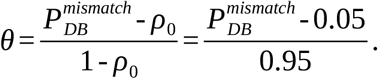

Empirical FDR estimation for QC-passing de novo PSMs. We applied π-xNovo-DDA (MassIVE-KB) to all spectra and performed PSM-level CCT QC at the target threshold (*q_target_* = 0.05) to obtain the QC-passing de novo PSM set, which included both database-identified and database-unidentified spectra. For each QC-passing PSM (sequence, modifications, and charge), we computed the theoretical precursor m/z of its ¹⁵N or ¹3C isotopologues from the peptide elemental composition and isotope mass shifts and quantified the labeled counterparts with π-NovoQC using the same RT and m/z tolerances. Let 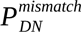 denote the fraction of QC-passing de novo PSMs whose expected isotopologues were missing. We used the corresponding reference-derived θ as the missingness probability for correct de novo PSMs within each raw file and isotope channel. Under the same mixture model, we estimated the empirical FDR as:

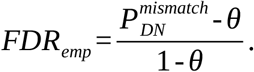

We calculated the empirical FDR separately for each raw file and isotope channel using the corresponding θ. The five concentration conditions and three isotope channels yielded 15 estimates for the complete QC-passing de novo PSM sets.

## Data availability

The nine-species dataset is available through PRIDE under PXD005025, PXD004948, PXD004325, PXD004565, PXD004536, PXD004947, PXD003868, PXD004467 and PXD004424. The GraphNovo dataset is available at https://doi.org/10.5281/zenodo.8000316. MassIVE-KB raw files and filtered identifications were obtained from the “All Candidate library spectra” section of MassIVE Knowledge Base spectral library v1 (https://massive.ucsd.edu/ProteoSAFe/static/massive-kb-libraries.jsp). DHP, Astral DDA/DIA and Dong-Ecoli-QE data are available through ProteomeXchange under PXD024364, PXD046453 and PXD008782, respectively. The Orbitrap DIA dataset is available from the CPTAC consortium under PDC000200. The timsTOF DDA data have been deposited in iProX under Project ID IPX0018169000 and ProteomeXchange identifier PXD080632. Supplementary Information accompanies this preprint. Source Data will accompany the journal publication.

## Code availability

Code availability will be documented in the journal publication.

## Acknowledgements

This work has been supported by the Prevention and Control of Emerging and Major Infectious Diseases-National Science and Technology Major Project (2026ZD01999704), the Major Project of Guangzhou National Laboratory (Grant No. SRPG22-001), direct national funding from the Chinese Ministry of Technology to Peng Cheng Laboratory, the Chinese Ministry of Technology to Peng Cheng Laboratory (PCL2025A09), the National Key Research and Development Program of China (2025YFA1309300), the National Natural Science Foundation of China (32088101) and the CAMS Innovation Fund for Medical Sciences (CIFMS, 2019-I2M-5-063).

## Author Contributions

Y.W. and C.C. designed the study and co-supervised the project. Z.L. designed CCT and π-NovoQC, performed the experiments, and wrote the initial manuscript. C. D. performed the quantms re-search analysis. T.L., T.Y., Y.Y., and Y.L. collected the data and performed the data analysis. L.X. helped to improve the model. F.H. and Y.H. provided suggestions for the study. All authors helped revise the manuscript and approved the final manuscript.

