## Supplementary Information for "False discovery rate control enables trustworthy deep-learning-based de novo peptide sequencing"

Zhendong Liang[1,2], Chengxin Dai[3,4], Tianze Ling[3,5], Tingpeng Yang[1], Yun Yang[4], Yeye Leng[4], Linhai Xie[3,4], Yonghong He[2], Fuchu He[3,4], Yu Wang[1*], Cheng Chang[3,4*]

1 Peng Cheng Laboratory, Shenzhen, 518055, China

2 Tsinghua Shenzhen International Graduate School, Shenzhen, 518055, China

3 State Key Laboratory of Medical Proteomics, National Center for Protein Sciences (Beijing), Research Unit of Proteomics-driven Cancer Precision Medicine (Chinese Academy of Medical Sciences), Beijing 102206, China

4 International Academy of Phronesis Medicine (Guangdong), 510320, Guangdong, China

5 School of Life Sciences, Tsinghua University, Beijing, 100084, China

* **To whom correspondence should be addressed:**

Yu Wang,

Cheng Chang,

The supplementary information includes:

- **Supplementary Note 1**: Counterpart Calibration Theory (CCT) Proofs
- **Supplementary Note 2: π-NovoQC workflow and feature definitions**
- **Supplementary Note 3**: Datasets and Preprocessing
- **Supplementary Note 4**: End-to-end Validation of π-NovoQC
- **Supplementary Tables 1–16**.
- **Supplementary Figs. 1–19**.

### Supplementary Note 1: Counterpart Calibration Theory (CCT) Proofs

In database search, target-decoy competition (TDC) uses decoy matches retained at a score threshold to estimate incorrect target matches retained at the same threshold [1, 2]. For balanced target and decoy search spaces, this estimation relies on the scoring and competition procedure giving incorrect assignments comparable probabilities of being retained on either side. In de novo sequencing, candidate generation and the resulting error distribution depend on the model's use of spectral evidence, learned contextual dependence among amino acids within a single peptide and decoding procedure. Reversing, shuffling or replacing residues in a predicted peptide does not by itself ensure that the resulting sequences represent these errors and their scores. Nevertheless, the principle of using references to estimate error counts can still inform de novo quality control.

CCT develops this strategy using paired target and counterpart features obtained from the same experimental spectrum and predicted peptide. These features are supplied to scoring models, whose outputs are used to rank the target and counterpart features. At a common score threshold, the total retained counterpart count is used to estimate the number of incorrect retained targets, and dividing this count by the total number of retained targets defines the FDR estimate. Counterparts associated with correct predictions can contribute an additional count. Under appropriate scoring and cumulative-count conditions, this contribution may lead to overestimation of the number of incorrect predictions and thereby make the FDR estimate conservative. Database-search identifications provide reference labels for scoring-model training and for evaluating and calibrating the resulting FDR estimates; correctness and observed FDP in these analyses are defined relative to those labels. The results below formulate the conditions required for conservative estimation and establish the associated calibration and threshold-selection procedures for a fixed score ranking.

#### 1. Problem definition

We consider the candidate peptide identifications produced by a de novo sequencing model. CCT associates each candidate with paired target and counterpart features extracted from the same experimental spectrum and predicted sequence. Target features represent the candidate identifications to be assessed, while counterpart features provide paired comparators for estimating error counts.

In reference-labelled data, each predicted peptide is compared with the database-search identification from the same spectrum. Combining the resulting agreement status with the feature type gives four feature groups:

- $T_t$: target features associated with predictions that agree with the database-search reference.
- $T_f$: target features associated with predictions that disagree with the database-search reference.
- $C_t$: counterpart features paired with target features in $T_t$.
- $C_f$: counterpart features paired with target features in $T_f$.

The subscripts $t$ and $f$ indicate the agreement status of the predicted peptide shared by each target-counterpart pair. These groups describe feature type and agreement with the reference; the binary labels used to train the scoring models depend on their respective training objectives.

The objective is to select a score threshold at a user-specified target FDR while retaining as many correct target identifications as possible. In reference-labelled data, the observed false discovery proportion (FDP) among retained target predictions is used to evaluate the conservativeness of the FDR estimate.

#### 2. Basic definitions and notation

##### 2.1 Scoring models and score thresholds

Let $M$ denote a fixed scoring scheme: relaxed, strict or combined. The relaxed model is trained with $T_t$ as the positive class and $T_f$, $C_t$ and $C_f$ as the negative class. The strict model is trained with $T_t$ and $C_t$ as the positive class and $T_f$ and $C_f$ as the negative class. The combined score is a linear weighted combination of the outputs of these two models.

For a target or counterpart feature vector $x$, let $s_M(x)$ denote its score under scheme $M$, with larger scores ranked higher. For a fixed input population, the distinct score values define a finite ordered set of thresholds,

$$S_M = \{s_1, s_2, \ldots, s_n\}, s_1 > s_2 > \cdots > s_n.$$

Here, $s_1$ is the most stringent threshold and $s_n$ is the most permissive threshold. At threshold $s$, all target and counterpart features satisfying

$$s_M(x) \geq s$$

are retained. Features with equal scores are retained together. If $u \leq s$, threshold $u$ is at least as permissive as threshold $s$, and every feature retained at $s$ is also retained at $u$. Threshold values are interpreted within their respective scoring schemes, because score distributions and rankings may differ between schemes.

##### 2.2 Cumulative counts

For a fixed scoring scheme $M$ and a common threshold $s \in S_M$, let

$$T_t^M(s), T_f^M(s), C_t^M(s), C_f^M(s)$$

denote the cumulative numbers of features from $T_t$, $T_f$, $C_t$ and $C_f$, respectively, whose scores satisfy $s_M(x) \geq s$. The total numbers of retained target and counterpart features are

$$T^M(s) = T_t^M(s) + T_f^M(s),$$
$$C^M(s) = C_t^M(s) + C_f^M(s).$$

When the scoring scheme is clear from the context, we omit the superscript $M$.

##### 2.3 Observed FDP, estimated FDR and adjusted estimated FDR

For a threshold $s$ with $T^M(s) > 0$, the observed FDP in reference-labelled data is the proportion of retained target predictions that disagree with the database-search reference:

$$FDP_M(s) = \frac{T_f^M(s)}{T^M(s)}.$$

CCT uses the total retained counterpart count $C^M(s)$ to estimate the number of incorrect retained target predictions. The corresponding FDR estimate is

$$\beta_M(s) = \frac{C^M(s)}{T^M(s)}.$$

For a fixed scoring scheme and a specified target FDR, a multiplicative correction factor $\gamma > 0$ gives the adjusted estimate

$$\beta_{adj,M}(s) = \gamma \beta_M(s) = \gamma \frac{C^M(s)}{T^M(s)}.$$

The factor $\gamma$ rescales the estimated FDR on the fixed score ranking. The scores, their ordering and the cumulative counts at a given score threshold remain unchanged. The conditions used to determine $\gamma$ are specified in the subsequent calibration theorems.

Thresholds with $T^M(s) = 0$ are not used as QC output thresholds. For notational completeness, $\beta_M(s)$ and $FDP_M(s)$ may be set to zero by convention at these empty-target thresholds.

#### 2.4 Conservativeness and the difference function

For a fixed scoring scheme $M$ and a threshold $s$ with $T^M(s) > 0$, the unadjusted FDR estimate is pointwise conservative relative to the observed FDP in reference-labelled data if

$$\beta_M(s) \geq FDP_M(s).$$

Because the two quantities share the same positive denominator $T^M(s)$, this condition is equivalent to

$$C^M(s) \geq T_f^M(s).$$

For the adjusted estimate, the corresponding condition is

$$\beta_{adj,M}(s) \geq FDP_M(s),$$

or equivalently,

$$\gamma C^M(s) \geq T_f^M(s).$$

For the unadjusted estimate, define the difference function

$$\Delta_M(s) = \beta_M(s) - FDP_M(s) = \frac{C^M(s) - T_f^M(s)}{T^M(s)}.$$

Thus, $\Delta_M(s) \geq 0$ indicates conservative estimation at $s$. A positive value indicates that the estimate exceeds the observed FDP, zero indicates equality, and a negative value indicates underestimation. In a $\beta$–FDP plot, conservative points lie on or below the identity line $y = x$ when estimated FDR is on the horizontal axis and observed FDP is on the vertical axis.

#### 2.5 Local conservativeness, endpoint conservativeness and the effective local conservative starting point

A scoring scheme $M$ is locally conservative from threshold $s$ if its unadjusted estimate is pointwise conservative at $s$ and at every more permissive threshold:

$$\Delta_M(u) \geq 0, \text{for all } u \in S_M \text{ with } u \leq s.$$

Equivalently, $C^M(u) \geq T_f^M(u)$ throughout this set of thresholds. The portion of the ordered threshold set extending from $s$ to the most permissive threshold $s_n$ is called a conservative suffix. Endpoint conservativeness holds when

$$\Delta_M(s_n) \geq 0.$$

In this case, at least the one-threshold suffix $\{s_n\}$ is conservative.

Provided that a conservative suffix exists, the effective local conservative starting point is defined as the most stringent threshold that begins such a suffix:

$$s_M^{LC} = max\left\{ s \in S_M : \Delta_M(u) \geq 0 \text{ for all } u \in S_M \text{ with } u \leq s \right\}.$$

The maximum is taken within the scoring scheme's own score scale, where larger thresholds are more stringent. Starting from $s_n$, one can identify this point by moving toward more stringent thresholds while requiring every threshold in the resulting suffix to remain conservative. Because the $\beta$–FDP curve may cross the identity line multiple times, $s_M^{LC}$ is determined by this sustained suffix condition, rather than by the first crossing of the plotted curve.

We also define the estimated-FDR coordinate associated with the starting point:

$$\beta_M^{LC} = \beta_M\left(s_M^{LC}\right).$$

This is the value of the unadjusted FDR estimate at $s_M^{LC}$. The starting point itself is defined in score-threshold order; the definition does not require estimated FDR to vary monotonically with the score threshold.

#### 2.6 Observed-to-estimated ratio

For adjustment-factor calibration, define the observed-to-estimated ratio at a threshold with $T^M(s) > 0$ and $\beta_M(s) > 0$:

$$r_M(s) = \frac{FDP_M(s)}{\beta_M(s)} = \frac{T_f^M(s)}{C^M(s)}.$$

The common target-count denominator cancels, so this ratio compares the reference-labelled incorrect target count with the counterpart count used to estimate it.

We use the following boundary conventions:

- If $\beta_M(s) = 0$ and $FDP_M(s) = 0$, set $r_M(s) = 0$.
- If $\beta_M(s) = 0$ and $FDP_M(s) > 0$, set $r_M(s) = \infty$.

Values $r_M(s) \leq 1$ indicate conservative unadjusted estimation, whereas $r_M(s) > 1$ indicate underestimation. For a finite positive correction factor $\gamma$, pointwise conservativeness of the adjusted estimate is equivalent to

$$\beta_{adj,M}(s) \geq FDP_M(s) \Leftrightarrow \gamma \geq r_M(s).$$

If $r_M(s) = \infty$, no finite multiplicative correction can make the estimate conservative at that threshold. Empty-target thresholds remain excluded from QC outputs as specified in Section 2.3.

### 3. Lemma and theorem proofs

#### 3.1 Lemma 1: Strict-model empirical conservativeness

Throughout this lemma, the strict scoring function is fixed. All counts and observed FDP values refer to this scoring scheme and to the reference-based feature groups defined in Section 1.

Let $m_f$ denote the number of pairs associated with incorrect predictions, with both the target and counterpart features of each pair included in the analysed population. Let $s^f_{T,j}$ and $s^f_{C,j}$ denote their strict scores. Assume that, for each pair $j$ and every fixed threshold $s$,

$$Pr\left(s^f_{T,j} \geq s\right) = Pr\left(s^f_{C,j} \geq s\right).$$

This is an assumption about the marginal threshold-passing probabilities of the paired incorrect features. It gives the expectation-level count identity

$$E\left[C_f(s)\right] = E\left[T_f(s)\right].$$

For a finite analysed population and a threshold with $T(s) > 0$, conservativeness holds exactly when

$$C_t(s) + C_f(s) \geq T_f(s),$$

or equivalently,

$$C_t(s) \geq T_f(s) - C_f(s).$$

For the asymptotic statement, additionally consider a sequence of growing analysed populations at a fixed threshold $s$, with $T(s) > 0$, and assume

$$\frac{C_f(s) - T_f(s)}{T(s)} = o(1).$$

Then

$$\Delta_{strict}(s) = \frac{C_t(s)}{T(s)} + o(1),$$

and hence

$$\liminf \Delta_{strict}(s) \geq 0.$$

Here, $o(1)$ denotes deterministic convergence to zero along the specified empirical population sequence, and the $\liminf$ is t aken along that same sequence. This is an asymptotic non-underestimation statement; finite-sample conservativeness is determined by the count condition above.

At the most permissive threshold $s_n$, all analysed features are retained. Complete pairing of the incorrect features gives exact endpoint conservativeness whenever $T(s_n) > 0$.

*Proof.*

For the $j$-th pair associated with an incorrect prediction, define

$$I^T_j(s) = I\left\{s^f_{T,j} \geq s\right\}, I^C_j(s) = I\left\{s^f_{C,j} \geq s\right\}.$$

The corresponding cumulative counts are

$$T_f(s) = \sum_{j=1}^{m_f} I^T_j(s), C_f(s) = \sum_{j=1}^{m_f} I^C_j(s).$$

The assumed equality of passing probabilities and linearity of expectation give

$$E\left[T_f(s)\right]=\sum_{j=1}^{m_f} E\left[I_j^T(s)\right]=\sum_{j=1}^{m_f} E\left[I_j^C(s)\right]=E\left[C_f(s)\right].$$

For any threshold with $T(s)>0$, substituting $C(s)=C_t(s)+C_f(s)$ into the difference function yields the exact decomposition

$$\Delta_{strict}(s)=\beta_{strict}(s)-FDP(s)=\frac{C_t(s)}{T(s)}+\frac{C_f(s)-T_f(s)}{T(s)}.$$

The first term is non-negative. In a finite population, the sum is non-negative exactly when the additional count $C_t(s)$ offsets any deficit of $C_f(s)$ relative to $T_f(s)$. This gives the stated finite-sample condition. Under the additional normalized-imbalance assumption, the second term is $o(1)$, establishing the asymptotic result.

Finally, at $s_n$, both members of every analysed incorrect pair are retained, so

$$C_f(s_n)=T_f(s_n).$$

Consequently,

$$\Delta_{strict}(s_n)=\frac{C_t(s_n)}{T(s_n)}\geq 0 \text{ when } T(s_n)>0.$$

This establishes exact finite-sample conservativeness at the most permissive endpoint.

##### 3.2 Theorem 1: Existence of the relaxed-model local conservative starting point

*Motivation.*

The relaxed scoring model is trained to enrich correct target features at high scores. Its unadjusted counterpart-count estimate may nevertheless be non-conservative at some stringent thresholds. We therefore examine whether the fixed relaxed ranking has a nonempty conservative suffix toward the permissive end. Such a suffix defines an effective local conservative starting point for subsequent calibration and threshold selection.

*Relaxed scoring model.*

The relaxed model assigns label 1 only to $T_t$, and label 0 to $T_f$, $C_t$ and $C_f$. These labels define the training objective; the existence proof below is based on complete pairing and the most permissive endpoint. All counts and observed FDP values in this section are computed under the fixed relaxed ranking, using the boundary conventions in Section 2.3.

*Theorem statement.*

Let $S_{relaxed}$ be the finite ordered threshold set for the relaxed score, with $s_n$ denoting its most permissive threshold. Assume that the target and counterpart features associated with incorrect predictions form complete one-to-one pairs within the analysed population, that all analysed features are retained at $s_n$, and that $T(s_n)>0$.

Then the effective local conservative starting point $s_{relaxed}^{LC}$ exists. Every threshold in the suffix beginning at this point satisfies

$$\Delta_{relaxed}(u)\geq 0, u\in S_{relaxed}, u\leq s_{relaxed}^{LC}.$$

Equivalently, $\beta_{relaxed}(u)\geq FDP(u)$ throughout this suffix.

*Proof.*

Define

$$A_{relaxed} = \{ s \in S_{relaxed} : \Delta_{relaxed}(u) \geq 0 \text{ for every } u \in S_{relaxed} \text{ with } u \leq s \}.$$

At $s_n$, all analysed features are retained. Complete pairing of incorrect features therefore gives

$$C_f(s_n) = T_f(s_n).$$

Using $C(s_n) = C_t(s_n) + C_f(s_n)$ and $T(s_n) > 0$, we obtain

$$\Delta_{relaxed}(s_n) = \frac{C(s_n) - T_f(s_n)}{T(s_n)} = \frac{C_t(s_n)}{T(s_n)} \geq 0.$$

The suffix beginning at $s_n$ contains only $s_n$. Thus, $s_n \in A_{relaxed}$, and $A_{relaxed}$ is nonempty.

Since $S_{relaxed}$ is finite, its nonempty subset $A_{relaxed}$ has a maximum. Consistent with Section 2.5, set

$$s^{LC}_{relaxed} = max\, A_{relaxed}.$$

Membership in $A_{relaxed}$ establishes $\Delta_{relaxed}(u) \geq 0$ at every threshold in the suffix beginning at $s^{LC}_{relaxed}$. This is the most stringent threshold that can begin a conservative suffix, proving the result.

*Interpretation.*

This existence result follows from the pairing and endpoint conditions together with the finite threshold set. The same argument applies to any fixed scoring scheme satisfying these conditions. The relaxed ranking determines the location of its effective local conservative starting point. The conservative suffix may contain only the most permissive endpoint; feasibility at a specified target FDR is determined separately by the calibration and output-selection conditions.

##### 3.3 Theorem 2: Near-strict no-later conservative onset of the combined model

*Combined scoring model.*

The combined model incorporates strict-model information into the relaxed ranking. For each candidate match $x$, let $s_{strict}(x)$ and $s_{relaxed}(x)$ denote its strict-model and relaxed-model scores. For a combination weight $\alpha \in [0,1]$, define

$$s_{combined,\alpha}(x) = \alpha\, s_{strict}(x) + (1-\alpha)\, s_{relaxed}(x).$$

Here $\alpha = 1$ gives the strict-model ranking, whereas $\alpha = 0$ gives the relaxed-model ranking. The combined model is not an average of two FDR curves. It creates a new candidate ranking, and all cumulative counts, observed FDPs and estimated FDRs are recomputed on that ranking.

Because equal-score features are retained together, the proof uses only accepted-target-count coordinates that are attainable at actual score thresholds. For a fixed combination weight $\alpha$, let $K_\alpha$ denote this set of attainable positive target counts. For $k \in K_\alpha$, let $\tau_k(\alpha)$ be the most stringent actual threshold at which exactly $k$ target-side candidates are accepted, and define

$$C_k(\alpha) = C^{combined,\alpha}(\tau_k(\alpha)),$$
$$F_k(\alpha) = T_f^{combined,\alpha}(\tau_k(\alpha)),$$

$$D_k(\alpha)=C_k(\alpha)-F_k(\alpha).$$

Here $C_k(\alpha)=C_{t,k}(\alpha)+C_{f,k}(\alpha)$ includes all retained counterpart features, including the contribution from $C_t$.

*Theorem statement.*

Let $s_{relaxed}^{LC}$ be the effective local conservative starting point of the relaxed model, and let

$$k_{relaxed}^{LC}=T^{relaxed}\left(s_{relaxed}^{LC}\right)$$

be its accepted-target-count coordinate.

For a combination weight $\alpha$, let $s_{combined,\alpha}^{LC}$ be the effective local conservative starting point of the combined model, and define

$$k_{combined}^{LC}(\alpha)=T^{combined,\alpha}\left(s_{combined,\alpha}^{LC}\right).$$

Assume that $k_{relaxed}^{LC}$ is attainable in the strict ranking, that the strict-model conservativeness condition supplied by Lemma 1 holds at every attainable strict-ranking coordinate $k\geq k_{relaxed}^{LC}$, and that this condition is expressed as

$$D_k(1)=C_k(1)-F_k(1)\geq 0.$$

Assume further that the relevant combined-model ranking is stable in a neighbourhood of the strict endpoint $\alpha=1$, including the ordered score groups, the attainable target-count coordinates and the associated cumulative counts.

Then there exists $0\leq\alpha_c<1$ such that, for every

$$\alpha\in[\alpha_c,1],$$

the combined model is conservative at every attainable accepted-target-count coordinate

$$k\in K_\alpha, k\geq k_{relaxed}^{LC}.$$

Consequently,

$$k_{combined}^{LC}(\alpha)\leq k_{relaxed}^{LC}, \alpha\in[\alpha_c,1].$$

Thus, in accepted-target-count coordinates, the near-strict combined model enters a sustained conservative suffix no later than the relaxed model.

*Proof.*

We compare the relaxed and combined models in accepted-target-count coordinates because raw score thresholds are not directly comparable across different rankings.

By Theorem 1, the relaxed model has an effective local conservative starting point $s_{relaxed}^{LC}$. Its accepted-target-count coordinate is $k_{relaxed}^{LC}$, which is the benchmark coordinate from which the relaxed model has a conservative suffix.

Now fix a combination weight $\alpha$. For every attainable coordinate $k\in K_\alpha$, let $\tau_k(\alpha)$ be the most stringent actual threshold at which exactly $k$ target-side candidates are accepted. At this threshold, $C_k(\alpha)$, $F_k(\alpha)$ and $D_k(\alpha)$ are defined as above.

At accepted-target-count coordinate $k$, the denominator of both the estimated FDR and the observed FDP is $k$. Therefore, conservativeness at that coordinate is equivalent to

$$D_k(\alpha) \geq 0.$$

As an actual threshold $u$ is relaxed while the retained target count remains $k$, only counterpart-side candidates can be added. Thus, $C^{combined,\alpha}(u)$ cannot decrease while $T_f^{combined,\alpha}(u)$ remains unchanged. Consequently, $D_j(\alpha) \geq 0$ at every attainable coordinate $j \geq k$ is sufficient to establish conservativeness at every actual threshold $u \leq \tau_k(\alpha)$.

For any two candidate matches $x$ and $y$,

$$\begin{aligned} & s_{combined,\alpha}(x) - s_{combined,\alpha}(y) \\ = & \alpha\left[s_{strict}(x) - s_{strict}(y)\right] \\ + & (1-\alpha)\left[s_{relaxed}(x) - s_{relaxed}(y)\right]. \end{aligned}$$

This is a linear function of $\alpha$. Hence, the relative order of a fixed pair of candidates can change only at isolated values of $\alpha$, unless their relative order remains unchanged throughout the interval.

Because the candidate set is finite, there are only finitely many candidate pairs and hence only finitely many relevant ranking transition points. By the assumed local stability near $\alpha = 1$, there is a neighbourhood of the strict endpoint in which the relevant ordered score groups, attainable target-count coordinates and cumulative rankings do not change. Consequently, for every relevant attainable coordinate $k$,

$$C_k(\alpha) = C_k(1), F_k(\alpha) = F_k(1), D_k(\alpha) = D_k(1).$$

At $\alpha = 1$, the combined score reduces to the strict score:

$$s_{combined,1}(x) = s_{strict}(x).$$

By the strict-model condition in the theorem statement, for every attainable coordinate $k \geq k_{relaxed}^{LC}$,

$$D_k(1) \geq 0.$$

Therefore, for each such attainable coordinate $k$, the local ranking stability gives a value $\alpha_k < 1$ such that

$$D_k(\alpha) \geq 0$$

for every $\alpha \in [\alpha_k, 1]$.

Only finitely many attainable target-count coordinates need to be considered. Let

$$K_{\geq} = \{k \in K_1 : k \geq k_{relaxed}^{LC}\}.$$

Define

$$\alpha_c = \max_{k \in K_{\geq}} \alpha_k.$$

Because $K_{\geq}$ is finite and every $\alpha_k < 1$, we have $\alpha_c < 1$. Hence, for every $\alpha \in [\alpha_c, 1]$ and every attainable coordinate $k \geq k_{relaxed}^{LC}$,

$$D_k(\alpha) \geq 0.$$

Thus, $\tau_{k^{LC}_{relaxed}}(\alpha)$ begins a conservative suffix under the combined ranking. By finiteness of the threshold set and the definition in Section 2.5, the effective local conservative starting point exists and satisfies

$$s^{LC}_{combined,\alpha} \geq \tau_{k^{LC}_{relaxed}}(\alpha).$$

The retained target count cannot increase as the threshold becomes more stringent. Therefore,

$$k^{LC}_{combined}(\alpha) = T^{combined,\alpha}\left(s^{LC}_{combined,\alpha}\right) \leq T^{combined,\alpha}\left(\tau_{k^{LC}_{relaxed}}(\alpha)\right) = k^{LC}_{relaxed}.$$

This completes the proof.

*Interpretation.*

The theorem establishes a near-strict stability result. The relaxed model's onset coordinate is used as a benchmark. When the combined score is sufficiently close to the strict score and the relevant ranking remains stable, the combined model reaches a conservative suffix no later than the relaxed model in accepted-target-count coordinates.

The result does not compare raw score thresholds across models. It also does not imply the same ordering in estimated-FDR coordinates, because the estimated FDR may be locally non-monotone as a function of the accepted-target-count coordinate.

#### 3.4 Theorem 3: Feasibility and optimality of the suffix adjustment factor

*Fixed-ranking adjustment.*

Once a scoring scheme $M$ has fixed the candidate ranking, the adjustment factor acts only on the estimated FDR values. It does not change the scores, the ranked order, the threshold set or any cumulative count. Thus, for a fixed ranking,

$$\beta_{adj,M}(s;\gamma) = \gamma\,\beta_M(s) = \gamma\,\frac{C^M(s)}{T^M(s)}.$$

The theorem applies to any fixed ranking, including the strict, relaxed and combined rankings. Its purpose is to choose a multiplicative factor that makes an entire suffix conservative and then select the most permissive actual threshold satisfying the target-FDR constraint.

For an actual threshold $s_i \in S_M$, define the suffix maximum ratio

$$\Gamma_M(s_i) = \max_{\substack{u \in S_M \\ u \leq s_i}} r_M(u),$$

where $u \leq s_i$ denotes $s_i$ and all thresholds at least as permissive as $s_i$. A finite positive factor makes this suffix adjusted-conservative exactly when it is at least $\Gamma_M(s_i)$. When $0 < \Gamma_M(s_i) < \infty$, this maximum is the smallest such factor. When $\Gamma_M(s_i) = 0$, every positive factor preserves suffix conservativeness.

For $\Gamma_M(s_i) < \infty$, define the lower bound on the adjusted FDR at $s_i$ as

$$H_M(s_i) = \Gamma_M(s_i)\,\beta_M(s_i).$$

If $\Gamma_M(s_i) = \infty$, then $s_i$ is excluded from the feasible set because no finite multiplicative factor can make the complete suffix conservative under the boundary conventions in Section 2.6.

For a user-specified target FDR $0 < \beta_{target} \leq 1$, define the feasible set

$$F_{\beta_{target}, M} = \{ s_i \in S_M : T^M(s_i) > 0, \Gamma_M(s_i) < \infty \text{ and } \Gamma_M(s_i) \beta_M(s_i) \leq \beta_{target} \}.$$

*Theorem statement.*

Assume that $F_{\beta_{target}, M} \neq \varnothing$. Choose the most permissive feasible actual threshold

$$s_{out} = \min F_{\beta_{target}, M},$$

where the minimum is taken in score value, so that $s_{out}$ is the most permissive feasible threshold. Define

$$\gamma_{out} = \begin{cases} \Gamma_M(s_{out}), & \Gamma_M(s_{out}) > 0, \\ \dfrac{\beta_{target}}{1 + \beta_M(s_{out})}, & \Gamma_M(s_{out}) = 0. \end{cases}$$

The zero-ratio branch selects a positive factor for an error-free suffix and satisfies the target-FDR constraint at the selected threshold.

Then the following properties hold.

First, the adjusted estimate is conservative throughout the suffix beginning at $s_{out}$:

$$FDP_M(u) \leq \beta_{adj,M}(u; \gamma_{out}) = \gamma_{out} \beta_M(u), u \in S_M, u \leq s_{out}.$$

Second, the selected threshold satisfies reference-relative observed-FDP control at the target FDR level:

$$FDP_M(s_{out}) \leq \beta_{adj,M}(s_{out}; \gamma_{out}) \leq \beta_{target}.$$

Third, $s_{out}$ is the most permissive actual threshold with $T^M(s) > 0$ among all thresholds that can satisfy both suffix-wise adjusted conservativeness and target-level adjusted FDR control using a finite positive multiplicative adjustment factor.

*Proof.*

We first prove suffix-wise adjusted conservativeness. By definition,

$$\Gamma_M(s_{out}) = \max_{\substack{u \in S_M \\ u \leq s_{out}}} r_M(u).$$

Therefore, for every threshold $u$ in the suffix beginning at $s_{out}$,

$$r_M(u) \leq \Gamma_M(s_{out}) \leq \gamma_{out}.$$

When $\beta_M(u) > 0$, the definition of $r_M(u)$ gives

$$\frac{FDP_M(u)}{\beta_M(u)} \leq \gamma_{out}.$$

Multiplying by $\beta_M(u)$ yields

$$FDP_M(u) \leq \gamma_{out} \beta_M(u) = \beta_{adj,M}(u; \gamma_{out}).$$

If $\beta_M(u) = 0$ and $FDP_M(u) = 0$, the inequality holds by the boundary convention in Section 2.6. If $\beta_M(u) = 0$ and $FDP_M(u) > 0$, then $r_M(u) = \infty$, and any suffix containing $u$ has an infinite suffix maximum. Such a threshold is therefore excluded from the feasible set. Hence every threshold in the output suffix is adjusted-conservative.

Next, because $s_{out} \in F_{\beta_{target},M}$,

$$\Gamma_M(s_{out})\beta_M(s_{out}) \leq \beta_{target}.$$

If $\Gamma_M(s_{out}) > 0$, then $\gamma_{out} = \Gamma_M(s_{out})$, giving

$$\beta_{adj,M}(s_{out};\gamma_{out}) = \gamma_{out}\beta_M(s_{out}) \leq \beta_{target}.$$

If $\Gamma_M(s_{out}) = 0$, the positive-factor branch instead gives

$$\gamma_{out}\beta_M(s_{out}) = \frac{\beta_{target}\beta_M(s_{out})}{1+\beta_M(s_{out})} \leq \beta_{target}.$$

In either case, suffix-wise adjusted conservativeness at $u = s_{out}$ gives

$$FDP_M(s_{out}) \leq \beta_{adj,M}(s_{out};\gamma_{out}).$$

Combining these inequalities yields

$$FDP_M(s_{out}) \leq \beta_{adj,M}(s_{out};\gamma_{out}) \leq \beta_{target}.$$

Thus, the selected threshold satisfies reference-relative observed-FDP control at the target FDR level.

It remains to prove retention optimality. Suppose that another actual threshold $s'$ with $T^M(s') > 0$ and another finite positive multiplicative factor $\gamma'$ satisfy the same two requirements: adjusted conservativeness throughout the suffix beginning at $s'$, and target-level adjusted FDR control at $s'$. Thus,

$$FDP_M(u) \leq \gamma' \beta_M(u), u \in S_M, u \leq s',$$

and

$$\gamma' \beta_M(s') \leq \beta_{target}.$$

At every threshold in this suffix with $\beta_M(u) > 0$, the first condition implies

$$r_M(u) = \frac{FDP_M(u)}{\beta_M(u)} \leq \gamma'.$$

The zero-boundary cases follow from the conventions in Section 2.6. Taking the maximum over the suffix gives

$$\Gamma_M(s') \leq \gamma'.$$

Therefore,

$$\Gamma_M(s')\beta_M(s') \leq \gamma'\beta_M(s') \leq \beta_{target}.$$

Hence $s' \in F_{\beta_{target},M}$. Because $s_{out}$ is defined as the most permissive threshold in this feasible set, $s'$ cannot be more permissive than $s_{out}$. Relaxing a threshold cannot decrease the number of accepted target-side matches. Therefore, $s_{out}$ maximizes target-side retention among all actual thresholds satisfying the two stated requirements.

This completes the proof.

*Interpretation.*

The suffix adjustment factor calibrates a fixed ranking; it does not change the ranking itself. If $\gamma_{out} < 1$, the unadjusted curve is more conservative than required over the selected suffix, and the adjustment compresses the estimated FDR values while preserving suffix conservativeness. If $\gamma_{out}=1$, no scaling is needed. If $\gamma_{out}>1$, the unadjusted estimate is optimistic somewhere in the suffix and must be expanded to restore conservative control. The theorem applies to any fixed ranking, including the strict, relaxed and combined rankings.

##### 3.5 Theorem 4: Interval conservativeness and interval-wise retention optimality

*Interval strategy.*

Theorem 3 enforces conservativeness from the selected output threshold to the most permissive endpoint. This full-suffix requirement can be unnecessarily restrictive when the FDR-FDP curve is non-monotone or when the distant permissive tail is noisy or overly constraining. The interval strategy instead uses a pre-specified local target operating interval and guarantees conservativeness only within that interval.

Specify the interval-construction rule and tolerance, apply this rule to the reference-labelled calibration data to determine the interval, and then hold that interval fixed while calculating the adjustment factor and selecting the final output threshold.

*Target operating interval.*

Given a target FDR $0<\beta_{target}\leq 1$ and tolerance $\varepsilon>0$, use the specified rule to choose a non-empty contiguous finite interval of actual thresholds

$$I_{target}\subseteq S_M$$

around the target operating region in the reference-labelled calibration set. For example, the stringent and permissive endpoints may be selected as thresholds whose observed FDP values are closest to the tolerance boundaries

$$\beta_{target}(1-\varepsilon)\ \text{and}\ \beta_{target}(1+\varepsilon),$$

subject to the requirement that the selected thresholds form one contiguous interval in the ordered threshold set.

If no non-empty interval can be formed, the calibration data do not support interval calibration around the target operating region. In that case, the tolerance can be enlarged, additional calibration data can be used, or a stricter scoring scheme can be selected.

For a non-empty target interval, use its maximum ratio as the adjustment factor when this maximum is positive. If all ratios are zero, choose a positive factor that satisfies the target constraint throughout the interval:

$$\gamma_I=\begin{cases}\max\limits_{u\in I_{target}} r_M(u), & \max\limits_{u\in I_{target}} r_M(u)>0,\\ \dfrac{\beta_{target}}{1+\max\limits_{u\in I_{target}}\beta_M(u)}, & \max\limits_{u\in I_{target}} r_M(u)=0.\end{cases}$$

For either branch, the adjusted estimate is

$$\beta_{adj,M}(s;\gamma_I)=\gamma_I\beta_M(s).$$

If $\gamma_I=\infty$, no finite adjustment factor can make the entire interval conservative. The theorem therefore assumes $\gamma_I<\infty$.

Define the interval-feasible set

$$F_{I,\beta_{target}}=\{s\in I_{target}:T^M(s)>0\ \text{and}\ \gamma_I\beta_M(s)\leq\beta_{target}\}.$$

*Theorem statement.*

Assume that $I_{target}$ is non-empty and fixed as specified above, that $0 < \gamma_I < \infty$, and that

$$F_{I,\beta_{target}} \neq \varnothing .$$

Define the final interval output threshold as the most permissive actual threshold in this feasible set:

$$s_{final} = \min F_{I,\beta_{target}},$$

where the minimum is taken in score value.

Then the following properties hold.

First, the adjusted estimate is conservative throughout the pre-specified interval:

$$FDP_M(u) \leq \beta_{adj,M}(u;\gamma_I), u \in I_{target}.$$

Second, the selected threshold satisfies reference-relative observed-FDP control at the target FDR level:

$$FDP_M(s_{final}) \leq \beta_{adj,M}(s_{final};\gamma_I) \leq \beta_{target}.$$

Third, within the fixed interval, $s_{final}$ is the most permissive actual threshold with $T^M(s) > 0$ that can satisfy interval-wise adjusted conservativeness and the target-FDR constraint using a finite positive adjustment factor. It therefore maximizes accepted target-side matches among interval-feasible thresholds. The theorem makes no conservativeness claim for thresholds outside $I_{target}$.

*Proof.*

We first prove interval-wise conservativeness. By construction,

$$\max_{u \in I_{target}} r_M(u) \leq \gamma_I .$$

Therefore, for every $u \in I_{target}$,

$$r_M(u) \leq \gamma_I .$$

When $\beta_M(u) > 0$, the definition of $r_M(u)$ gives

$$\frac{FDP_M(u)}{\beta_M(u)} = r_M(u) \leq \gamma_I .$$

Multiplying by $\beta_M(u)$ yields

$$FDP_M(u) \leq \gamma_I \beta_M(u) = \beta_{adj,M}(u;\gamma_I).$$

If $\beta_M(u) = 0$ and $FDP_M(u) = 0$, the inequality holds by the boundary convention in Section 2.6. If $\beta_M(u) = 0$ and $FDP_M(u) > 0$, then $r_M(u) = \infty$, which would imply $\gamma_I = \infty$, contradicting the theorem assumption. Therefore, every threshold in the pre-specified interval is adjusted-conservative.

Because

$$s_{final} \in F_{I,\beta_{target}},$$

we have

$$\gamma_I \beta_M(s_{final}) \leq \beta_{target}.$$

Using the definition of the adjusted estimate,

$$\beta_{adj,M}(s_{final};\gamma_I)=\gamma_I\beta_M(s_{final})\le\beta_{target}.$$

Interval-wise conservativeness at $s_{final}$ gives

$$FDP_M(s_{final})\le\beta_{adj,M}(s_{final};\gamma_I).$$

Combining the two inequalities gives

$$FDP_M(s_{final})\le\beta_{adj,M}(s_{final};\gamma_I)\le\beta_{target}.$$

Thus, the selected threshold satisfies reference-relative observed-FDP control at the target FDR level.

It remains to prove interval-wise retention optimality. First, suppose that another threshold $s'\in I_{target}$ with $T^M(s')>0$ satisfies the target-level adjusted FDR constraint under the same interval factor:

$$\gamma_I\beta_M(s')\le\beta_{target}.$$

Then $s'\in F_{I,\beta_{target}}$. Because $s_{final}$ is defined as the most permissive threshold in this feasible set, $s'$ cannot be more permissive than $s_{final}$. Therefore, no threshold in the pre-specified interval using the factor $\gamma_I$ can retain more target-side matches while satisfying the target-FDR constraint.

If the maximum ratio is zero, all observed FDP values in the interval are zero, and the positive-factor branch satisfies

$$\gamma_I\beta_M(u)=\frac{\beta_{target}\beta_M(u)}{1+\max_{v\in I_{target}}\beta_M(v)}\le\beta_{target},u\in I_{target}.$$

Every threshold in the interval with $T^M(u)>0$ is therefore feasible. Selecting the most permissive one maximizes target retention over the entire eligible interval.

For a positive maximum ratio, consider any alternative finite positive adjustment factor $\gamma'$ that also guarantees interval-wise conservativeness over $I_{target}$. For every $u\in I_{target}$,

$$FDP_M(u)\le\gamma'\beta_M(u).$$

For thresholds with $\beta_M(u)>0$, this is equivalent to

$$r_M(u)\le\gamma'.$$

The zero-boundary cases follow from the conventions in Section 2.6. Taking the maximum over the interval gives

$$\gamma_I=\max_{u\in I_{target}} r_M(u)\le\gamma'.$$

In this positive-ratio case, any finite positive adjustment factor that guarantees interval-wise conservativeness must be at least $\gamma_I$.

If a more permissive threshold $s'\in I_{target}$ with $T^M(s')>0$ satisfied target-level control under such an alternative factor, then

$$\gamma'\beta_M(s')\le\beta_{target}.$$

Since $\gamma_I \leq \gamma'$,

$$\gamma_I \beta_M(s') \leq \gamma' \beta_M(s') \leq \beta_{target}.$$

Therefore,

$$s' \in F_{I,\beta_{target}},$$

which contradicts the definition of $s_{final}$ as the most permissive member of this set. Hence, $s_{final}$ is the most permissive threshold within the pre-specified interval satisfying interval-wise adjusted conservativeness and target-level adjusted FDR control.

This completes the proof.

*Interpretation.*

The interval theorem is a local calibration strategy. It intentionally guarantees conservativeness only within the pre-specified target interval. This is useful when behaviour in the distant permissive tail would make full-suffix calibration unnecessarily restrictive or unstable.

The interval is constructed from the calibration data using the specified rule and remains fixed during adjustment-factor calculation and output-threshold selection. For a given suffix beginning at $s_i$, its maximum ratio equals $\Gamma_M(s_i)$ in Theorem 3. Theorem 3 evaluates a separate suffix factor for each candidate output threshold, whereas Theorem 4 uses one factor over its fixed interval.

#### 4. Conclusion

Lemma 1 and Theorems 1–4 establish the theoretical basis of CCT by connecting score ranking, counterpart counting and FDR calibration. For strict scores, Lemma 1 establishes the expected-count correspondence between incorrect targets and their counterparts under the stated threshold-passing symmetry assumption, and characterizes the cumulative-count conditions under which the additional contribution from $C_t$ makes the estimate conservative. For the relaxed ranking, complete pairing and a finite threshold set ensure endpoint conservativeness and the existence of a local conservative starting point (Theorem 1).

The combined score integrates information from the strict and relaxed scores. Under the conditions of Theorem 2, a combined ranking sufficiently close to the strict-score endpoint reaches a conservative suffix no later than the relaxed ranking in accepted-target-count coordinates. This connects the strict ranking's conservativeness to the location of the combined ranking's conservative onset.

Once the ranking is fixed, Theorems 3 and 4 provide calibration over a full suffix and a specified target interval, respectively. When the corresponding feasible set is nonempty, each procedure selects the most permissive threshold satisfying conservativeness over its prescribed range and the target-FDR constraint. The selected threshold therefore yields a reference-relative observed FDP at or below the target level and maximizes target retention among the feasible thresholds within that fixed ranking. These results provide a theoretical basis for counterpart-based error estimation, calibration and threshold selection in de novo peptide quality control.

### Supplementary Note 2: π-NovoQC workflow and feature definitions

π-NovoQC takes raw MS/MS spectra, database-search identifications and a reference protein database as input. Database-search identifications provide reference labels for fitting the CCT scoring models and calibrating the FDR estimates, while the reference protein database is used for peptide mapping and protein inference. The workflow comprises four modules: (i) de novo peptide prediction and construction of paired target and counterpart QC features; (ii) CCT quality control at the peptide-spectrum-match (PSM) and peptide levels; (iii) peptide mapping and protein inference with separate target-decoy FDR control; and (iv) MS1-based peptide and protein quantification. Accepted de novo peptide evidence is integrated with database-search results for protein quantification. The reported results also include candidate peptides without a qualifying match to the reference protein database.

#### Feature-based construction of paired target-counterpart representations from decoder cross-attention

π-xNovo uses acquisition-specific Transformer-based sequencing models [3] for DDA and DIA data. The spectrum encoder represents the experimental MS/MS peaks, and the peptide decoder predicts a candidate sequence from the C terminus towards the N terminus. Decoder cross-attention links sequence positions to spectral peaks and is used to construct paired target and counterpart QC features for the same candidate peptide and experimental spectrum.

For a candidate peptide of length $L$, row 0 of the decoder cross-attention matrix corresponds to the precursor query, rows 1 to $L$ correspond to the input residues in decoding order, and the final row corresponds to the stop token. Target features are extracted from rows $0{:}L$, and counterpart features from rows $1{:}1{+}L$, using zero-based, end-exclusive indexing (Fig. 1b). These windows represent adjacent decoder states with different sequence prefixes while attending to the same experimental spectrum. The same feature-extraction procedure is applied separately to the two windows. Attention-derived features are computed from their respective windows, while spectrum-peptide matching quantities, including fragment coverage and explained intensity, are shared between the paired feature vectors. DDA processing additionally includes precursor-mass error, giving 33 scalar features; DIA processing uses the corresponding 32 features without the precursor-error scalar. The feature definitions and their roles are summarized below and in Supplementary Table 2.

Residue-confidence features combine decoder cross-attention with the intensities and fragment-ion assignments of the experimental spectrum. For each position in the predicted peptide, high-attention spectral peaks are evaluated for support from the corresponding theoretical fragments. The resulting position-wise confidence scores are summarized across the peptide.

The residue-confidence summaries comprise the mean, variance, minimum and maximum, together with the numbers of low- and high-confidence residues. The anchor_ratio feature measures the proportion of high-confidence residues. Supplementary Table 2 lists the scalar features.

Spectrum-peptide agreement is represented by fragment coverage, explained intensity and a global QC score. Observed peaks are matched to theoretical fragments from the predicted peptide. Fragment coverage measures the proportion of annotated spectral input positions, and explained intensity measures the fraction of total spectral intensity assigned to matching fragments. DDA analyses additionally include precursor-mass error.

Attention-derived features describe how spectral evidence is distributed over the selected decoder-query positions. They include attention entropy and concentration, the mass offset of the highest-attention peak, the displacement of the attention-weighted mass centre, and the mass dispersion around that centre. Each position-wise statistic is summarized by its mean, standard deviation, minimum and maximum across the peptide.

Two further features summarize consistency across positions: the mean cosine similarity between adjacent attention vectors and the Spearman correlation between decoding position and the mass-to-charge ratio of the highest-attention peak. The same feature definitions are applied to target and counterpart representations using their respective attention-row selections.

These summaries were concatenated to form a PSM feature vector capturing residue-level evidence and overall spectrum-peptide agreement. The shared feature set contained seven residue-confidence summaries, including anchor_ratio; three overall agreement features (global_qc_score, fragment_coverage and explained_intensity); 20 attention-statistic summaries; and the two position-consistency features described above, giving 32 scalars. DDA processing additionally included precursor_error_ppm, giving 33 scalars. The same extraction procedure was applied to both members of each target-counterpart pair, with attention-derived quantities calculated from their respective windows.

#### Feature groups and ablation configurations

For the ablation experiments in **Supplementary Figs. 9 and 10, t**he scalar features defined above were partitioned into five nonoverlapping groups. These experimental groups differ from the descriptive grouping used for feature extraction above; in particular, anchor_ratio was assigned to spectrum support.

Amino-acid confidence (6 features): aa_conf_mean, aa_conf_var, aa_conf_min, aa_conf_max, num_low_conf and num_high_conf.

Spectrum support (5 DDA features; 4 DIA features): global_qc_score, fragment_coverage, explained_intensity, anchor_ratio and, for DDA only, precursor_error_ppm.

Attention uncertainty (8 features): the mean, standard deviation, minimum and maximum of aa_entropy and aa_concentration.

Attention geometry (12 features): the mean, standard deviation, minimum and maximum of aa_offset, aa_center_drift and aa_span.

Cross-view consistency (2 features): avg_attention_similarity and fragment_alignment_consistency.

The eight DDA configurations were All 33; omission of amino-acid confidence (27 features), spectrum support (28), attention uncertainty (25), attention geometry (21) or cross-view consistency (31); Compact 11; and Compact 19. Compact 11 combines amino-acid confidence and spectrum support. Compact 19 adds the attention-uncertainty group to Compact 11.

The DIA configurations use the same groups but omit precursor_error_ppm throughout: All 32; the five respective single-group omissions (26, 28, 24, 20 and 30 features); Compact 10; and Compact 18. Compact 10 combines amino-acid confidence and spectrum support; Compact 18 additionally includes attention uncertainty. Every configuration constructs its peptide-level population from its own PSM-filtered results.

#### Dual-level CCT control at the PSM and peptide levels

π-NovoQC applies CCT at the PSM and peptide levels using the target and counterpart features described above. Database-search identifications provide reference labels for fitting the scoring models and calculating the adjustment factor $\gamma$. The reference set for each analysis comprises the specified sample-related database-search PSMs, including auxiliary references where applicable. PSM-level and peptide-level scoring models and adjustment factors are fitted separately using their respective feature populations. At each level, target and counterpart features are pooled, ranked by the combined score and evaluated at a shared score threshold. The resulting cumulative counts are used to estimate FDR, and $\gamma$ multiplicatively adjusts this estimate.

#### Scoring model configuration and training

For spectra with database-search identifications, comparison of the predicted peptide with the reference peptide for the same spectrum determines the four feature-category assignments. Agreement assigns the target feature to $T_t$ and its paired counterpart feature to $C_t$; disagreement assigns them to $T_f$ and $C_f$, respectively. These assignments describe agreement with the database reference and provide the basis for defining the training labels of the two scoring models.

The strict and relaxed scoring models were implemented as feedforward multilayer perceptrons (MLPs), using ReLU activations [4], batch normalization [5] and a sigmoid output for candidate ranking. Scoring-model dropout [6] is distinct from the construction of target-counterpart features, which uses two row selections from the same cross-attention matrix.

The scoring MLPs were trained on database-reference-labelled feature vectors using binary cross-entropy and AdamW [7]. Label-stratified training and validation subsets were used to fit and select the scoring models; feature standardization was fitted on the training subset. PSM-level and peptide-level models were trained separately.

The strict model labelled $T_t$ and $C_t$ positive (1), and $T_f$ and $C_f$ negative (0), producing $s_{strict}$. The relaxed model labelled only $T_t$ positive and $T_f$, $C_t$ and $C_f$ negative, producing $s_{relaxed}$. The models share feature definitions but assign different labels to $C_t$.

For a feature vector $x$, the combined score was calculated from the two model outputs as

$$s_{combined}(x) = \alpha\, s_{strict}(x) + (1-\alpha)\, s_{relaxed}(x), \alpha \in [0,1].$$

The default $\alpha = 0.5$ averages the strict and relaxed scores. Target and counterpart counts accumulated along this combined ranking support FDR estimation and calibration under the conditions in **Supplementary Note 1**.

#### Adjustment factor gamma (γ)

We sorted the mixed score queue in descending order of the combined score $s_{combined}$. For a score threshold $s_i$ with $T(s_i) > 0$, the unadjusted FDR estimate was defined as

$$\beta(s_i) = \frac{C(s_i)}{T(s_i)}, T(s_i) > 0.$$

where $T(s_i)$ is the number of target PSMs or peptides with scores $\geq s_i$, and $C(s_i)$ is the number of counterpart-side PSMs or peptides with scores $\geq s_i$. These counts include all passing targets and counterparts, respectively, regardless of their scoring-model training labels.

We calibrated this estimate at the user-specified target FDR, $\beta_{target}$, using database-search identifications as references. In the reference-labelled population, the number of incorrect targets passing $s_i$, $T_f(s_i)$, gives the observed FDP, $T_f(s_i) / T(s_i)$. A positive multiplicative adjustment factor $\gamma$ was calculated from the relationship between the estimated FDR and observed FDP, yielding

$$\beta_\gamma(s_i) = \gamma\beta(s_i) = \gamma\frac{C(s_i)}{T(s_i)}.$$

Suffix calibration makes the adjusted estimate conservative from the selected threshold through all more permissive thresholds. Interval calibration applies within a target operating interval defined by a specified rule and held fixed during factor calculation and threshold selection (**Supplementary Note 1**, Theorems 3 and 4). Each procedure selects the most permissive feasible threshold satisfying $\beta_\gamma(s_i) \le \beta_{target}$ within its calibration range.

The factor $\gamma$ rescales FDR estimates without changing scores, ranking or counts at a fixed threshold, but can change the selected cutoff and retained set. After fitting the models and $\gamma$, candidate selection uses scores and counts without requiring candidate correctness labels.

#### PSM-level FDR control (CCT-PSM QC)

The PSM-level workflow was as follows:

1. Model fitting and calibration. We fitted strict and relaxed models to PSM target/counterpart features with database-reference labels, ranked their $s_{combined}$ scores and calculated the PSM-level adjustment factor $\gamma$ at $\beta_{target}$.
2. FDR estimation. We scored all candidate target/counterpart PSM features from the same raw file and ranked the pooled entries by $s_{combined}$. For each threshold $s_i$ with $T(s_i) > 0$, cumulative passing counts gave:

$$\beta_\gamma(s_i) = \gamma\frac{C(s_i)}{T(s_i)}.$$

3. PSM selection. We chose the most permissive threshold $s_\gamma$ satisfying $\beta_\gamma(s_i) \le \beta_{target}$ and retained target PSMs with scores $\ge s_\gamma$.
4. Paired-feature recovery. For each pair with either side passing $s_\gamma$, we recovered both target and counterpart entries from the full feature table for sequence collapsing and peptide-level QC.

#### Peptide-level FDR control (CCT-Peptide QC)

Peptide-level QC used sequence representatives from the recovered paired-feature population:

1. Sequence collapsing. Within each sample or predefined pooled experiment, we grouped entries by full modified peptide sequence and retained the highest combined-score PSM and its features as the sequence representative.
2. Model fitting and calibration. Representatives with database-reference labels trained new strict and relaxed peptide models. Their combined scores and the specified target FDR determined a separate peptide-level adjustment factor $\gamma$.
3. Peptide selection. We scored and ranked the pooled target/counterpart representatives, calculated adjusted FDR from their cumulative counts and peptide-level $\gamma$, and selected the most permissive threshold satisfying the peptide target FDR. Target sequences passing this threshold were retained. The default target FDR is 5% at each QC level.
4. **Output for downstream analysis.** Among the accepted target peptides, those originating from spectra without database-search identifications were carried forward to protein inference and protein-level FDR control.

#### Protein-level FDR control

Each target FASTA sequence was reversed and assigned a decoy-prefixed identifier and description. Target and decoy sequences were deduplicated separately, then concatenated for peptide mapping and protein inference.

Peptides originating from spectra without database-search identifications and accepted by PSM- and peptide-level CCT QC were processed as follows:

1. Sequence export. We removed modification mass-shift annotations from accepted peptides and exported amino acid sequences in FASTA format, retaining spectrum-index associations.
2. Protein mapping. BLAST [8] searched the target-decoy database. E-value, alignment coverage and permitted substitutions filtered peptide-protein assignments and alignment scores.
3. Protein inference. PyProteinInference [9] used peptide-protein assignments, spectrum identifiers and alignment scores for parsimonious target-decoy inference, followed by filtering at protein q ≤0.01.
4. Output integration. Peptide sequences linked inference results to supporting PSM entries; duplicate protein entries were removed.

For subsequent quantification, the accepted de novo peptide evidence was merged with database-search evidence, and protein groups were inferred again from the combined evidence, without an additional protein-FDR filter at that stage.

#### Peptide quantification

The peptide quantification procedure was as follows:

1. Precursor m/z calculation. We combined protein-inference output, PSM metadata and mzML MS1/MS/MS data for de novo and database-search peptides. Peptide sequences and modifications supplied masses through pyteomics mass.calculate_mass [10]. For charge $z$, theoretical m/z was

$$\frac{m}{z}_{theoretical} = \frac{M + z m_p}{z},$$

   where $M$ is the neutral peptide mass, including modifications, and $m_p$ is the proton mass. The calculated values were added to the sample-level PSM table.
2. Isotope XICs. MS1 peaks were extracted within the specified ppm tolerance around monoisotopic m/z and successive isotope positions, separated by approximately $1.003355/z$. Intensities at the same retention time (RT) were summed for each isotope to form its XIC.
3. RT window. After Savitzky-Golay smoothing, the single-isotope XIC maximum defined the apex. The window extended in both directions until intensity fell to a specified fraction of peak height (default, 10%), defining boundaries $RT_{min}$ and $RT_{max}$ and apex RT $RT_{peak}$.
4. Isotope selection. pyteomics mass.Composition supplied peptide elemental composition and theoretical isotope distributions. Pearson correlation, dot-product angle and residual sum of squares measured observed/theoretical abundance agreement. Isotope traces were retained by their correlation with the monoisotopic XIC within the selected RT window.

5. Intensity and quality metrics. Peptide intensity sums retained-isotope peak areas within the RT window. Recorded features include MS1 scan and peak counts, retained-isotope count, original and isotope-weighted corrected precursor m/z, RT boundaries/apex/width, isotope-distribution agreement, XIC correlations and signal-to-noise ratio (main peak intensity/mean noise intensity). Predefined thresholds assign an A/B/C quality grade.

6. Output. Processing the predefined peptide list produced a table of intensities, isotope-consistency metrics, quality grades and quantitative features for protein quantification.

#### Protein quantification

Protein quantification integrated DB and de novo peptide intensities with protein-inference results:

1. Peptide merging. Within a sample, the maximum intensity represented a peptide supported by multiple PSMs. Across samples, duplicate peptide-intensity entries were summed. Sequence-based merging retained source annotations (db or denovo) and prioritized DB peptides during deduplication.
2. Joint inference. PyProteinInference used peptide quantitative values without additional protein filtering. Adding de novo peptides can change the peptide-protein graph for DB identifications.
3. Theoretical peptide counts. Proteins were digested in silico using predefined rules for the experimental enzyme (for example, trypsin, GluC, LysC, AspN or chymotrypsin). The estimated observable-peptide count supplied coverage and iBAQ denominators.
4. MS/MS weights. DB acquisition counts and de novo observations per peptide in PSM-FDR results were summed to weight mean intensities.
5. After merging peptide intensities with inference results, per-protein, per-sample metrics were:
    - **Number of peptides:** the number of observed peptides.
    - **Protein coverage:** number of observed peptides / total theoretical peptide count.
    - **iBAQ (intensity-based absolute quantification):** protein score (PyProteinInference score) / total theoretical peptide count.
    - **TopN mean intensity:** for each protein in each sample, the N peptides with the highest intensities (default N = 3) were selected and averaged.
    - **Weighted mean intensity:** peptide intensities were averaged using total MS/MS counts as weights:

    $$\text{Weighted Average} = \frac{\sum_i Area_i \times MS2_{\text{total},i}}{\sum_i MS2_{\text{total},i}}$$

    where $Area_i$ is the area of the ith peptide.
6. Quantification results were computed under two schemes (DB-only and DB + de novo):
    - **DB-only quantification:** protein inference and quantification were performed using only DB peptides.
    - DB + de novo quantification: infer and quantify proteins after merging both peptide sources; compare with DB-only results to assess added coverage and quantitative changes.

The output reports peptide and theoretical-peptide counts, coverage, iBAQ, TopN and weighted mean intensities, and total MS/MS count for each protein and sample.

### Supplementary Note 3: Datasets and Preprocessing

#### Datasets

Nine-species dataset: We downloaded the nine-species dataset from Tran et al. [11]. It contains ~1.52 million peptide-spectrum matches (PSMs) identified by database search at 1% false discovery rate (FDR) from tryptic digests acquired with higher-energy collisional dissociation (HCD) on a Thermo Scientific Q-Exactive mass spectrometer. Carbamidomethylation of cysteine (C) was specified as a fixed modification, and variable modifications included oxidation of methionine (M) and deamidation of asparagine (N) and glutamine (Q). We did not distinguish isoleucine (I) from leucine (L). We evaluated model performance using leave-one-species-out cross-validation: in each fold, one species was held out for testing, and the remaining species were used for training.

Sequencers used only nine-species training data, without MassIVE-KB data or pretrained weights. Downstream QC used an 8:2 scoring-MLP training/validation split of database-reference labels within each held-out species, then γ calibration and evaluation on the full valid population, including training PSMs. Thus, leave-one-species-out testing applies to sequencing; downstream QC is sample-internal. In the human fold, π-xNovo, π-HelixNovo and Casanovo retained 38,456, 17,816 and 20,169 correct PSMs and 11,376, 6,429 and 5,796 correct peptides, respectively.

GraphNovo dataset [12]: human HeLa and mouse cerebellum data supplied ~1,659,763 training PSMs; *A. thaliana*, *C. elegans* and *E. coli* supplied 12,500 test PSMs each. Tryptic spectra were acquired by HCD on a Thermo Q-Exactive and identified by database search at 1% FDR. Cysteine carbamidomethylation was fixed and methionine oxidation variable. Isoleucine and leucine were not distinguished.

MassIVE-KB dataset: We downloaded the MassIVE-KB dataset from Wang et al. [13], which contains ~30 million high-quality PSMs filtered at a highly stringent FDR (reported as ~0%); for each unique peptide-charge combination, at most 100 PSMs were retained. Most spectra were generated from tryptic digests. The fixed modification was carbamidomethylation of cysteine (C). We considered seven variable modifications: oxidation of methionine (M), deamidation of asparagine (N) and glutamine (Q), N-terminal acetylation, N-terminal carbonylation, N-terminal NH3 loss, and the combination of N-terminal carbonylation and NH3 loss. *A. thaliana*, *C. elegans*, and *E. coli* were used as external test species for the π-xNovo model. The numbers of overlapping PSMs (peptides) between the test sets and this dataset were 31 (22), 130 (98), and 12 (6), respectively. The resulting average test-set overlap rate was ≤0.42%, which we considered negligible for our conclusions.

DHP dataset: We downloaded 2,460 mzML files from the Deep Human Proteome (DHP) project described by Sinitcyn et al. [14], comprising ~161 million MS/MS spectra. Spectra were acquired from six human cell lines (hES1, HeLa S3, HepG2, GM12878, K562, and HUVEC) digested with six parallel proteases (LysC, LysN, AspN, chymotrypsin, GluC, and trypsin) and fragmented using HCD, collision-induced dissociation (CID) or electron-transfer dissociation (ETD) on Thermo Orbitrap Fusion or Thermo Orbitrap Lumos instruments. EThcD spectra were treated as ETD. The original study searched 2,491 raw files against four protein sequence databases (UniProt canonical 2017_02; UP000005640_9606; UniProt isoforms UP000005640_9606_additional; Ensembl canonical v86 GRCh38.pep.all; and Ensembl isoforms GRCh38.pep.abinitio) using MaxQuant (v1.5.7.5) [15, 16]. We extracted 16,746,362 HCD PSMs, 2,389,865 ETD PSMs, and 1,482,573 EThcD PSMs from the corresponding msms.txt files provided for each cell line. Because fragmentation patterns differ, we trained separate π-xNovo models for HCD and ETD. For each cell line-enzyme combination, we randomly sampled 10,000 spectra as the test set; the HCD and ETD test sets contained 360,000 and 300,000 PSMs, respectively (trypsin-generated spectra were HCD). In addition to the PSMs described above, we extracted an additional 174,945 PSMs from msms.txt in txt.zip; these do not have corresponding downloadable mzML files. In total, we obtained 20,793,745 PSMs from the project. The specified fixed modification was carbamidomethylation of cysteine (C), and the variable modifications were oxidation of methionine (M) and protein N-terminal acetylation. After removing redundant sequences across the four protein sequence databases, we constructed a new non-redundant protein sequence database containing 142,459 protein sequences. This database was used as input to the π-NovoQC platform for in-database peptide analysis and quantms [17] re-search. We also obtained variant protein databases (proteins.fa) for six cell lines from variationExtraction.zip, which were used for single amino-acid polymorphism (SAP) analysis.

Astral DDA and DIA datasets: We downloaded narrow-window Astral DDA and DIA comparison data (PXD046453) from Guzman et al. [18]. Samples were trypsin-digested and analyzed by LC-MS/MS on an Orbitrap Astral instrument using HCD with a 2 Th isolation window. Astral DDA (n=3) and DIA (n=6) data for 1,000 ng tryptic HEK293 peptides were used to validate π-NovoQC. The DIA experiment included two sets of three replicates acquired with maximum injection times of 2.5 ms and 3.5 ms. For the DDA experiment, we collected PSMs from 15 samples for training the $\pi$-xNovo model, including the three replicates paired with the DIA experiment (samples 1-3 contained 71,017, 69,843, and 69,481 PSMs). To maintain a consistent bioinformatics workflow, the original study analyzed both DIA and DDA data using DIA-NN 1.8.1 [19] against a human protein reference database (UniProt 2022; 20,598 sequences). The fixed modification was carbamidomethylation of cysteine (C), and variable modifications included oxidation of methionine (M) and N-terminal acetylation.

Orbitrap DIA dataset: We downloaded 194 wide-window DIA raw files for clear-cell renal cell carcinoma (PDC000200) from Clark et al. [20]. Samples were trypsin-digested and analyzed by LC-MS/MS on an Orbitrap Fusion Lumos instrument using HCD with variable-width isolation windows. In Sanders et al. [21], the dataset was searched against a human protein reference database using MSFragger-DIA v21.1 with the standard DIA_Speclib_Quant workflow and default settings [22]. The fixed modification was carbamidomethylation of cysteine (C).

TimsTOF DDA dataset: We collected in-house 100 HEK293T samples and 100 mouse tissue samples used for routine quality control on a timsTOF mass spectrometer. We performed database searches with PEAKS-Online (version 12) for human and mouse samples and filtered identifications at 1% FDR, yielding 32,018,308 PSMs. The fixed modification was carbamidomethylation of cysteine (C), and the variable modification was oxidation of methionine (M).

Dong-Ecoli-QE isotope-labeled dataset: We downloaded the Dong-Ecoli-QE dataset (PXD008782) from Chi et al. [23]. The dataset comprises metabolically isotope-labeled *E. coli* grown in media containing unlabeled, $^{15}$N-labeled, and $^{13}$C-labeled sources mixed 1:1:1. The dataset contains five raw files (202,452 MS/MS spectra) corresponding to labeling conditions of 10/30/60/150/1000 mM. Data were acquired on an Orbitrap Q-Exactive instrument after tryptic digestion. The fixed modification was carbamidomethylation of cysteine (C). The original study performed a database search using Open-pFind (v1.0) [23].

#### Preprocessing for DDA datasets

For the nine-species, GraphNovo, MassIVE-KB, DHP, timsTOF DDA, and Dong-Ecoli-QE isotope-labeled datasets, the original studies provided raw spectra and database-search results with a unique peptide assignment for each MS/MS spectrum. We linked peptides to spectra using spectrum identifiers. To standardize model inputs and improve computational efficiency, we applied the following preprocessing steps to all MS/MS spectra, including spectra without database identifications:

1. **m/z range filtering:** discard peaks with m/z outside 50-2,500.

2. **Intensity thresholding:** discard peaks with intensity below 10% of the maximum peak intensity within the spectrum.

3. **Peak list truncation:** retain up to the 150 most intense peaks per spectrum.

4. **Intensity normalization:** rescale retained peak intensities within each spectrum to a common scale.

For the Astral DDA dataset, we first converted raw files to mzML using msconvert (ProteoWizard) [24]. To enforce a unique peptide assignment for each MS/MS spectrum, we removed co-eluting peptides from the database-search outputs before peptide-spectrum matching. Training data were generated using the same preprocessing pipeline. The Astral DDA dataset contained 580,736 PSMs, including 27,136 PSMs in the test set.

#### Preprocessing for DIA datasets

DIA database searches report run-level identifications with many-to-many peptide/spectrum relationships. Cascadia [21] combines MS1/MS/MS signals into a peptide-specific enhanced spectrum. π-xNovo instead predicts multiple peptides directly from original DIA MS/MS spectra, retaining all co-eluting database-search peptides for each spectrum.

pyteomics [10] extracted RT and isolation-window metadata from DIA mzML spectra. A database-search peptide was associated with a spectrum when its theoretical precursor m/z lay in that isolation window and its RT differed by at most 1 second (Astral) or 5 seconds (Orbitrap). Entries recorded spectrum ID, peptide, RT and precursor m/z, allowing multiple peptides per spectrum. Astral and Orbitrap yielded 782,793 and 5,990,677 PSMs, with test sets of 134,532 and 31,250. For Cascadia benchmarking, default enhanced-spectrum generation yielded 902,814 and 5,473,994 PSMs, with test sets of 161,322 and 28,864, respectively.

### Supplementary Note 4: End-to-end Validation of π-NovoQC

#### Spectral similarity calculation

AlphaPeptDeep [25] predicted fragment intensities from peptide sequence, modifications and charge: b/y ions for HCD/CID and c/z for ETD. Each theoretical fragment m/z was matched to the nearest experimental mzML peak within 0.02 Da; unmatched fragments received zero intensity. Cosine similarity compared the aligned predicted and observed vectors.

DHP QC covered 2,458 raw files and 390,319 distinct in-database peptides. Quantification included 2,456 files and 387,854 positive-intensity peptides after excluding 2,465 zeros. Without cross-sample deduplication, spectral-similarity counts were 438,790 shared DB/de novo and 460,412 additional de novo peptides (899,202 total); precursor-error analysis used 901,941 peptide observations. Spectral similarity was also calculated for the GraphNovo *A. thaliana* test set.

#### Quantification comparison

DDA quantification compared π-NovoQC, MaxQuant [15, 16] and PANDA [26] using one raw file from each of 64 DHP conditions. PANDA input came from cell-line MaxQuant evidence.txt files: scan IDs, sequences, protein assignments, variant sites and metadata. Default settings used 20-ppm XIC tolerance, signal-to-noise threshold 2 and a 5-minute RT window. MaxQuant intensities came directly from evidence.txt. DIA comparisons used DIA-NN [19] and all six Astral runs.

#### End-to-end validation

After all three QC levels, DHP yielded 901,941 in-database peptide observations across 2,458 files without cross-file deduplication. The 899,202 observations used for cosine similarity resembled DB identifications but shifted lower; GraphNovo *A. thaliana* QC-passing predictions had higher similarities than non-passing predictions (**Supplementary Fig. 17a**). More than 99% of DHP precursor errors were within ±20 ppm, centred near zero (**Supplementary Fig. 17b**).

Across 64 DHP samples, $\log_2$-intensity differences from MaxQuant had mean bias 0.040 for π-NovoQC (median 0.073; IQR 0.000–0.099) and 0.044 for PANDA (median −0.016; IQR −0.073–0.063). Median standard deviations were 1.273 and 1.307, respectively (2.6% difference); π-NovoQC was lower in 60.9% of samples (**Supplementary Fig. 17c**).

Mean/median peptide-intensity correlations with MaxQuant were 0.869/0.886 for π-NovoQC and 0.869/0.894 for PANDA. The paired difference Δr = r_PANDA − r_π-NovoQC averaged −0.001 (95% CI −0.032 to 0.031; paired t-test P = 0.97; Wilcoxon P = 0.68); PANDA was higher in 51.6% of samples (**Supplementary Fig. 17d**). Median protein-level correlation between π-NovoQC and PANDA was 0.818 (reported bootstrap 95% CI 0.803–0.832; **Supplementary Fig. 17e**).

Across six Astral DIA runs, π-NovoQC/DIA-NN peptide intensities had Pearson r ≈0.57–0.59, $R^2$ ≈0.33–0.34 and regression slopes ≈0.57–0.59, with similar slopes and intercepts across runs (**Supplementary Fig. 18**).

#### Additional QC evaluation details for Figs. 2–5

The fixed-score weight analysis in **Supplementary Figs. 7 and 8** uses 37,321 target and 36,887 counterpart PSM entries and a common peptide population after default PSM selection. At weight 0.5 and nominal 5% FDR, it retained 24,024 correct PSMs and 16,690 correct peptides, with observed FDPs of 5.0097% and 4.1851%. Relaxed scoring retained 22,062 correct PSMs and 14,469 correct peptides, with FDPs of 4.9994% and 5.0030%. PSM AUC increased from 0.9352 to 0.9514 and peptide AUC from 0.7455 to 0.9328. This analysis is separate from the original calibrated comparison in **Fig. 2** and **Supplementary Fig. 1**.

In the DDA feature ablation (**Supplementary Fig. 9**), Compact 11 and Compact 19 retained mean fractions of 77.31% and 86.93% of All-33 reference-correct PSMs and 73.81% and 86.50% of its correct peptides. Mean peptide AUCs were 0.7264, 0.8265 and 0.9381 for Compact 11, Compact 19 and All 33. Removing attention geometry retained 88.72% of correct PSMs and 87.54% of correct peptides, with peptide AUC 0.8409. The DIA experiment comprised 48 sample-by-configuration comparisons and 96 results across the two QC levels (**Supplementary Fig. 10**).

For the sequencing-model comparison in **Fig. 4a,b and t**he nine-species analysis, training datasets and QC populations are defined in **Supplementary Note 3**. In the nine-species analysis, pooled retention–adjustment-factor correlations were approximately −0.74 at the PSM level and −0.51 at the peptide level (**Supplementary Fig. 11**).

For **Fig. 4g, o**verlap is the fraction of each replicate's peptides shared with at least one other replicate, averaged over the three replicates. The DDA, 2.5-ms DIA and 3.5-ms DIA peptide unions contained 28,026, 94,257 and 97,806 modified sequences. These sets use per-entry adjusted FDR ≤0.05. The near-threshold cumulative selection in **Fig. 4b,d,f** can differ: for 3.5-ms Rep 2, the two definitions yield 63,476 and 63,500 peptides. The held-out designation in **Fig. 4e** refers to sequencing-model training, not a downstream QC split.

In the Cascadia comparison (Fig. 4h; Supplementary Table 5), the native Cascadia feature set was evaluated alone or combined with auxiliary reference-PSM features extracted by π-xNovo-DDA from GraphNovo or by π-xNovo-DIA from Astral data. Cascadia predictions and their native features were retained in all three settings. The combined feature populations were used for QC model training and FDR calibration. The three settings retained 87, 3,110 and 14,428 reference-correct Cascadia PSMs, respectively; auxiliary reference PSMs are excluded from these counts. Relative to the native-feature setting, the latter totals were 35.75-fold and 165.84-fold higher. The corresponding 2.5-ms subtotals were 38, 1,545 and 7,422, and the 3.5-ms subtotals were 49, 1,565 and 7,006. Three native-feature runs had no usable calibration threshold and contributed zero to the totals. This comparison evaluates the combined settings; it does not isolate the effects of the reference data and the model used to extract the auxiliary features.

After sequence harmonisation on GraphNovo, representative-PSM disagreement proportions were 6.7704%, 8.6976% and 3.0409% for π-NovoQC, NovoBoard and Winnow, respectively (**Supplementary Table 10**). For native two-stage DHP QC, the 302,573 peptide-stage identifications included 292,351 reference-correct and 10,222 reference-incorrect entries (**Supplementary Table 12**).

#### Reference availability and timing in the DHP method comparison

Across the 85 evaluation raw files, MS2-spectrum counts ranged from 73,407 to 120,178 and database-identified distinct spectra from 1,860 to 24,019, giving identification fractions of 2.0662%–28.0146%. Sixty-nine files used their own references and 16 used auxiliary references (**Supplementary Figs. 14 and 15**). For the sparse-reference examples in the main Results, the fraction-11 LysN-digested HepG2 raw file had a database-identification fraction of 2.0662% and retained one reference-incorrect PSM and two reference-incorrect peptides. The fraction-5 AspN-digested GM12878 raw file had a database-identification fraction of 2.5338% and retained nine reference-incorrect PSMs and 14 reference-incorrect peptides. Both used their own database references.

Timing used 9,777 candidate PSMs from the HCD raw file for fraction 10 of AspN-digested GM12878 on one Tesla V100 32 GB GPU with eight CPU threads (**Fig. 5f**; **Supplementary Tables 14 and 15**). π-NovoQC required 416.566 seconds from spectrum reading through feature generation, model training and both QC stages; NovoBoard required 313.986 seconds through PSM filtering. Winnow required 201.086 seconds for local feature recalculation and PSM filtering. Including candidate generation and intensity/iRT/RT preparation, with preparation time estimated from measured throughput, gave an estimated total of 371.599 seconds. Scaling by 1,138,277/9,777 candidates yielded serial estimates of approximately 13.5, 10.2 and 12.0 hours for π-NovoQC, NovoBoard and Winnow, respectively.

#### Analysis definitions for Fig. 6

In **Fig. 6a-c**, DB denotes database-search support, +DN denotes protein groups added by de novo evidence and Total denotes the inferred union. Shared DB and Shared DN represent shared peptide sequences quantified by the respective workflows. **Fig. 6c** uses positive observations; 721 zero Shared DN values are excluded. Abundances are not normalized between workflows. Boxplots in **Fig. 6c, e, f and i** use medians, interquartile ranges and whiskers extending to observations within 1.5 interquartile ranges; outlier markers are suppressed in c without removing positive observations from the statistics.

Among the 387,854 positive-intensity in-database peptides in **Fig. 6a, 130,853** overlapped database-search identifications and were supported by 1,304,321 de novo PSMs. The additional 257,001 peptides comprised 175,739 peptides supported by 904,511 PSMs mapping to DB-supported groups, and 81,262 peptides supported by 371,932 PSMs supporting 18,439 additional protein groups. Additional recovery refers to absence from database-search results, not absence from reference protein sequences.

For the top-abundance subset in **Fig. 6d** and **Supplementary Fig. 16, r**etaining Ensembl and UniProt entries produced 1,055 input protein identifiers: 898 Ensembl and 157 UniProt. Of these, 1,016 were supported by at least two peptides. Metascape mapped 824 identifiers to 675 genes; 231 identifiers were not mapped. The common intensity floor, abundance medians and their comparison are reported in the main Results.

For functional enrichment in Fig. 6d, the 675 mapped genes from the highest-abundance 10% of de novo-supported protein groups were tested against the 30,238-gene Metascape background. The plot shows unadjusted hypergeometric P values.

In **Fig. 6e, q**uantms support requires agreement in peptide sequence and spectrum index among up to five candidates per spectrum before 1% FDR filtering. **Fig. 6f** classifies BLAST alignments using E ≤0.1, query length ≥6 and full-query alignment. Fully matched sequences have no nonidentical positions; partly matched sequences have 1-100 nonidentical positions and no fully matched assignment. Unmatched PSMs meet neither definition. Proportions are calculated separately for each raw file.

The isotope analysis in **Fig. 6g** examines experimental support for identifications recovered beyond database searching. The displayed estimates describe the complete QC-passing de novo PSM populations, which include database-identified and database-unidentified spectra. The isotope-missingness model and the distinct reference-error assumption and database-label threshold are specified in Methods.

For **Fig. 6h, s**hared protein bars count de novo variant protein IDs supported by shared SAP peptides; they are not intersections of protein sets. **Fig. 6i** uses DB cell-line/peptide combinations and de novo raw-file/peptide combinations. The input counts are 19,944 and 1,989, with 19,944 and 1,974 available abundance values, respectively. The boxplot statistics retain 284 DB zeros and seven de novo zeros, which are not visible on the logarithmic axis. Abundances retain their original, unnormalized units.

Fig. 6j counts SAP-supporting assignments, including repeated support for a peptide. The 46 assignments outside the displayed ±20-ppm window are not shown in that panel.

### Supplementary Tables

#### Supplementary Table 1: Reference-correct PSM retention by species for relaxed and combined scoring on the GraphNovo test set (Supplementary Fig. 1)

| Test species | Combined scoring with γ calibration | Relaxed scoring with γ calibration | Count difference |
|---|---|---|---|
| *A. thaliana* | 7125 | 6523 | 602 |
| *C. elegans* | 7845 | 7088 | 757 |
| *E. coli* | 9065 | 8466 | 599 |
| Total | 24035 | 22077 | 1958 |

Counts in **Supplementary Table 1** are species subsets of the pooled GraphNovo comparison in **Supplementary Fig. 1, u**sing the same fitted models and adjustment factors for all three species.

#### Supplementary Table 2: QC feature definitions and scalar inputs for DDA and DIA (Fig. 3b; Supplementary Note 2)

| Feature Name | Abbreviation | Feature Meaning | Calculation Method |
|---|---|---|---|
| Amino Acid Confidence Score | AA_Confidence | Fusion of top-K attention and intensity rank for each amino acid, scored after matching theoretical ions. Intermediate residue-level vector; its scalar summaries, not this vector as one scalar, enter the QC feature matrix. | For attention[i], check whether top-K peaks match theoretical b/y ions; if matched, score += fused(attn, rank) |
| Mean Amino Acid Confidence | AA_Confidence_ Mean | Mean confidence score of amino acids derived from the fusion of attention and intensity ranking | mean(aa_conf_list) |
| Variance of Amino Acid Confidence | AA_Confidence_Var | Degree of variation in the distribution of amino acid confidence scores | var(aa_conf_list) |
| Minimum Amino Acid Confidence | AA_Confidence_Min | The minimum value of the amino acid confidence | min(aa_conf_list) |
| Maximum Amino Acid Confidence | AA_Confidence_ Max | The maximum value of the amino acid confidence | max(aa_conf_list) |
| Number of Low-Confidence Amino Acids | Num_Low_ Confidence | Number of amino acids with confidence < 0.2 | sum(conf < 0.2) |
| Number of High-Confidence Amino Acids | Num_High_ Confidence | Number of amino acids with confidence > 0.8 | sum(conf > 0.8) |
| Anchor Amino Acid Ratio | Anchor_Ratio | Proportion of anchor amino acids supported by theoretical ions | (#anchor amino acids) / (peptide length) |
| Mean Attention Entropy | AA_Entropy_Mean | Mean information entropy of attention for each amino acid | mean(entropy_i) |
| Standard Deviation of Attention Entropy | AA_Entropy_Std | Variation of attention entropy values | std(entropy_i) |

| Feature Name | Abbreviation | Feature Meaning | Calculation Method |
|---|---|---|---|
| Minimum Attention Entropy | AA_Entropy_Min | Minimum value of information entropy | min(entropy_i) |
| Maximum Attention Entropy | AA_Entropy_Max | Maximum value of information entropy | max(entropy_i) |
| Mean Attention Concentration | AA_Concentration_Mean | Mean sum of top-K attentions | mean(topk_attention_sum) |
| Standard Deviation of Attention Concentration | AA_Concentration_Std | Variance of the sum of top-K attentions | std(topk_attention_sum) |
| Minimum Attention Concentration | AA_Concentration_Min | Minimum concentration of top-K attentions | min(topk_attention_sum) |
| Maximum Attention Concentration | AA_Concentration_Max | Maximum concentration of top-K attentions | max(topk_attention_sum) |
| Mean Attention Offset | AA_Offset_Mean | Mean difference between attention peak positions and theoretical fragment positions | mean(abs(mz_i - theor_mz_i)) |
| Standard Deviation of Attention Offset | AA_Offset_Std | Standard deviation of the above offsets | std(abs(mz_i - theor_mz_i)) |
| Minimum Attention Offset | AA_Offset_Min | Minimum offset value | min(abs(mz_i - theor_mz_i)) |
| Maximum Attention Offset | AA_Offset_Max | Maximum offset value | max(abs(mz_i - theor_mz_i)) |
| Mean Center Drift | AA_Center_Drift_Mean | Mean difference between attention center positions and theoretical fragment positions | mean(abs(center - theor_mz)) |
| Standard Deviation of Center Drift | AA_Center_Drift_Std | Standard deviation of the above drifts | std(abs(center - theor_mz)) |
| Minimum Center Drift | AA_Center_Drift_Min | Minimum center offset value | min(abs(center - theor_mz)) |
| Maximum Center Drift | AA_Center_Drift_Max | Maximum center offset value | max(abs(center - theor_mz)) |
| Mean Attention Span | AA_Span_Mean | Mean span of m/z range for peaks corresponding to top-K attention | mean(max - min of m/z) |
| Standard Deviation of Attention Span | AA_Span_Std | Variation in attention m/z span | std(m/z span) |
| Minimum Attention Span | AA_Span_Min | Minimum span | min(m/z span) |
| Maximum Attention Span | AA_Span_Max | Maximum span | max(m/z span) |
| Global QC Score | Global_QC_Score | Average of explained intensity and fragment coverage | (explained_intensity + fragment_coverage) / 2; if the precursor ion doesn’t match, the score is multiplied by 0.1 |
| Theoretical Fragment Coverage | Fragment_Coverage | # of matched theoretical b/y ions divided by total theoretical ions | #matched / #total |

| Feature Name | Abbreviation | Feature Meaning | Calculation Method |
|---|---|---|---|
| Explained Intensity Ratio | Explained_Intensity | Proportion of peak intensity explained by theoretical ions relative to the total spectrum | sum(intensity[matched]) / total |
| Average Attention Similarity | Avg_Attention_ Similarity | Mean cosine similarity of attention vectors between adjacent amino acids | mean(cos_sim(ai, ai+1)) |
| Fragment Alignment Consistency | Fragment_ Alignment_ Consistency | Whether theoretical b/y ions cluster in high attention regions | Average attention for matched b/y ions |
| Absolute precursor m/z error (DDA) | precursor_error_ppm | Absolute difference between predicted and observed precursor m/z, normalised by predicted m/z | abs(predicted_mz − observed_mz) / predicted_mz × $10^6$; predicted_mz is computed from the candidate sequence and precursor charge |

In **Supplementary Table 2**, AA_Confidence is an intermediate residue-level vector; its scalar summaries enter the feature matrix. DDA uses 33 scalar inputs, including the absolute precursor m/z error defined above; DIA uses the corresponding 32 inputs without that precursor-error scalar.

#### Supplementary Table 3: Datasets, acquisition settings and database-search reference construction (Supplementary Note 3)

| Dataset | acquisition modes | Instruments | FDR of Database Search | Database Search Software |
|---|---|---|---|---|
| Nine-species Dataset (including *Homo sapiens*) | DDA | Thermo Scientific Q-Exactive | 1% | PEAKS DB |
| GraphNovo Dataset | DDA | Thermo Scientific Q-Exactive | 1% | SEQUEST [27] |
| MassIVE-KB Dataset | DDA | - | ~0% | MSGF+ [28] |
| DHP Dataset | DDA | Thermo Orbitrap Fusion or Thermo Orbitrap Lumos | 1% | MaxQuant / Quantms |
| Astral DDA and DIA datasets | DDA / DIA | Orbitrap Astral | 1% | DIA-NN 1.8.1 |
| Orbitrap DIA dataset | DIA | Orbitrap Fusion Lumos | 1% | MSFragger-DIA v21.1 |
| TimsTOF DDA dataset | DDA | TimsTOF | 1% | PEAKS-Online (version 12) |
| Dong-Ecoli-QE isotope-labeled dataset | DDA | Orbitrap Q-Exactive | 1% | Open-pFind (v1.0) |

Database-search FDR specifies reference-label selection. For the isotope analysis in **Fig. 6g, t**he upstream label cutoff q = 0.001 and the mixture model's reference-error assumption $\rho_0$ = 0.05 are separate parameters. De novo PSMs were selected at nominal 5% PSM FDR; the isotope model is defined in Methods.

#### Supplementary Table 4: External sequencing-model QC on the GraphNovo test set using per-entry calibrated FDR ≤5% (Fig. 4a,b)

| Sequencing model | Sequencer training source | Level | Total retained targets | Reference-correct count | Reference-incorrect count | Calibrated FDR estimate | Observed FDP |
|---|---|---|---|---|---|---|---|
| Casanovo | GraphNovo | PSM | 17,513 | 16,997 | 516 | 4.9997% | 2.9464% |
| Casanovo | MassIVE-KB | PSM | 18,266 | 17,352 | 914 | 4.9963% | 5.0038% |
| π-HelixNovo | GraphNovo | PSM | 22,230 | 21,117 | 1,113 | 4.9981% | 5.0067% |
| π-HelixNovo | MassIVE-KB | PSM | 25,342 | 24,138 | 1,204 | 4.9995% | 4.7510% |
| Casanovo | GraphNovo | Peptide | 11,623 | 11,235 | 388 | 4.9962% | 3.3382% |
| Casanovo | MassIVE-KB | Peptide | 11,817 | 11,688 | 129 | 4.9998% | 1.0916% |
| π-HelixNovo | GraphNovo | Peptide | 13,703 | 13,502 | 201 | 4.9998% | 1.4668% |
| π-HelixNovo | MassIVE-KB | Peptide | 16,064 | 15,733 | 331 | 4.9999% | 2.0605% |

The training source refers to the sequencing model; every row is evaluated on the pooled GraphNovo three-species test set. Counts and FDP use the same population selected at a per-entry adjusted FDR estimate ≤5%. **Fig. 4b** uses the specified tolerance around the 5% crossing, so its retention endpoints follow a different selection rule.

#### Supplementary Table 5: Reference-correct Cascadia PSMs retained using native features alone or with auxiliary reference-PSM features in six Astral DIA raw files (Fig. 4h)

| Acquisition setting and replicate | Native Cascadia features | Native Cascadia features + GraphNovo reference features | Native Cascadia features + Astral reference features |
|---|---|---|---|
| 2.5 ms, replicate1 | 38 | 676 | 2761 |
| 2.5 ms, replicate2 | 0* | 374 | 2042 |
| 2.5 ms, replicate3 | 0* | 495 | 2619 |
| 3.5 ms, replicate1 | 28 | 622 | 1846 |
| 3.5 ms, replicate2 | 0* | 346 | 2525 |
| 3.5 ms, replicate3 | 21 | 597 | 2635 |
| Total | 87 | 3110 | 14428 |

Counts include Cascadia candidates only. All settings retain the native Cascadia candidate features. The GraphNovo and Astral auxiliary reference-PSM feature sets were extracted by π-xNovo-DDA and π-xNovo-DIA, respectively, and combined with the native set for QC analysis. Auxiliary references are excluded from the reported yield. An asterisk denotes a native-feature run with no finite adjustment factor.

#### Supplementary Table 6: Development and evaluation raw files for the DHP HCD method comparison (Fig. 5)

| Purpose | Trypsin | AspN | LysN | Total |
|---|---|---|---|---|
| Winnow training and validation | 7 | 7 | 8 | 22 |
| Method evaluation | 27 | 26 | 32 | 85 |

#### Supplementary Table 7: Three-method PSM comparison on the GraphNovo development population at nominal 5% FDR

| Method | Scored PSMs | Retained PSMs | Reference-correct | Reference-incorrect | FDP |
|---|---|---|---|---|---|
| π-NovoQC | 36,887 | 25,146 | 23,886 | 1,260 | 5.0107% |
| NovoBoard | 36,887 | 23,446 | 21,868 | 1,578 | 6.7304% |
| Winnow | 36,546 | 8,581 | 8,372 | 209 | 2.4356% |

#### Supplementary Table 8: Pooled PSM results for three methods across 85 DHP HCD evaluation raw files (Fig. 5a)

| Method | Scored PSMs | Retained PSMs | Reference-correct | Reference-incorrect | Pooled FDP |
|---|---|---|---|---|---|
| π-NovoQC | 1,138,263 | 567,258 | 558,393 | 8,865 | 1.5628% |
| NovoBoard | 1,138,277 | 162,507 | 147,936 | 14,571 | 8.9664% |
| Winnow | 1,065,018 | 467,876 | 445,663 | 22,213 | 4.7476% |

All methods filter within each evaluation raw file at nominal 5% PSM FDR. Pooled FDP divides the sum of reference-incorrect retained PSMs by the sum of retained target PSMs across the 85 files.

#### Supplementary Table 9: Per-file PSM FDP for three methods across 85 DHP HCD evaluation raw files (Fig. 5b)

| Method | Files with FDP above 5% | Maximum per-file FDP |
|---|---|---|
| π-NovoQC | 0 | 4.7686% |
| NovoBoard | 8 | 36.9443% |
| Winnow | 39 | 8.3289% |

#### Supplementary Table 10: Sequence-deduplicated PSM results for three methods on the GraphNovo development population

| Method | Unique sequences | Correct representative PSMs | Incorrect representative PSMs |
|---|---|---|---|
| π-NovoQC | 16,705 | 15,574 | 1,131 |
| NovoBoard | 15,740 | 14,371 | 1,369 |
| Winnow | 5,985 | 5,803 | 182 |

All methods use nominal 5% PSM FDR, followed by deduplication by full modified sequence. For each sequence, the highest-scoring PSM under that method provides the reference-correctness label. GraphNovo is also used for method development and parameter selection. These counts describe sequence representatives after PSM filtering.

#### Supplementary Table 11: Sequence-deduplicated PSM results for three methods across 85 DHP HCD evaluation raw files (Fig. 5d)

| Method | Sum of within-file unique peptide counts | Unique peptides after pooling all 85 files |
|---|---|---|
| π-NovoQC | 298,716 | 236,541 |
| NovoBoard | 60,788 | 46,856 |
| Winnow | 260,540 | 211,641 |

#### Supplementary Table 12: Native peptide-level QC by π-NovoQC across 85 DHP HCD evaluation raw files

| Method | Sum of per-file retained peptide counts | Reference-incorrect | Pooled FDP |
|---|---|---|---|
| π-NovoQC | 302,573 | 10,222 | 3.3784% |

**Supplementary Table 11** deduplicates full modified sequences after PSM filtering: within-file counts are summed across 85 evaluation raw files, while pooled counts deduplicate across all 85 files. **Supplementary Table 12** applies π-NovoQC peptide-stage QC and sums retained peptide counts across those files; two per-file peptide FDPs exceed 5%.

#### Supplementary Table 13: π-NovoQC PSM- and peptide-level QC by protease in the 85-file DHP HCD method-comparison evaluation

| Protease | QC level | Raw-file count | Total retained count | Maximum per-file FDP |
|---|---|---|---|---|
| AspN | PSM | 26 | 94,516 | 4.7686% |
| AspN | Peptide | 26 | 48,730 | 4.0873% |
| LysN | PSM | 32 | 124,478 | 3.9846% |
| LysN | Peptide | 32 | 60,745 | 5.9144% |
| Trypsin | PSM | 27 | 348,264 | 2.6490% |
| Trypsin | Peptide | 27 | 193,098 | 4.5318% |

The 85 evaluation raw files comprise 26 AspN, 27 trypsin and 32 LysN files (**Supplementary Table 6**; **Fig. 5**), with 22 development files kept separate. This method-comparison subset is drawn from the 2,458-file DHP QC application. The same 85 files are summarized at both QC levels; their numbers are not added across levels.

#### Supplementary Table 14: Three-method workflow runtimes for one DHP HCD raw file and extrapolations to the 85-file evaluation (Fig. 5f)

| Method | Representative-file workflow | Measurement type | Estimate for 85 raw files |
|---|---|---|---|
| π-NovoQC | 416.57 seconds | Measured | 13.5 hours |
| NovoBoard | 313.99 seconds | Measured | 10.2 hours |
| Winnow | 371.6 seconds | Measured local processing plus estimated upstream steps | 12.0 hours |

#### Supplementary Table 15: Measured and estimated processing-stage times for the representative DHP HCD raw file (Fig. 5f)

| Method | Stage | Seconds |
|---|---|---|
| π-NovoQC | Opening spectra | 5.64 |
| π-NovoQC | Model loading | 1.17 |
| π-NovoQC | Native prediction and feature generation | 245.01 |
| π-NovoQC | Counterpart-side feature generation | 142.13 |
| π-NovoQC | Scorer training and two-level QC | 19.96 |
| NovoBoard | Spectrum preparation | 25.18 |
| NovoBoard | Inference on original spectra | 102.92 |
| NovoBoard | Inference on perturbed spectra | 181.17 |
| NovoBoard | PSM filtering | 0.33 |
| Winnow | Full feature recalculation and PSM filtering (measured) | 201.09 |

The representative file is HCD fraction 10 of an AspN-digested GM12878 sample. The 85-file extrapolation scales by candidate PSM count (1,138,277/9,777). Winnow combines measured local processing with estimated upstream preparation; these are workflow estimates rather than measured whole-cohort cold-start times.

#### Supplementary Table 16: DHP raw files used for database re-search with quantms in the proteomic application (Fig. 6e)

| Cell line | Enzyme | Fragmentation | Number of raw files |
|---|---|---|---|
| K562 | Trypsin | HCD | 34 |
| GM12878 | AspN | HCD | 33 |
| GM12878 | AspN | ETD | 35 |
| HepG2 | LysN | HCD | 40 |
| HepG2 | LysN | ETD | 39 |

### Supplementary Figures

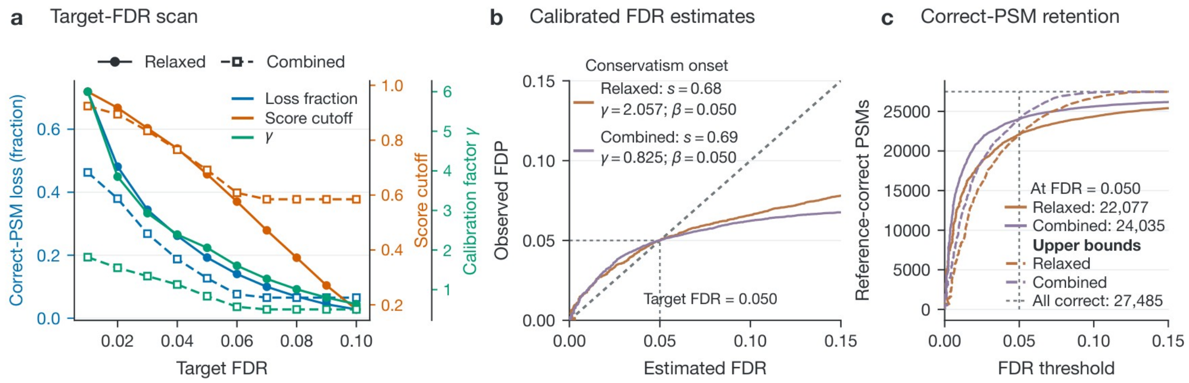


**Supplementary Fig. 1: Calibration and reference-correct PSM retention for relaxed and combined scoring on the GraphNovo test set.** a, Fraction of reference-correct PSMs lost at target FDRs from 0.01 to 0.10, calculated as (27,485 − retained reference-correct PSMs)/27,485. b, Calibrated FDR estimates versus observed FDP. c, Reference-correct PSM retention versus the FDR threshold. The analyzed population contains 37,321 target and 36,887 counterpart feature entries. Panel a uses the saved threshold-scan calibration; panels b and c use fixed adjustment factors for each scoring model. The separate weight experiments are shown in **Supplementary Figs. 7 and 8**.

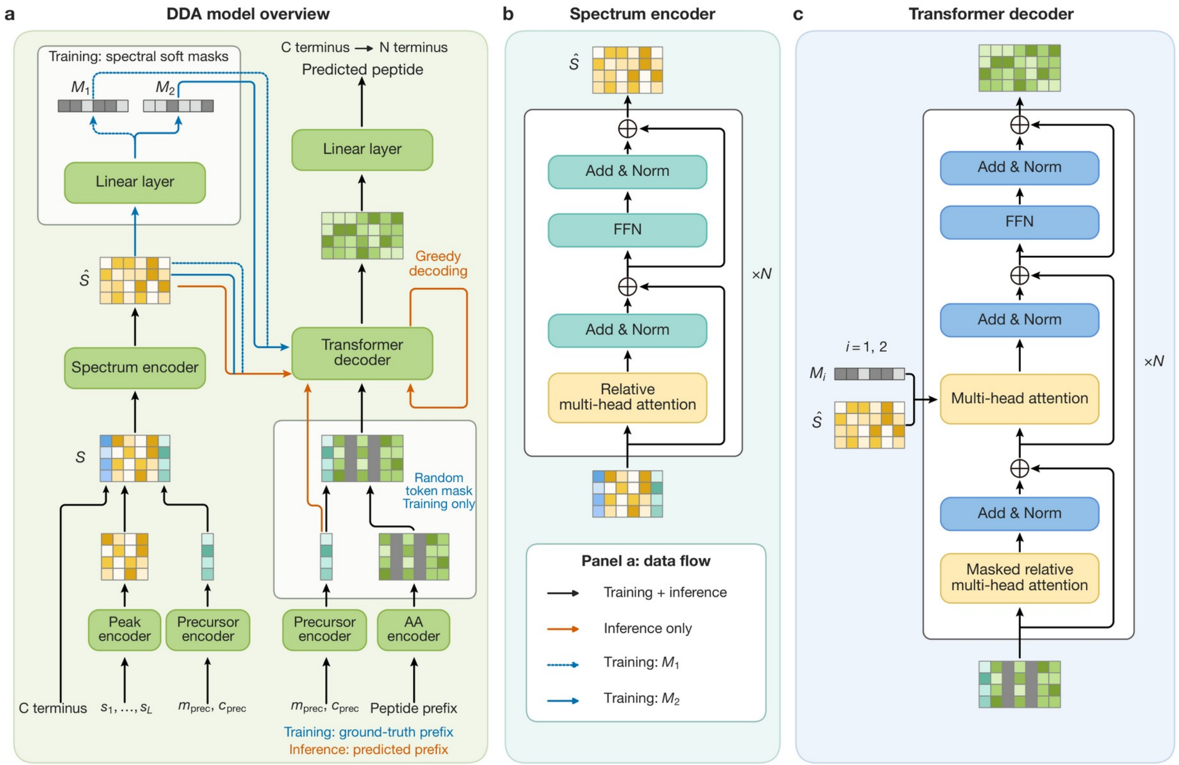


**Supplementary Fig. 2: Architecture of the π-xNovo DDA sequencing model.** a, Model overview. b, Spectrum encoder. c, Peptide decoder. Training uses complementary soft masks for the two spectrum views, token masking and teacher forcing. Inference uses autoregressive decoding with causal attention and disables the training perturbations. The sequencing-model framework is outlined in Methods; construction of paired QC features is described in Supplementary Note 2.

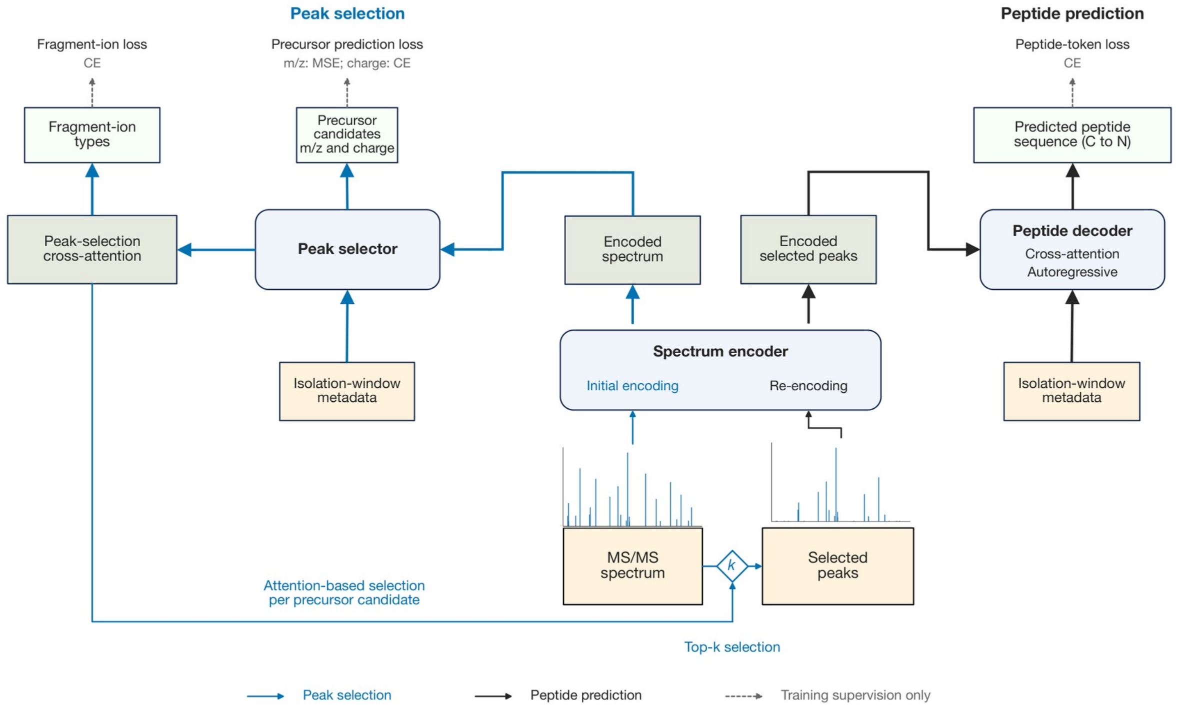


**Supplementary Fig. 3: Architecture of the π-xNovo DIA sequencing model.** The spectrum encoder, precursor-conditioned peak selector and peptide decoder form the sequencing workflow. The peak selector uses precursor information to select spectral peaks from the acquisition window, and the peptide decoder uses the selected 150-peak representation. The sequencing-model framework is outlined in Methods.

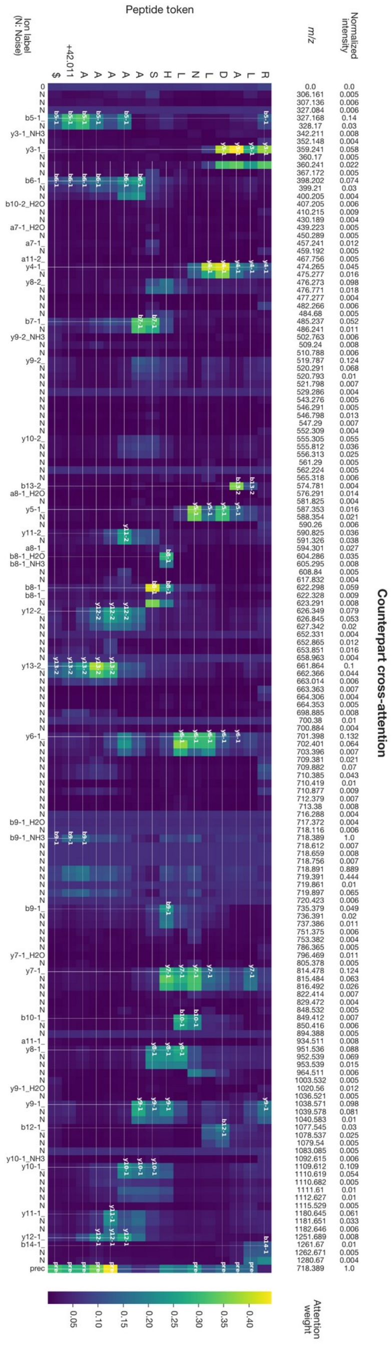


**Supplementary Fig. 4: Target and counterpart attention features for the same peptide prediction.** The attention map is aggregated across decoder layers and heads. Peptide queries include the stop token; spectrum positions include the precursor and the m/z = 0 token. N denotes peaks unmatched to theoretical fragment ions. Target and counterpart extraction windows are defined in Methods and **Supplementary Note 2**.

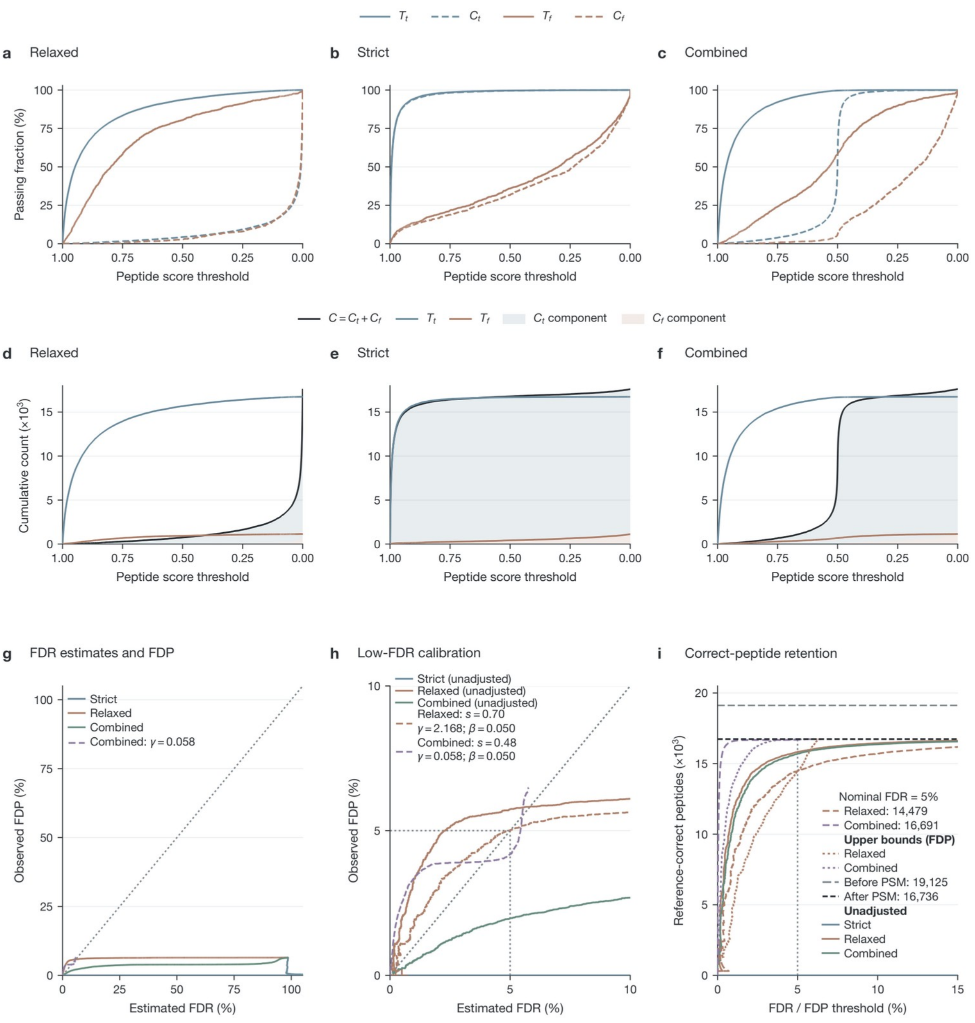

**Supplementary Fig. 5: Peptide-level scoring and calibration after PSM-level QC on the GraphNovo test set.** a-c, Fractions of entries passing each score threshold, normalized separately within each target or counterpart correctness group. d-f, Corresponding cumulative counts. g, Counting-based FDR estimates versus observed FDP. h, Calibration of relaxed and combined scoring near the selected FDR threshold. i, Reference-correct peptide retention versus the threshold. The shared peptide-stage population contains 35,540 target and counterpart feature entries, including 16,736 reference-correct targets. The curves use complete score ties. The calibrated endpoints in h and i use the saved near-threshold selection rule; the per-entry FDR ≤5% rule for the separate weight experiment is specified in **Supplementary Fig. 7**.

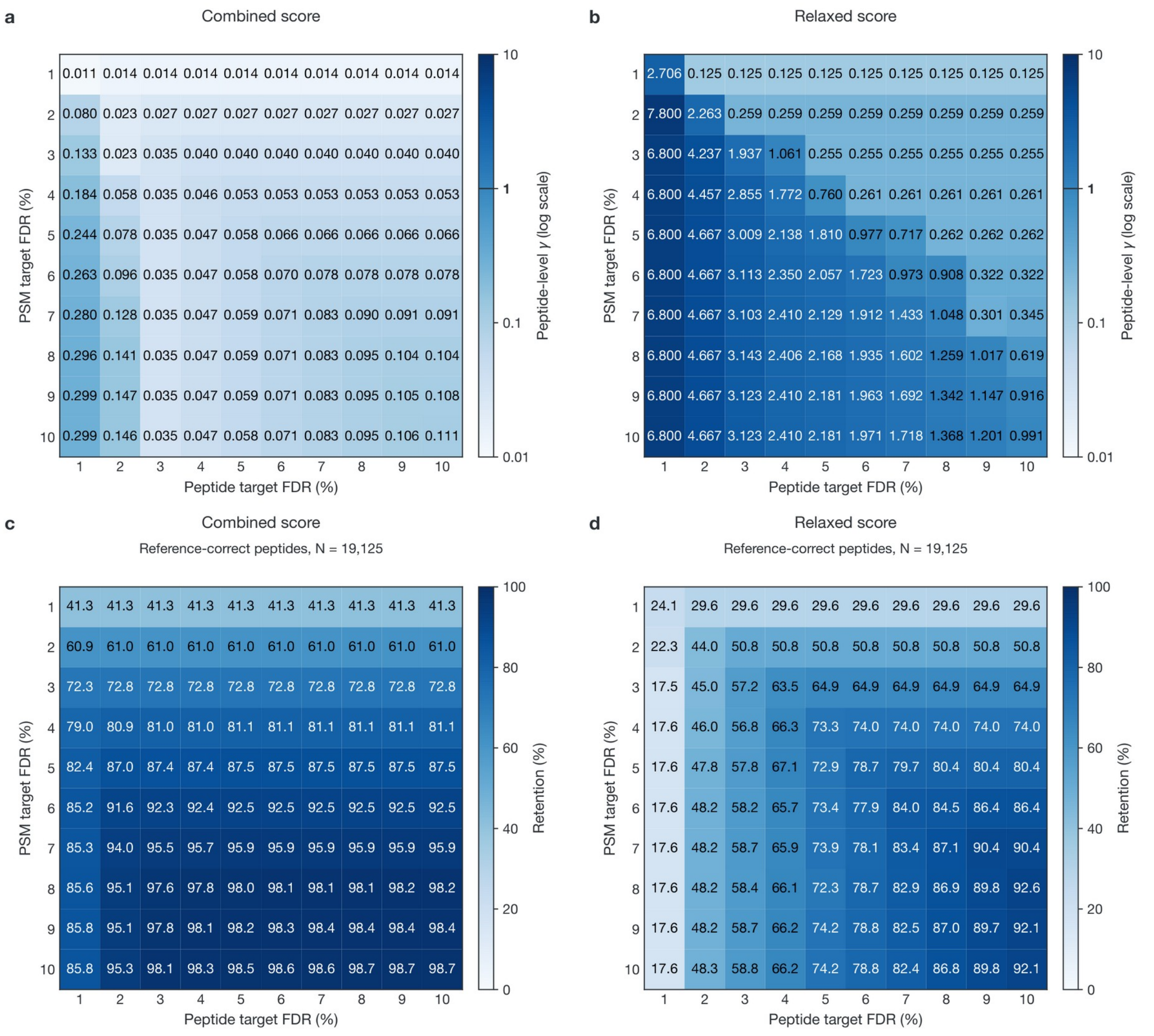


**Supplementary Fig. 6: Joint PSM- and peptide-level threshold scan on the GraphNovo test set.** a,b, Peptide-stage adjustment factors for combined and relaxed scoring, respectively. c,d, Corresponding reference-correct peptide retention. Both nominal FDR thresholds range from 1% to 10%. Retention is normalized to the common baseline of 19,125 reference-correct peptides before PSM filtering.

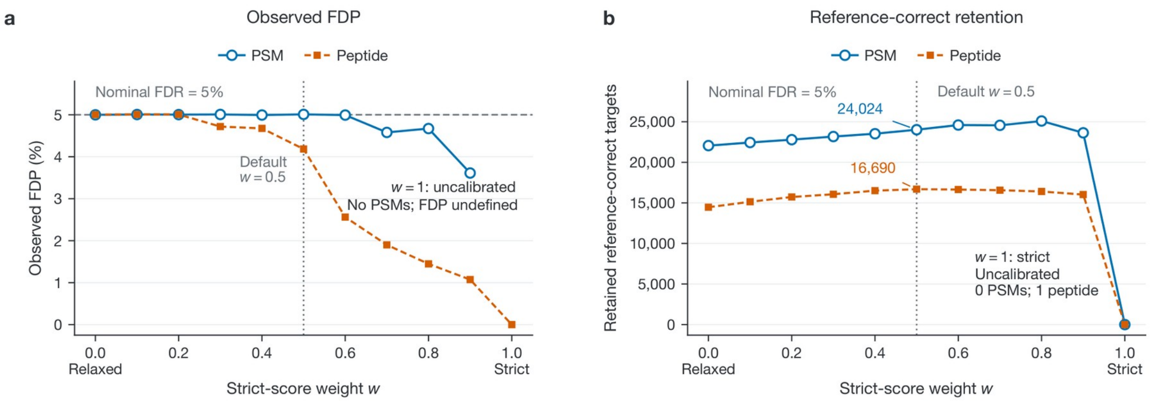


**Supplementary Fig. 7: Effect of the strict-model weight on final QC selection in the GraphNovo test set.** a, Observed FDP. b, Retained reference-correct targets. Eleven combined scores $w \times \text{strict} + (1 - w) \times \text{relaxed}$ use $w$ from 0 to 1 in steps of 0.1, with the fitted models held fixed. PSM-level analysis uses 37,321 target and 36,887 counterpart entries. Peptide-level analysis uses the same 35,540-entry population obtained after the default PSM selection for every weight. For $w < 1$, calibration uses the full reference-labeled population and selection requires a per-entry adjusted FDR estimate ≤5%. At $w = 1$, uncalibrated strict scoring retains no PSMs and one reference-correct peptide; the PSM FDP is undefined.

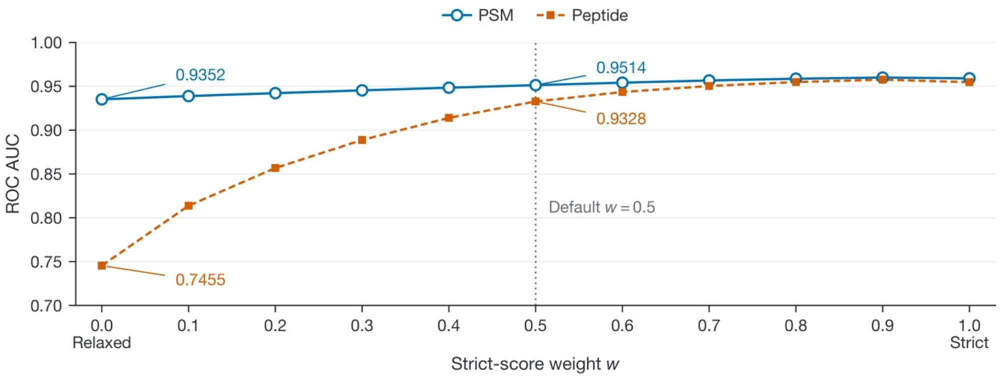


**Supplementary Fig. 8: Discrimination before FDR selection across strict-model weights on the GraphNovo test set.** ROC AUC distinguishes reference-correct from reference-incorrect targets using the fixed models and 11 weights defined in **Supplementary Fig. 7**. PSM-level AUC uses 37,321 targets; peptide-level AUC uses 17,896 targets from the fixed peptide-stage population. Peptide entries are not reconstructed for each weight.

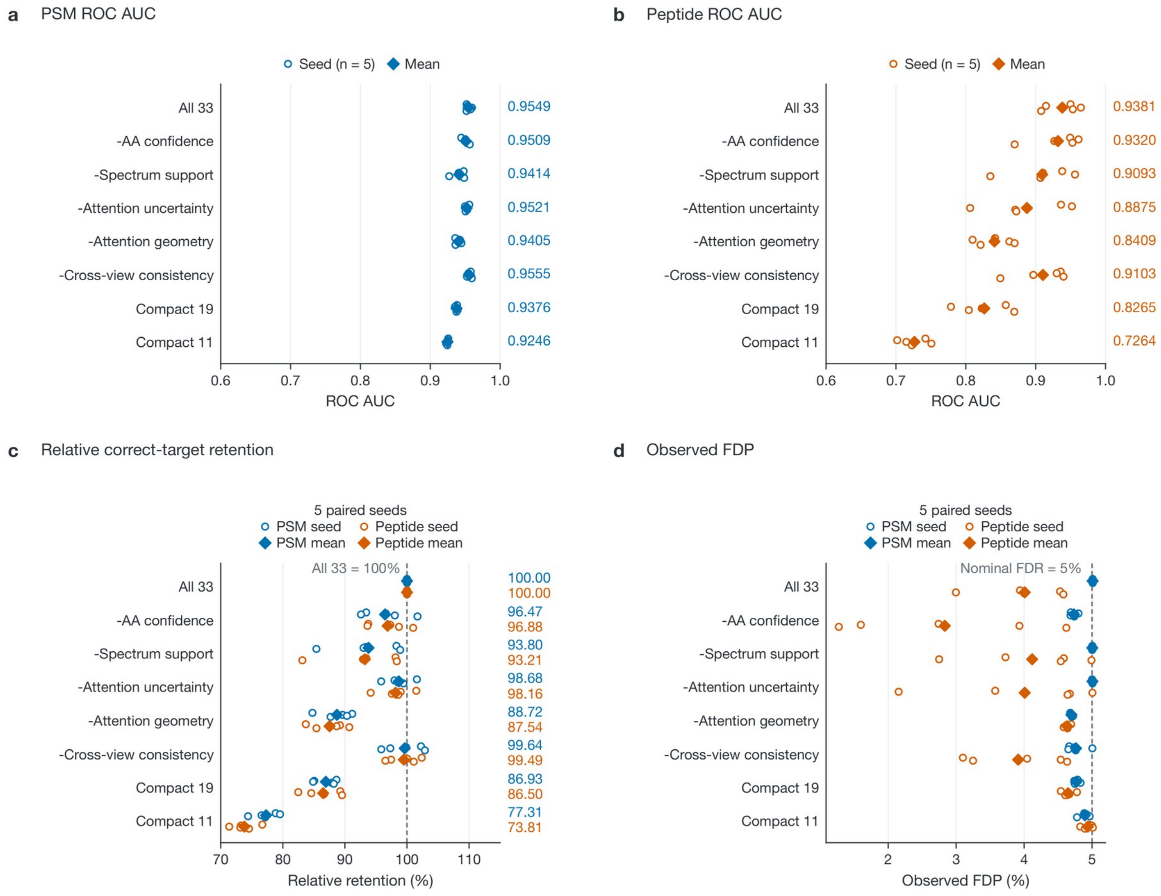


**Supplementary Fig. 9: DDA feature ablation on the GraphNovo test set.** Eight feature configurations are evaluated at the PSM and peptide levels using five paired random seeds (454, 2026, 3407, 6412 and 9103). a,b, ROC AUC before FDR selection at each QC level. c, Reference-correct retention relative to All 33 for the same seed. d, Observed FDP at a nominal FDR of 5%. Seed-level results and their arithmetic means are shown. The initial PSM population contains 74,208 target and counterpart feature entries; each configuration and seed constructs its own peptide population after PSM filtering. The five feature groups and all eight configurations, including Compact 11 and Compact 19, are defined in **Supplementary Note 2**.

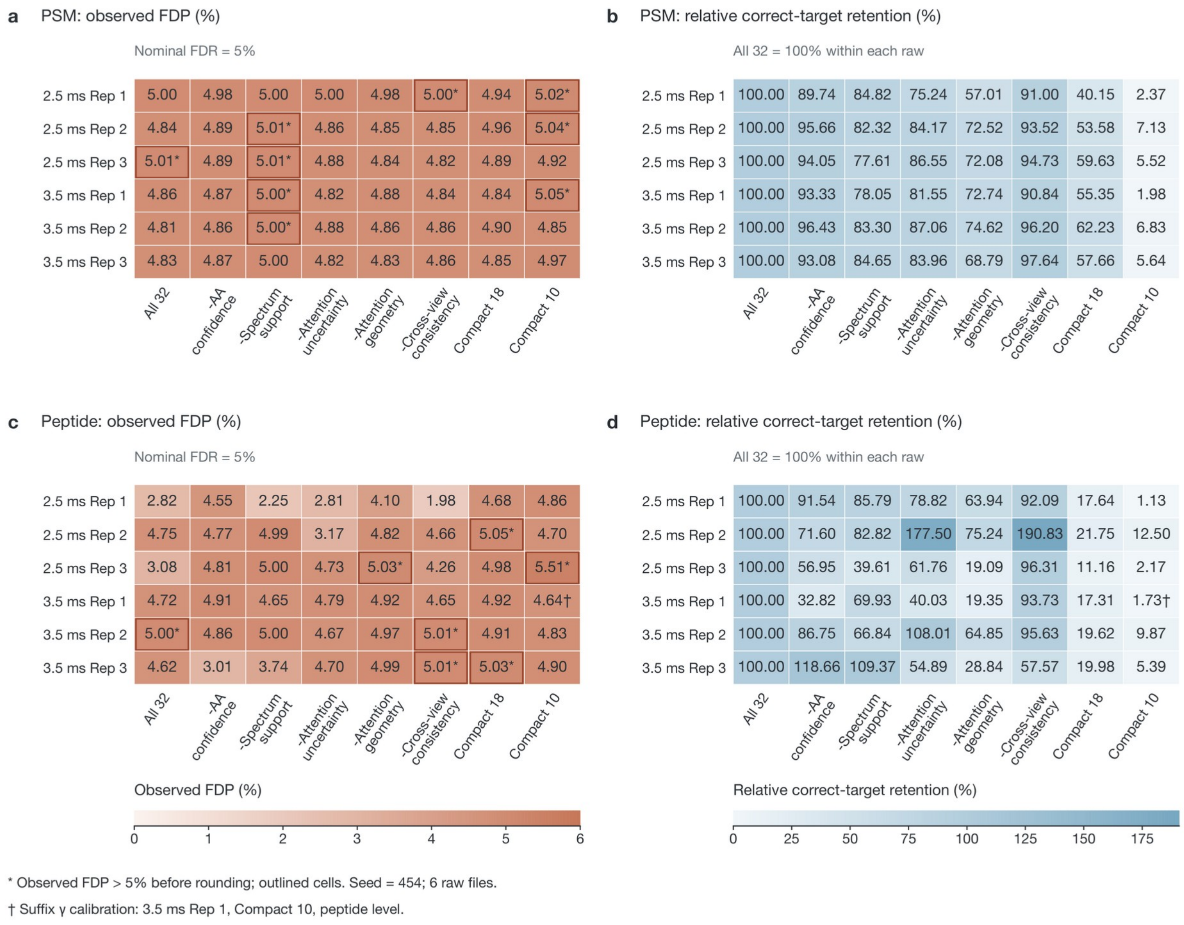


**Supplementary Fig. 10: DIA feature ablation across six Astral raw files.** Eight configurations use seed 454 and a nominal FDR of 5% at each QC level. a,c, Observed PSM and peptide FDP. b,d, Reference-correct retention relative to All 32 within the same raw file and QC level. Each configuration constructs its own peptide population after PSM filtering. The feature groups and All 32, Compact 10 and Compact 18 configurations are defined in **Supplementary Note 2**. An asterisk marks an FDP above 5% before rounding, including values displayed as 5.00*. The dagger marks Compact 10 for the 3.5-ms Rep 1 run: an empty local target-FDP neighborhood required suffix calibration on the fixed ranking (**Supplementary Note 1**, Theorem 3; $\gamma = 0.5271317829457365$), yielding 4.64% FDP and 1.73% relative retention.

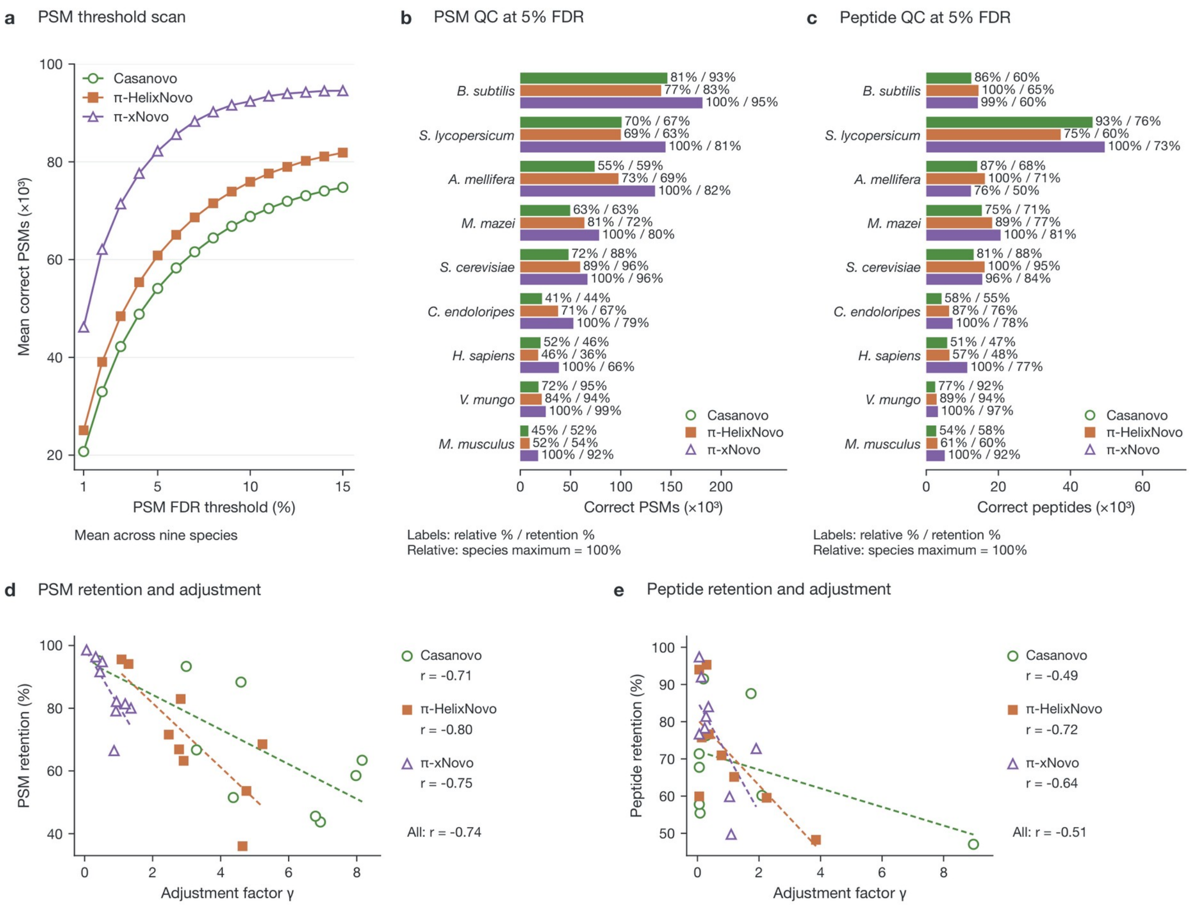


**Supplementary Fig. 11: Quality control of three sequencing models on the nine-species benchmark.** a, Mean retained reference-correct PSM counts across species versus target FDR. b,c, Reference-correct PSM and peptide retention at a nominal FDR of 5%. The two percentages express retention relative to the best model within the same species and QC level, and relative to all reference-correct predictions of that model, respectively. d,e, PSM and peptide retention versus the adjustment factor. Each point represents one model-species combination. Model-specific Pearson r uses nine species; All pools the 27 combinations. Sequencing models use leave-one-species-out training. Downstream QC models use an 8:2 reference split within each held-out species, followed by calibration and selection over the full candidate population (**Supplementary Note 3**).

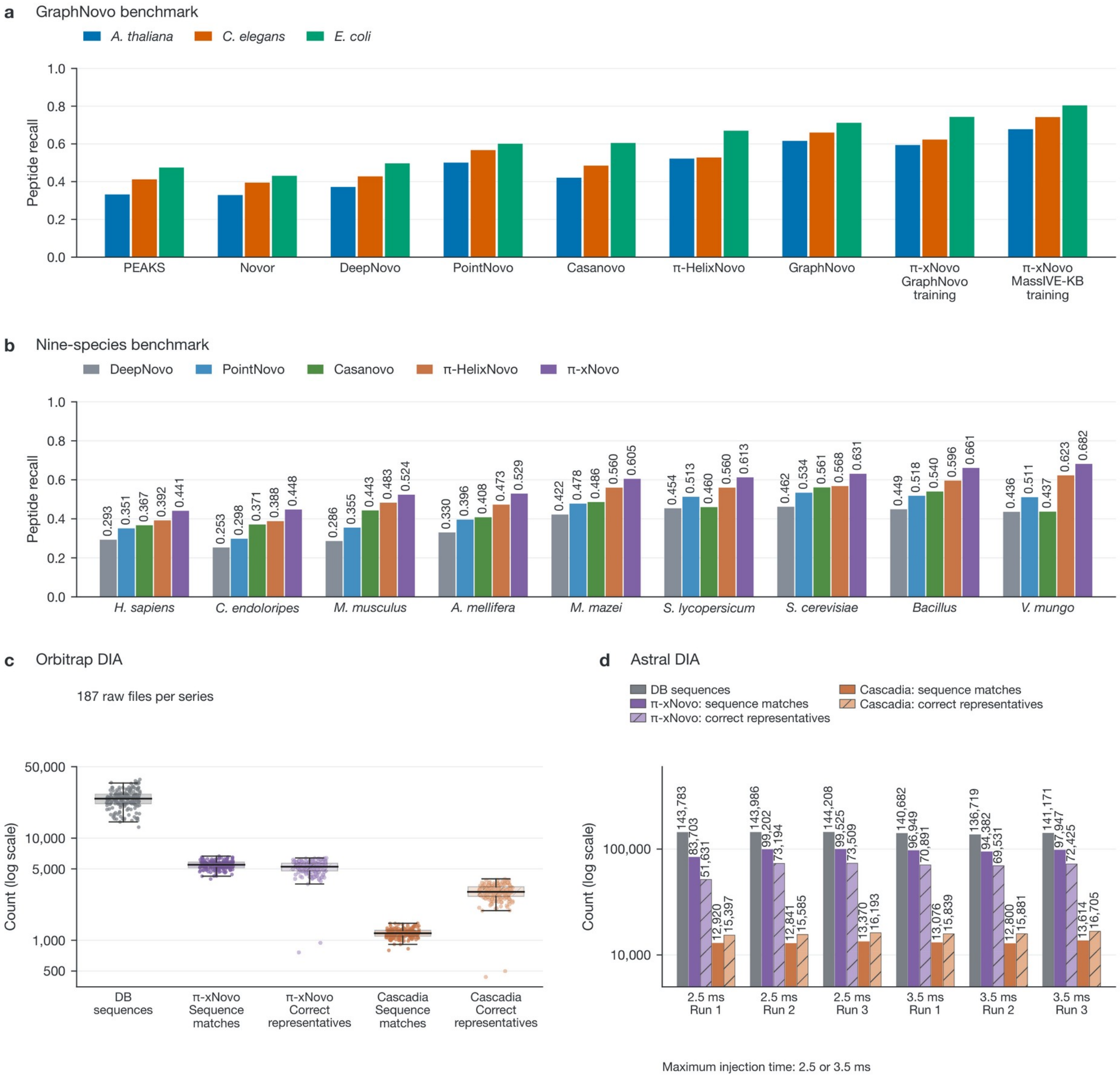


**Supplementary Fig. 12: Peptide-recall benchmarks and reference-supported DIA sequence recovery.** a, Peptide recall on the three-species GraphNovo test set for PEAKS [31], Novor [32], DeepNovo [11], PointNovo [33], Casanovo [34], π-HelixNovo [35], GraphNovo [12] and π-xNovo trained on GraphNovo or MassIVE-KB. b, Peptide recall on the nine-species benchmark. c, Sequence and representative-PSM counts for π-xNovo and Cascadia [21] across 187 Orbitrap DIA raw files; seven files without π-xNovo target features are excluded from all five series. Boxes span the interquartile range, center lines mark medians, and whiskers extend to observations within 1.5 interquartile ranges. d, Corresponding counts for six Astral DIA raw files. In c and d, DB sequences are distinct database-reference strings; sequence matches are distinct predicted strings present in the reference set. Correct representatives sum reference-correctness labels after retaining the first PSM for each predicted sequence in input order, without score sorting. No additional FDR filter is applied to these counts.

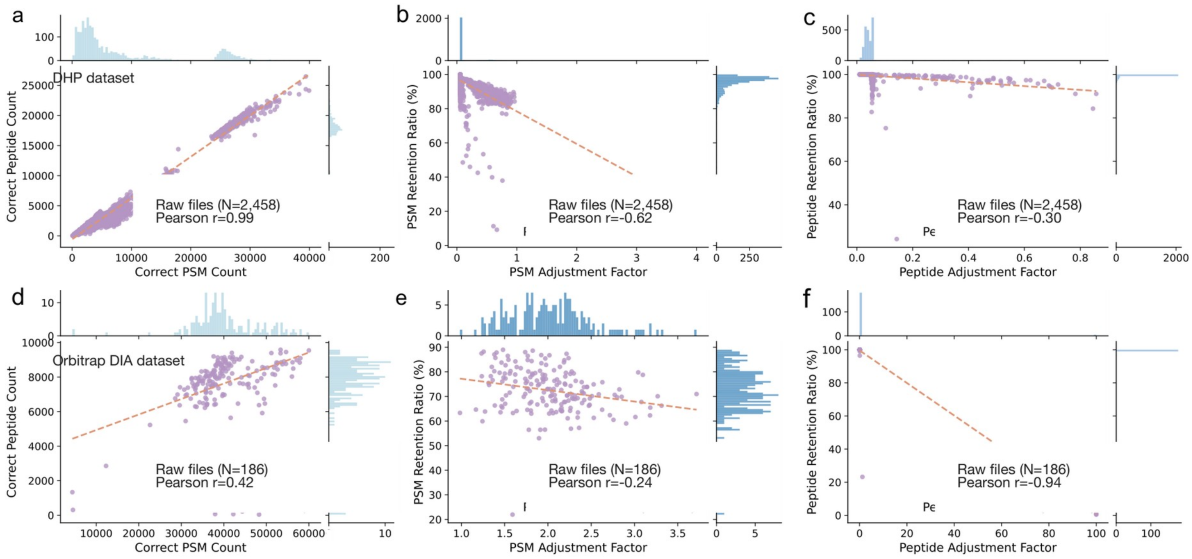


**Supplementary Fig. 13: PSM- and peptide-level QC in the large-scale DHP and Orbitrap DIA applications.** a-c, The full DHP QC application (2,458 raw files). d-f, The Orbitrap DIA QC application (186 raw files). a,d, Reference-correct peptide counts versus reference-correct PSM counts. b,e, PSM retention versus the adjustment factor. c,f, Peptide retention versus the adjustment factor. Each observation is one raw file; r is Pearson correlation.

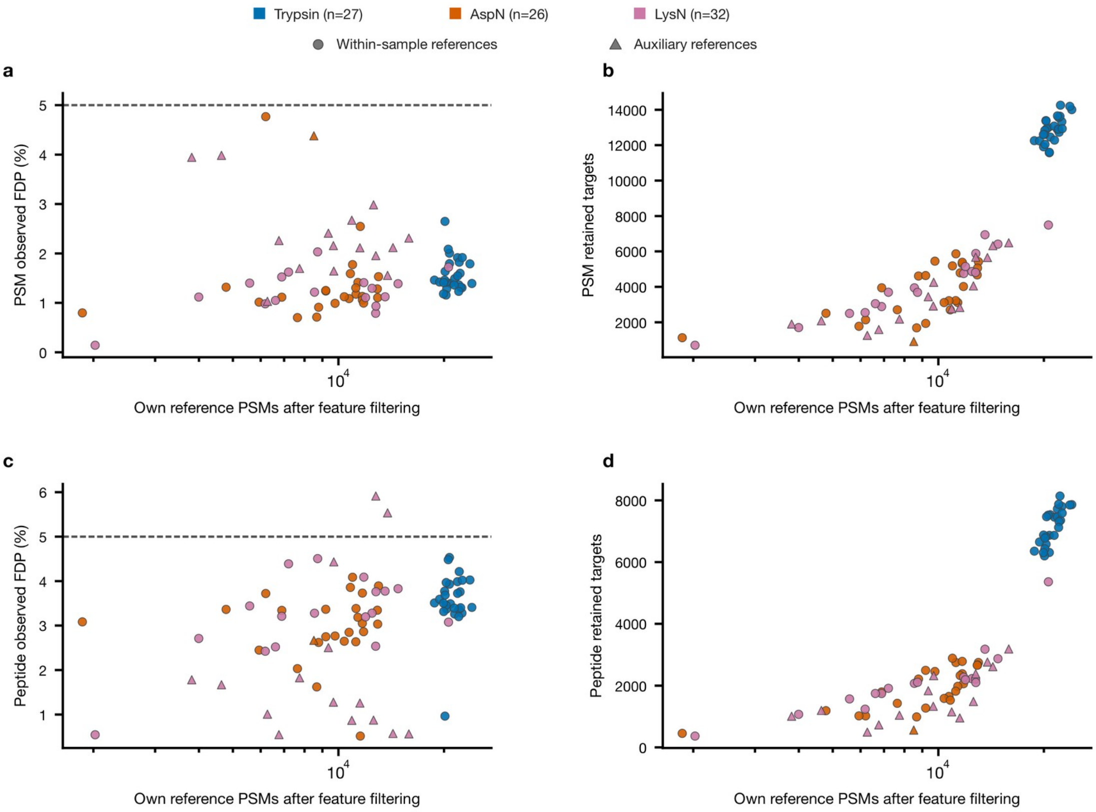


**Supplementary Fig. 14: Reference availability in the 85-file DHP HCD method-comparison evaluation.** The evaluation comprises 27 trypsin, 26 AspN and 32 LysN raw files, separate from the 22 development files (**Supplementary Table 6**; **Fig. 5**). Top, PSM-level results; bottom, peptide-level results. Left, observed FDP; right, retained target counts. Each point represents one evaluation raw file. The x axis counts that file's reference PSMs after feature filtering and excludes auxiliary references.

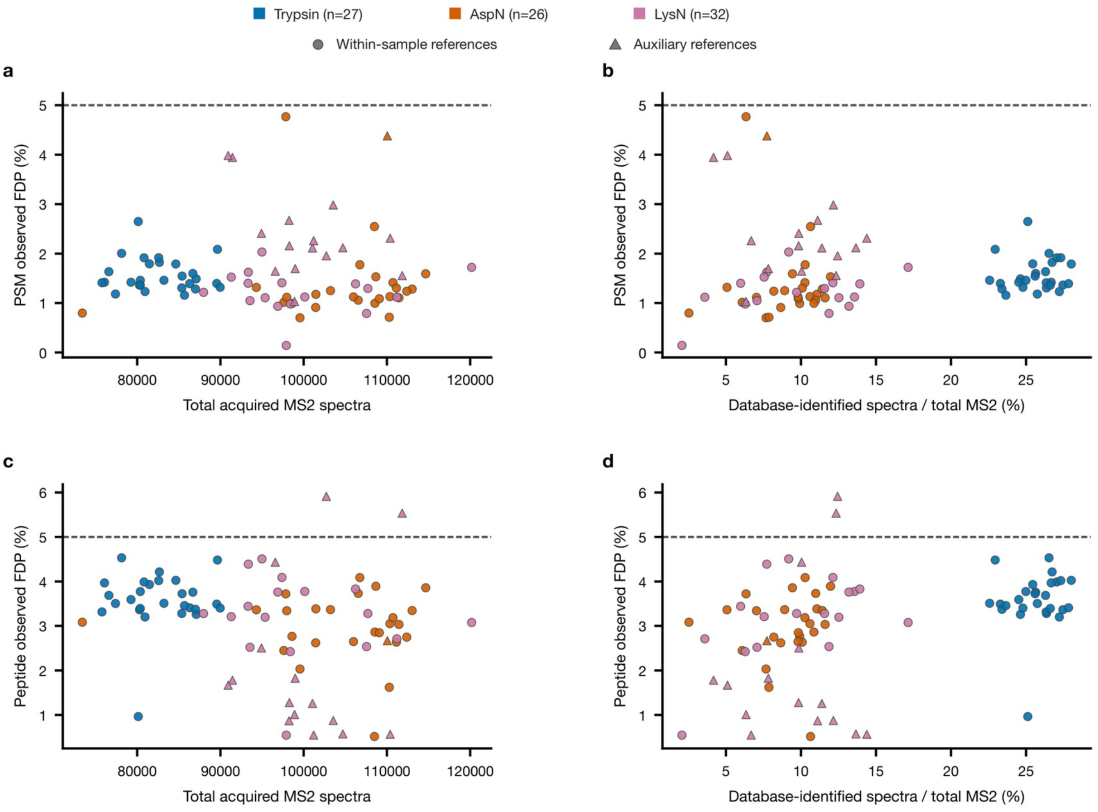


**Supplementary Fig. 15: QC performance versus spectrum coverage in the 85-file DHP HCD method-comparison evaluation.** The evaluation population is defined in **Supplementary Table 6** and **Supplementary Fig. 14**. Top, PSM-level observed FDP; bottom, peptide-level observed FDP. Each point represents one raw file. Left, total acquired MS2 spectra before database subtraction. Right, the fraction of those spectra with database-search identifications; auxiliary references are excluded from the numerator.

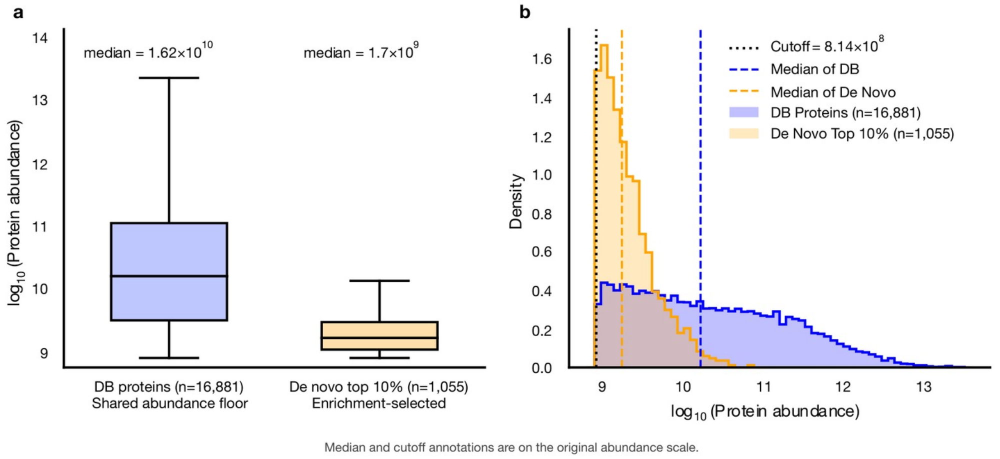


**Supplementary Fig. 16: Comparison of highly abundant protein intensities in the DHP quantitative application.** a, Log10 protein-intensity distributions. b, Density distributions of the same values. The top 10% of de novo protein intensities comprise 1,055 protein identifiers. Their minimum intensity defines a common floor of 814,048,865.1650392, above which 16,881 database-search observations are retained. Median annotations report intensities on the original scale. Quantification populations are described in **Supplementary Note 4** and **Fig. 6**.

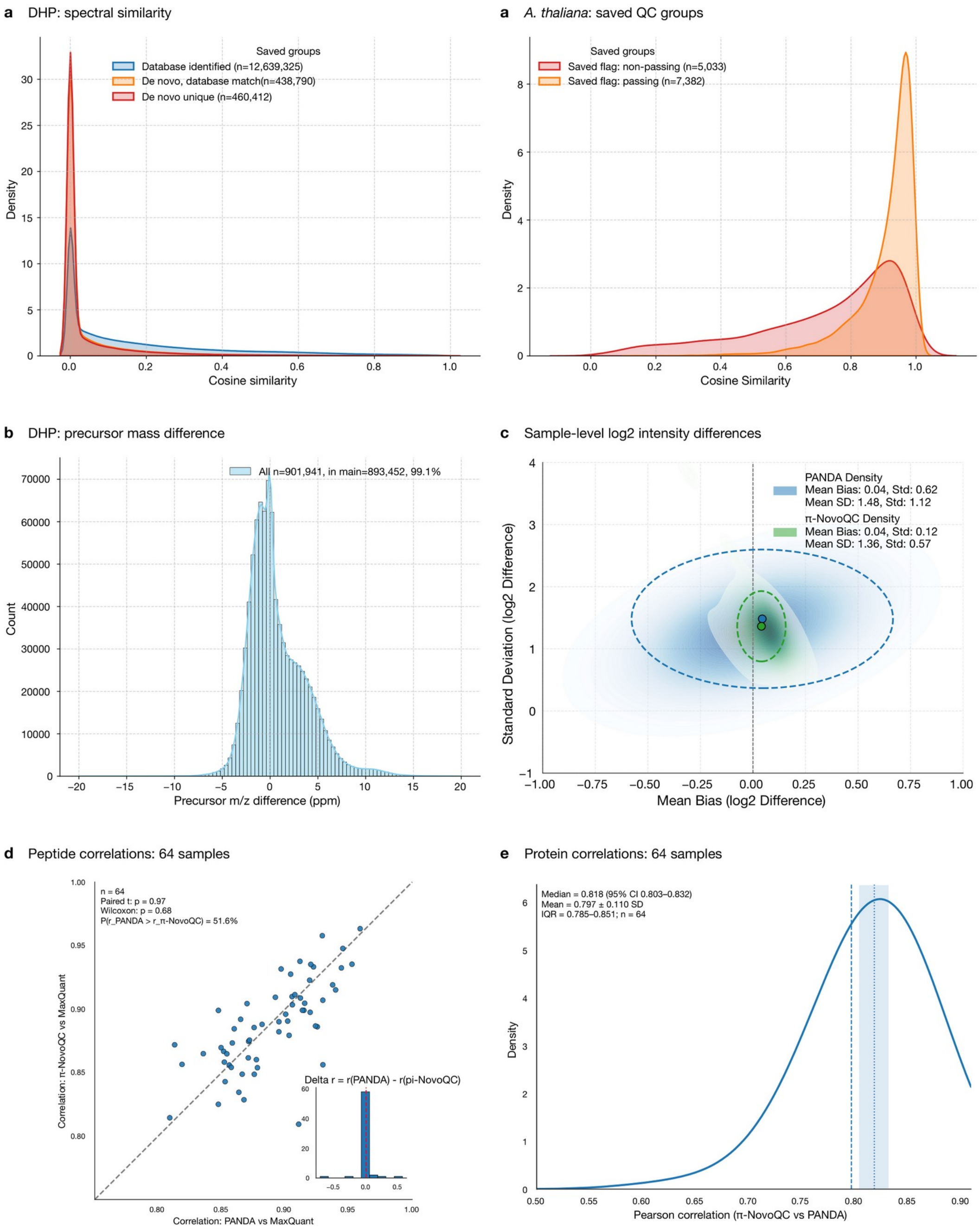


**Supplementary Fig. 17: Spectral support and quantitative agreement in the DHP application and GraphNovo test set.** a, Spectral cosine similarities for the three DHP identification groups (left), and for the *A. thaliana* test set partitioned by the stored QC flag (right; 7,382 passing and 5,033 non-passing observations). b, Precursor m/z differences for DHP peptide observations. c, Sample-level mean bias and standard deviation of log2 intensity differences from MaxQuant after median-offset normalization. Ellipse centers are the means of the sample statistics; widths and heights are twice the across-sample standard deviations of the respective coordinates. d, Peptide-quantification correlations with MaxQuant, with paired t and Wilcoxon tests across 64 sample summaries; the inset shows $\Delta r = r(\text{PANDA}) - r(\pi\text{-NovoQC})$. e, Protein-quantification correlations between π-NovoQC and PANDA across the same 64 summaries, with the reported median bootstrap interval. Quantification procedures are described in **Supplementary Note 4**.

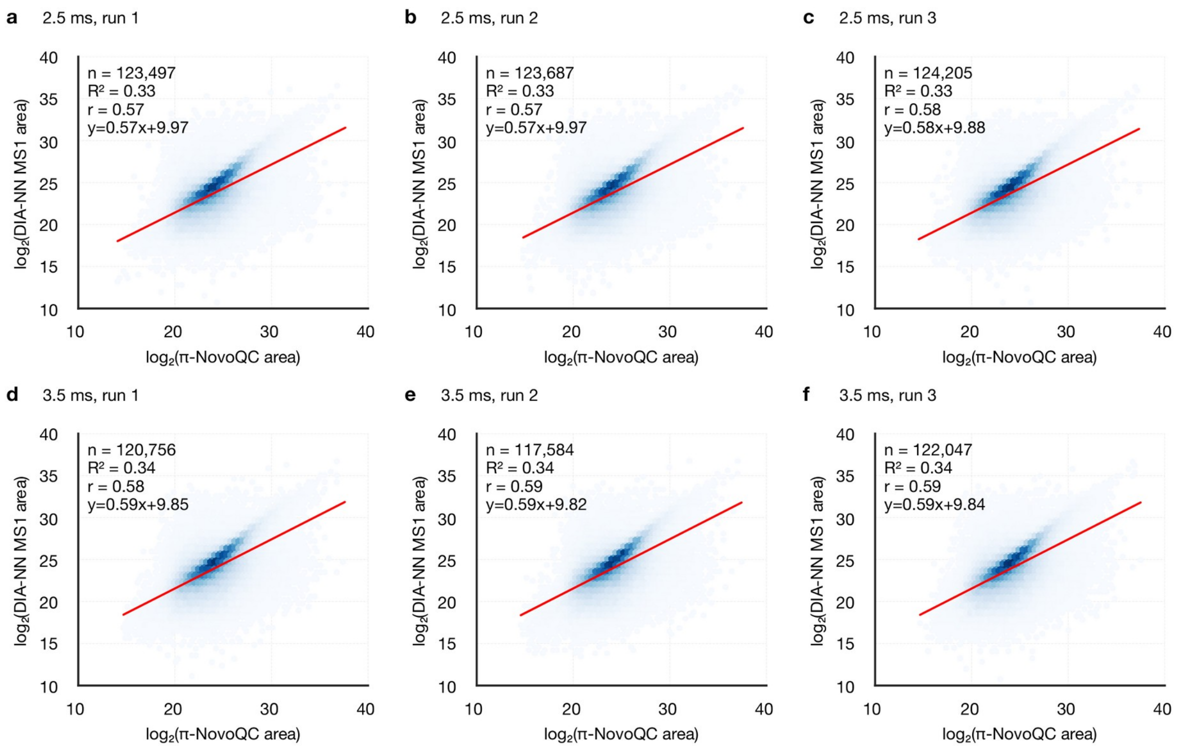


**Supplementary Fig. 18: Agreement of de novo and DIA-NN peptide quantification in six Astral DIA raw files.** a-c, The three 2.5-ms runs. d-f, The three 3.5-ms runs. Matched peptide intensities are compared on the log2 scale using π-NovoQC peak areas and DIA-NN MS1 areas. Each panel reports the matched-pair count, Pearson r and the linear fit.

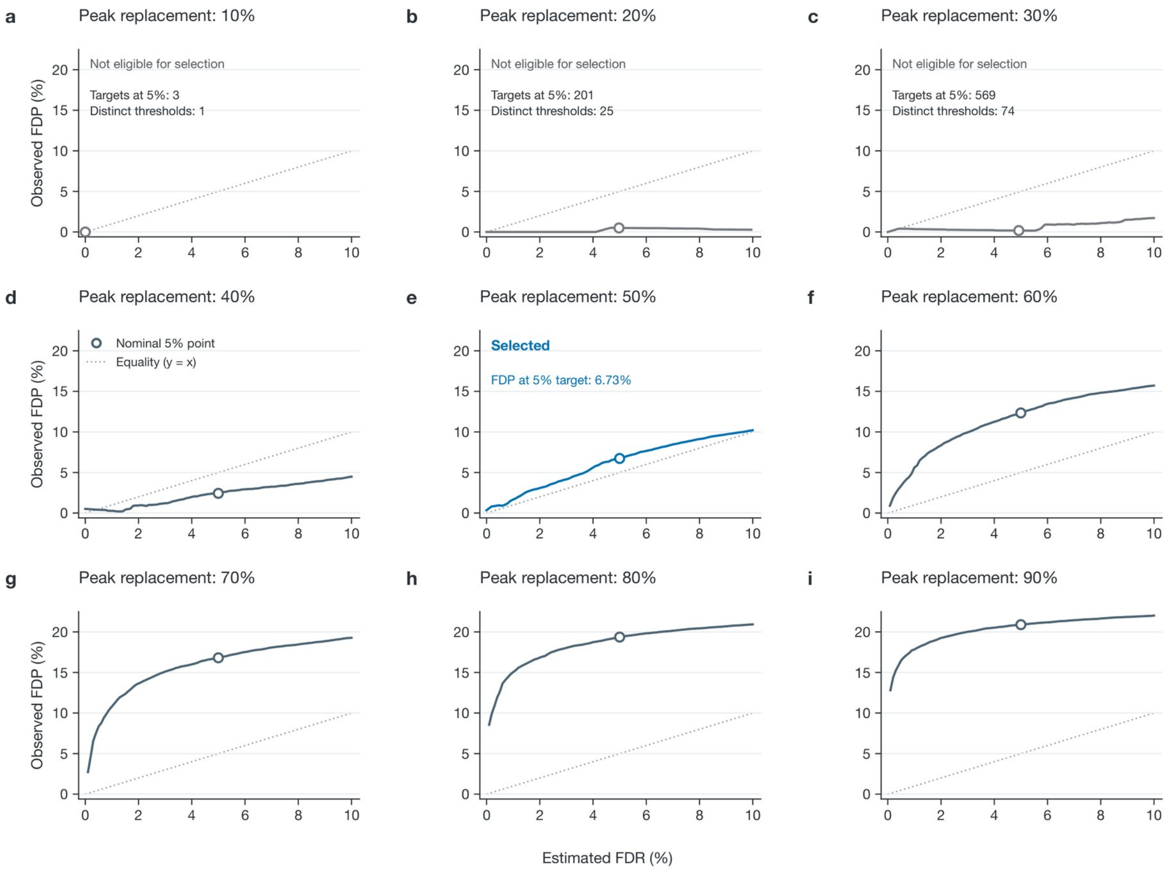


**Supplementary Fig. 19: NovoBoard fraction-parameter selection on the GraphNovo development population.** Nine fraction settings (0.1–0.9) use the same 36,887 target PSMs and seed 99. Each setting is evaluated at 100 nominal FDR thresholds from 0.1% to 10%. The x coordinate is the reported NovoBoard FDR estimate; observed FDP is the fraction of retained targets that disagree with the database reference. Repeated operating points overlap; for fraction = 0.1, all thresholds select the same three reference-correct PSMs. The selection rule in Methods chooses fraction = 0.5 on this development population. Its observed FDP at the nominal 5% setting is 6.7304% (**Supplementary Table 7**).
